# Fragment Binding to the Nsp3 Macrodomain of SARS-CoV-2 Identified Through Crystallographic Screening and Computational Docking

**DOI:** 10.1101/2020.11.24.393405

**Authors:** Marion Schuller, Galen J. Correy, Stefan Gahbauer, Daren Fearon, Taiasean Wu, Roberto Efraín Díaz, Iris D. Young, Luan Carvalho Martins, Dominique H. Smith, Ursula Schulze-Gahmen, Tristan W. Owens, Ishan Deshpande, Gregory E. Merz, Aye C. Thwin, Justin T. Biel, Jessica K. Peters, Michelle Moritz, Nadia Herrera, Huong T. Kratochvil, QCRG Structural Biology Consortium, Anthony Aimon, James M. Bennett, Jose Brandao Neto, Aina E. Cohen, Alexandre Dias, Alice Douangamath, Louise Dunnett, Oleg Fedorov, Matteo P. Ferla, Martin Fuchs, Tyler J. Gorrie-Stone, James M. Holton, Michael G. Johnson, Tobias Krojer, George Meigs, Ailsa J. Powell, Johannes Gregor Matthias Rack, Victor L Rangel, Silvia Russi, Rachael E. Skyner, Clyde A. Smith, Alexei S. Soares, Jennifer L. Wierman, Kang Zhu, Natalia Jura, Alan Ashworth, John Irwin, Michael C. Thompson, Jason E. Gestwicki, Frank von Delft, Brian K. Shoichet, James S. Fraser, Ivan Ahel

## Abstract

The SARS-CoV-2 macrodomain (Mac1) within the non-structural protein 3 (Nsp3) counteracts host-mediated antiviral ADP-ribosylation signalling. This enzyme is a promising antiviral target because catalytic mutations render viruses non-pathogenic. Here, we report a massive crystallographic screening and computational docking effort, identifying new chemical matter primarily targeting the active site of the macrodomain. Crystallographic screening of diverse fragment libraries resulted in 214 unique macrodomain-binding fragments, out of 2,683 screened. An additional 60 molecules were selected from docking over 20 million fragments, of which 20 were crystallographically confirmed. X-ray data collection to ultra-high resolution and at physiological temperature enabled assessment of the conformational heterogeneity around the active site. Several crystallographic and docking fragment hits were validated for solution binding using three biophysical techniques (DSF, HTRF, ITC). Overall, the 234 fragment structures presented explore a wide range of chemotypes and provide starting points for development of potent SARS-CoV-2 macrodomain inhibitors.

## INTRODUCTION

Macrodomains are conserved protein domains found in all kingdoms of life and certain viruses (*1*). Viral macrodomains have been found to recognize and remove host-derived ADP-ribosylation, a post-translational modification of host and pathogen proteins (*2*–*4*). The innate immune response involves signalling by ADP-ribosylation, which contributes to the suppression of viral replication (*4*–*9*). Upon viral infection, ADP-ribosylation is catalyzed by an interferon-induced subset of mammalian ADP-ribosyltransferases (ARTs), collectively termed ‘antiviral poly(ADP-ribosyl) polymerases’ (PARPs) (*4, 10*). These enzymes transfer the ADP-ribose (‘ADPr’) moiety of NAD^+^ onto target proteins (*4, 10*). For example, during coronavirus infection, PARP14 stimulates interleukin 4 (IL-4)-dependent transcription, which leads to the production of pro-inflammatory, antiviral cytokines (*11*). Viral macrodomains, which are found primarily in corona-, alpha-, rubi- and herpes-viruses, can counteract this host defense mechanism via their (ADP-ribosyl)hydrolase activity, contributing to the host-viral arms race for control of cell signalling (*12*).

Coronaviruses (CoVs) are important pathogens of livestock and humans. Three strains out of seven known to infect humans have caused major outbreaks within the last two decades: the severe acute respiratory syndrome (SARS) coronavirus, causing the SARS epidemic from 2002-2004, the Middle East respiratory syndrome (MERS) coronavirus, causing outbreaks in 2012, 2015 and 2018, and SARS-CoV-2, causing the current COVID-19 pandemic (*13, 14*). The coronaviral conserved macrodomain (called ‘Mac1’ here; also known as ‘S2-MacroD’ or ‘X domain’) is encoded as part of the non-structural protein 3 (Nsp3), a 200 kDa multi-domain protein (*15, 16*). While cell culture experiments suggest that SARS Mac1 is dispensable for viral replication in some cell lines (*6, 7, 17, 18*), animal studies have shown that its hydrolytic activity promotes immune evasion and that it is essential for viral replication and pathogenicity in the host (*8, 9*). The critical role of macrodomains is further supported by experiments using catalytic null mutations of the murine hepatitis virus (MHV), which render that virus essentially non-pathogenic (*6*–*8, 19*). Collectively, these findings lend strong support to the idea that SARS-CoV-2 Mac1 is a promising drug target for disrupting the viral life-cycle.

A barrier for macrodomain inhibitor discovery has been the lack of well-behaved inhibitors for this domain. Making matters worse, there are few biochemical assays suitable for screening for such inhibitors. Thus far, PDD00017273, an inhibitor of the poly(ADP-ribose)glycohydrolase (PARG), a macrodomain-type (ADP-ribosyl)hydrolase, remains the only well-characterized inhibitor with convincing on-target pharmacology and selectivity (*20*). The initial hit was discovered by a homogeneous time-resolved fluorescence (HTRF)-based assay which measures PARG activity, and renders the assay unsuitable for macrodomains that lack this activity (*21*). A selective allosteric inhibitor targeting PARP14 was identified in an AlphaScreen-based high-throughput screen (HTS) (*22*). While this inhibitor showed on-target activity in cells, its unique allosteric binding site is difficult to translate to other macrodomains. While potential Mac1 inhibitors have emerged with the advent of SARS-CoV-2 (*23*–*27*), their binding mechanisms and efficacy remain unclear. The lack of a biochemical assay specific for Mac1 has hindered their development. Furthermore, structures of the new inhibitors bound to Mac1 have not yet been reported, making optimization of initial hits, however promising, difficult.

To address the lack of chemical matter against Mac1, we turned to fragment-based ligand discovery using crystallography as a primary readout (**Fig. 1**). Fragment screens can efficiently address a large and relatively unbiased chemical space (*28*). Despite having generally weak overall binding affinity, fragments often have high ligand efficiency, and can provide templates for further chemical elaboration into lead-like molecules (*29, 30*). Crystallography can be used as a primary screening method for fragment discovery (*31*), and recent automation and processing software at synchrotron radiation sources has made this routinely possible at facilities like the XChem platform at Diamond Light Source (*32*–*37*). As part of Diamond’s contribution toward efforts to combat COVID-19, fragment screening expertise and infrastructure was made immediately available to any users working on SARS-CoV-2 targets (*38*). Similarly, synchrotron access for essential COVID-19-related research was also made available at the US Department of Energy light sources.

**Figure 1.**
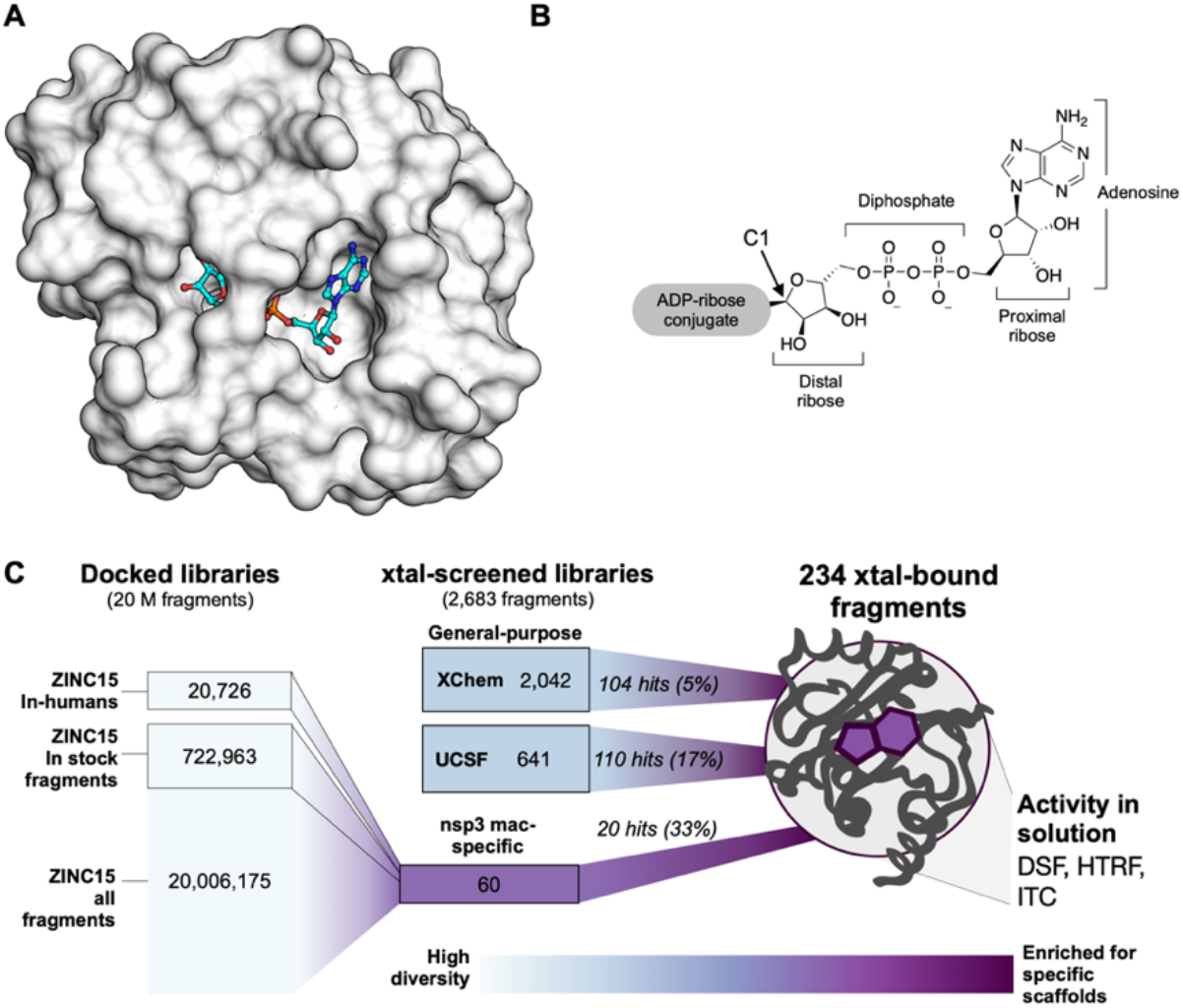
Overview of the fragment discovery approach for SARS-CoV-2 Nsp3 Mac1 presented in this study. **A)** Surface representation of Nsp3 Mac1 with ADP-ribose bound (cyan) in a deep and open binding cleft. **B)** Nsp3 Mac1 possesses ADP-ribosylhydrolase activity which removes ADP-ribosylation modifications attached to host and pathogen targets. ADP-ribose is conjugated through C1 of the distal ribose. **C)** Summary of the fragment discovery campaign presented in this work. Three fragment libraries were screened by crystallography: two general-purpose (XChem and UCSF), and a third bespoke library of 60 compounds, curated for Mac1 by molecular docking of over 20M fragments. Crystallographic studies identified 214 unique fragments binding to Mac1, while the molecular docking effort yielded in 20 crystallographically confirmed hits. Several crystallographic and docking fragments were validated by ITC, DSF, and an HTRF-based ADPr-peptide displacement assay.

Because crystallographic fragment screens can generate hits that bind anywhere on the protein surface, we wanted to supplement those screens with molecular docking intentionally targeting the active site. Docking has the additional benefit of exploring a much larger chemical space than an empirical fragment library. While an empirical library of ~1000-to-2000 fragments can represent a chemical space as large as, or larger, than that of a classic HTS library of several million compounds, exploration of chemotypes, including those that are well-suited to a particular target subsite, will inevitably be limited (*39*). Conversely, docking a much larger virtual library allows finer grained sampling around many chemotypes. A potential drawback of molecular docking is doubt about its ability to predict weakly-binding fragment geometries with high fidelity. While docking has identified potent ligands from libraries of lead-like molecules (250 to 350 amu) (*40*–*46*), such molecules offer more functional group handles for protein matching than do most fragments (150 to 250 amu), and docking is thought to struggle with the smaller, less complex, and geometrically more promiscuous fragments (*47, 48*). Thus, the pragmatism of this approach has been uncertain (*49, 50*).

Here, we present a combination of experimental crystallographic-based and computational docking-based fragment screens performed against Nsp3 Mac1 of SARS-CoV-2 (**Fig. 1**). Using X-ray crystallography, we screened fragment libraries of 2,683 compounds, yielding 214 unique fragment-bound Mac1 structures at atomic resolution. Docking of more than 20 million compounds prioritized 60 molecules for structure determination, yielding the structures of 20 additional compounds bound to Mac1. Additional X-ray data collection to ultra-high resolution and at physiological temperature illuminated the conformational heterogeneity in the Mac1 active site. We were able to confirm the binding of several fragments with differential scanning fluorimetry (DSF), isothermal titration calorimetry (ITC), and an ADPr-peptide displacement assay (HTRF), validating the activity of these molecules and providing a foundation for their optimization. The new fragments explore a wide range of chemotypes that interact with the catalytic site of the macrodomain. Together, these results create a roadmap for inhibitor development against Mac1, which may help to combat the pathogenicity of SARS-CoV-2.

## RESULTS

### Two crystal forms of Nsp3 Mac1 reveal differences in active site accessibility

We sought a crystal system that enabled consistent ligand soaking for fragment screening and for testing docking predictions. Six Mac1 crystal forms have previously been reported (**Supplementary data set 1**). Initially, we designed a construct based on PDB entry 6VXS (*25*). This construct has been reported to crystallize in P1, C2 and P21 with either 1 or 2 molecules in the asymmetric unit (ASU) (**Supplementary data set 1**). This construct crystallized reproducibly in C2 with microseeding and diffracted to a maximum resolution of 0.77 Å (**Supplementary data set 1**, **Fig. S1**, **Fig. S2** A). This high resolution data yielded electron density maps at true atomic resolution with abundant alternative conformations (**Fig. S1**). The electron density maps also revealed features that are rarely observed in macromolecular crystallography, such as explicit hydrogen atoms, and covalent bond density (**Fig. S1**). Although the active site appears accessible (**Fig. S3**), efforts to soak ADP-ribose into the crystals were unsuccessful. Additionally, soaking revealed that this crystal form suffers from inconsistent DMSO tolerance (**Fig. S2**), which is problematic for fragment soaking. In attempts to overcome this problem, we experimented with lysine methylation (*51*), which unfortunately increases the occlusion of the active site, and dehydration, which increased DMSO tolerance at the cost of non-isomorphism (**Fig. S2**).

In parallel, we designed a new Mac1 construct that crystallized in P4_3_ with two molecules in the ASU (**Supplementary data set 1**). This construct crystallized reproducibly with microseeding and diffracted to a maximum resolution of 0.85 Å (**Supplementary data set 1**). The sequence differences between the two constructs were slight (**Supplementary data set 1**), yet resulted in substantially different crystal packing (**Fig. S3**). Although the active site of protomer B was obstructed, the active site of protomer A was accessible (**Fig. S3**), and we were able to soak ADP-ribose into the crystals (**Fig. S4**B,C). This new structure also revealed a notable difference compared to previously reported Mac1-ADPr structures: the *α*-anomer of the terminal ribose was observed instead of the *β*-anomer (**Fig. S4**B-E). Despite this, alignment of ADP-ribose is excellent between all Mac1-ADPr structures (**Fig. S4**A). The DMSO tolerance of the P4_3_ crystals was excellent (**Fig. S2**). Accordingly, most of our fragment soaking work proceeded with this construct.

### Identifying new ligands for Nsp3 Mac1 using crystallographic fragment screening and docking

#### Characterization of experimental and virtual screening libraries

Crystal soaking screens at the XChem facility were performed with the P4_3_ crystals and a collection of fragment libraries (e.g. Diamond, SGC and iNEXT (DSI)-poised Library including 768 molecules (*34*) and the EU Open screen containing 969 molecules (*52*)) accumulating to 2,126 molecules (**Fig. S5**, **Supplementary data set 1**). Crystals were screened at the Diamond Light Source. At UCSF, a fragment library composed of Enamine’s Essential Fragment library with 320 compounds, augmented by an additional 91 molecules from an in-house library (UCSF_91), was screened against both the P4_3_ and C2 crystal forms at the Advanced Light Source (ALS), the Stanford Synchrotron Radiation Lightsource (SSRL) and the National Synchrotron Light Source-II (NSLS-II). On average, molecules across the X-Chem and UCSF collections had molecular weights of 192 + 47 amu, cLogP −1.8 - 3.8, 13 + 3 heavy atoms, and on average 2 rotatable bonds (**Fig. S5**).

Two fragment libraries were computationally docked against the structure of the macrodomain (PDB 6W02): a library of 722,963 fragments “in-stock” at commercial vendors, and the entire ZINC15 fragment library of 20,006,175 mainly make-on-demand fragments that have not been previously synthesized, but can readily be made, available predominantly from Enamine and Wuxi (*25*). Molecules from the ZINC15 fragment library had molecular weights ≤ 250 amu, cLogP ≤ 3.5, with an average of 4 rotatable bonds, and typically 4-19 heavy atoms (**Fig. S5**). In addition, an “in-human” library of 20,726 drugs also comprising investigational new drugs, and metabolites beyond the fragment chemical space, that have been tested in humans were included into the docking screen, with a view to potential repurposing opportunities. All three sets can be downloaded from ZINC15 (http://zinc15.docking.org) (*53*).

We investigated the fragment libraries for their diversity and their representation of chemotypes likely to bind at the adenine recognition site of the macrodomain (**Fig. S5**). Bemis-Murcko (BM) scaffold analysis revealed 179 unique scaffolds in the UCSF libraries, and 809 distinct scaffolds in the XChem fragment libraries. The in-stock fragment docking library contained 69,244 distinct scaffolds, while 803,333 scaffolds were present in the entire ZINC15 20M fragment collection. Taken together, the experimentally screened libraries contained roughly two compounds per BM scaffold, while the docking libraries contained approximately 10 fragments per scaffold, consistent with the expected higher granularity of the docking libraries afforded by their much larger size.

Since adenine-containing compounds are the only structurally characterized binders of Mac1 and fragment libraries are intended to cover a wide chemotype space, we assessed the prevalence of pyrimidines in the libraries. We observed pyrimidines in 12 of the 411 fragments in the UCSF libraries and 72 of the 2,126 XChem fragments (3.39% of the physically-screened fragments (**Fig. S5**). Pyrimidines were found in 41,531 of the 722,963 (5.74%) in-stock fragments and in 890,199 molecules of the 20,006,175 compound fragment library (4.44%). While the percentages of molecules carrying the pyrimidine substructure were similar between the physical and docked fragments, the absolute numbers in the latter sets were far higher. Aside from bearing a pyrimidine substructure, they were otherwise diverse: among the 890,199 pyrimidine-containing docking fragments, 60,919 distinct BM scaffolds were identified. Adenine was present in 5,457 fragments (582 different scaffolds). Furthermore, as ADP-ribose is negatively charged, anionic compounds were considered to exhibit favorable properties to bind to the macrodomain by targeting the diphosphate region. Fortuitously, a substantial fraction (35%) of the UCSF fragment libraries is anionic (**Fig. S5**).

#### Hit rates and Mac1 interaction sites of fragments

Across both crystal forms and facilities, we collected diffraction data for Mac1 crystals soaked with 2,850 fragments (**Fig. 2**). The diffraction characteristics of the P4_3_ crystals were excellent: the average resolution was 1.1 Å, and 98% of crystals diffracted beyond 1.35 Å (**Fig. 2**C,E). Although diffraction data was collected for 368 fragments soaked into the C2 crystals at UCSF, data pathologies meant that only 234 datasets could be analysed. The datasets collected from C2 crystals had a mean resolution of 1.4 Å and ranged from 1.0 to Å (**Fig. 2**A). In total, we identified 234 unique fragments binding to Mac1 using the PanDDA method (**Fig. 2**, **Supplementary data set 2**) (*54*). Of these, 221 were identified using P4_3_ crystals (hit rate of 8.8%) and 13 using C2 crystals (hit rate of 5.6%). 80% of the fragments were identified in the Mac1 active site, near to or overlapping with the regions occupied by the nucleotide (the adenosine site) or the phospho-ribose (the catalytic site) (**Fig. 2**G). Additional fragments were scattered across the surface of the enzyme, with an enrichment at a distal macrodomain-conserved pocket near lysine 90 (the ‘K90 site’, 14 fragments) and with many others stabilized by crystal contacts (**Fig. 2**B,D,F, **Fig. S6**). Coordinates, structure factors, and PanDDA electron density maps for all the fragments have been deposited in the Protein Data Bank (PDB) and are available through the Fragalysis webtool (https://fragalysis.diamond.ac.uk).

**Figure 2.**
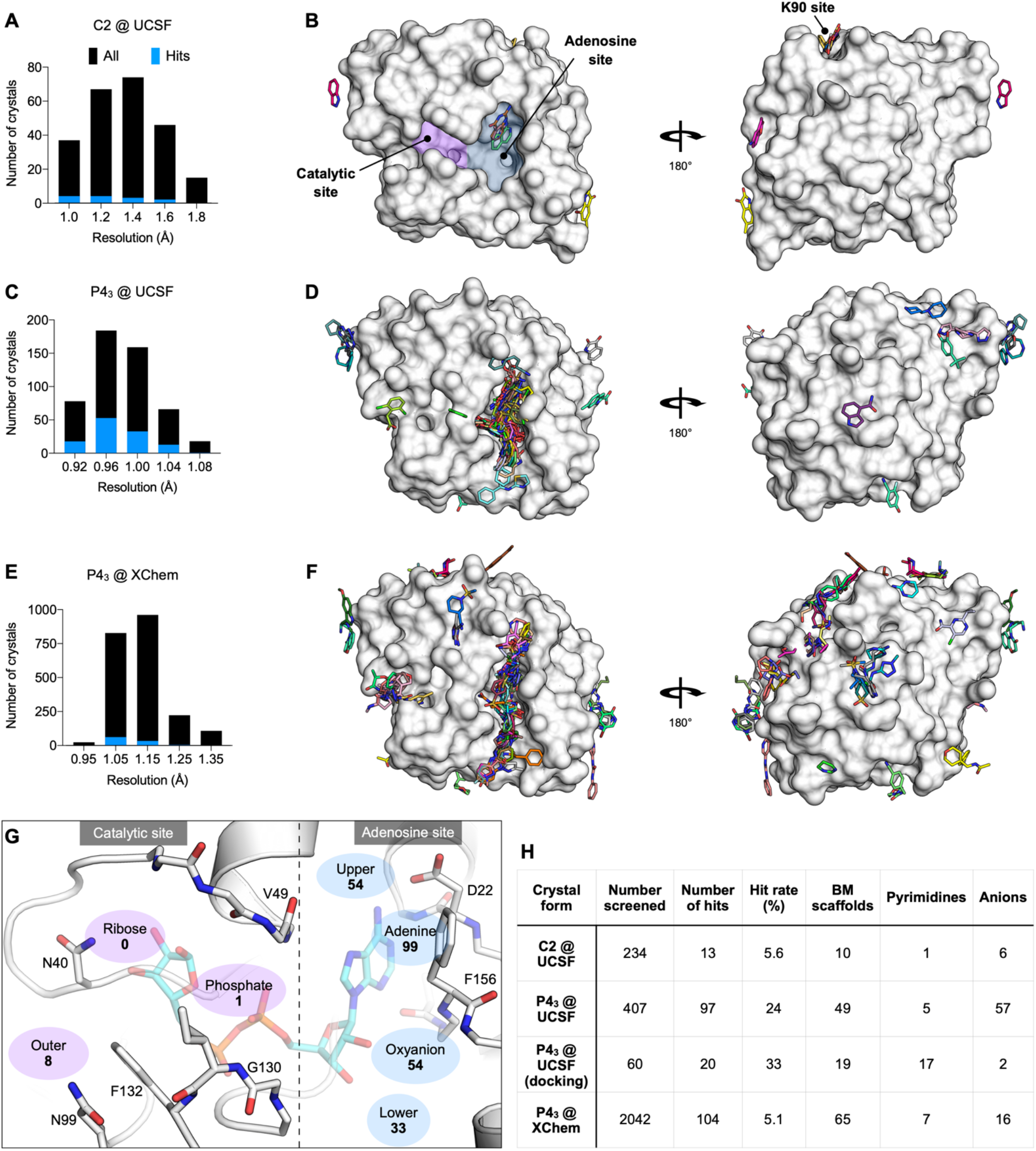
Crystallographic screening identified 234 fragments bound to Mac1. **A,C,E)** Histograms showing the resolution of the crystallographic fragment screening data. The resolution of datasets where fragments were identified are shown with blue bars. **B,D,F)** Surface representation of Mac1 with fragments shown as sticks. **G)** The Mac1 active site can be divided based on the interactions made with ADP-ribose. The ‘catalytic’ site recognizes the distal ribose and phosphate portion of the ADP-ribose, and harbours the catalytic residue Asn40 (*12*). The ‘adenosine’ site recognizes adenine and the proximal ribose. The number of fragments binding in each site is indicated. **H)** Summary of the fragments screened by X-ray crystallography, including the number of Bemis-Murcko (BM) scaffolds and anionic fragments identified as hits in each screen.

The unusually high hit rate for the adenosine site in the P4_3_ form with the Enamine Essential library (21%) was in contrast to the relatively low hit rate with this library with the C2 form (1.3%). Of the five pairs of fragments identified in both crystal forms, two pairs were identified in the adenine subside in both crystal forms, two in the adenine site in P4_3_ crystals but in the K90 site in C2 crystals, and the remaining pair bound to a surface site in the P4_3_ crystals and in the K90 site in the C2 crystals (**Supplementary data set 1**). Additional paired high quality datasets were available for 54 fragments that were bound within the P4_3_ crystals, but all showed no density for fragments in the C2 crystals (**Supplementary data set 1**). It is possible that competition for binding with the N-terminal residues may have contributed to the relatively low hit rate for the C2 form (**Fig. S3**).

#### Docking hits mimic the adenine recognition pattern

Docking the entire (20 million) ZINC15 fragment library, after careful calibration of docking parameters using different control calculations (see Methods) (*53, 55*), was completed in just under 5 hours of elapsed time on a 500 core cluster. The 20,006,175 fragments were sampled in over 4.4 trillion complexes. Top-ranked molecules were inspected for their ability to form hydrogen bonds similar to adenine (e.g. with the side chain of Asp22 and with the backbones of Ile23 and Phe156) while molecules with internal molecular strain or unsatisfied hydrogen bond donors were deprioritized. Ultimately, we selected 54 fragments from the entire ZINC15 fragment library screen, 9 of which were immediately available for purchase from Enamine and 33 of 45 make-on-demand molecules were successfully synthesized *de novo.* Furthermore, 8 fragments were purchased from the ZINC15 in-stock fragment library screen, and an additional 10 compounds were sourced based on the ‘in-human’ library docking (**Supplementary Table 1**).

Of the 60 molecules tested for complex formation by crystal soaking, 20 were observed with unambiguous electron density in complex with Mac1 (**Supplementary data set 1**). Here too, the crystals diffracted to exceptionally high resolution, between 0.94 and 1.01 Å. The predicted docking poses typically superposed well on the observed crystallographic results (Hungarian method root mean square deviations (*56*) ranging from 1-to-5 Å) and 19 out of the 20 docking hits bound to the adenosine sub-pocket of the Mac1, as targeted by docking (**Fig. 3 and Fig. S7**).

**Figure 3.**
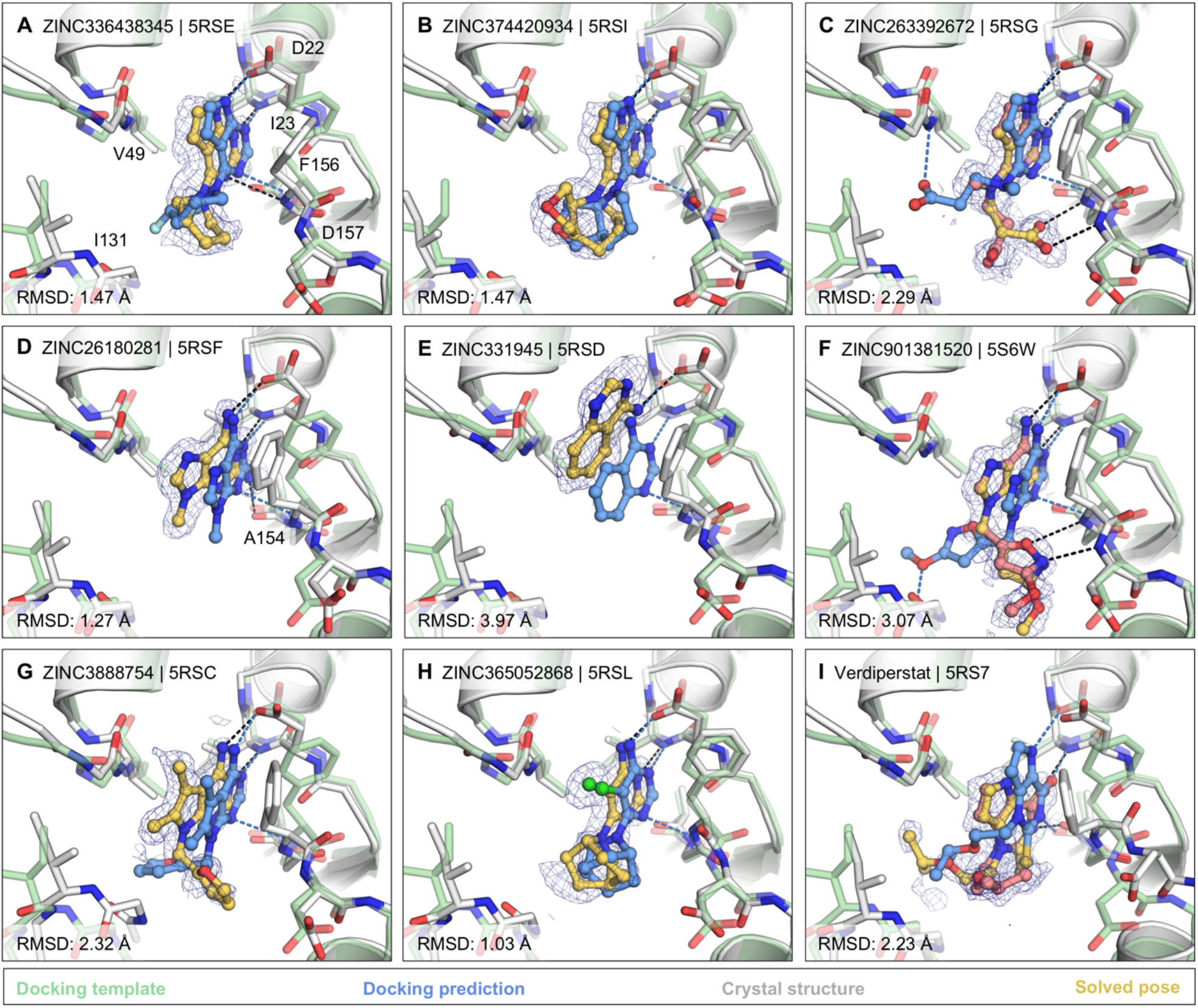
Docking hits confirmed by high-resolution crystal structures. The protein structure (PDB 6W02) (*25*), prepared for virtual screens is shown in green, predicted binding poses are shown in blue, the crystal protein structures are shown in grey, the solved fragment poses are shown in yellow, with alternative conformations shown in light pink. PanDDA event maps are shown as a blue mesh. Protein-ligand hydrogen bonds predicted by docking or observed in crystal structures are colored light blue or black, respectively. Hungarian RMSD values are presented between docked and crystallographically determined ligand poses (binding poses for additional docking hits are shown in **Fig. S7**).

The most commonly observed scaffold among the docking hits was 7H-pyrrolo(2,3-d)pyrimidine occupying the adenine-binding pocket for ADP-ribose (**Fig. 3**A-C). This ring system is typically hydrogen bonded with Asp22, Ile23 and Phe156. Fragments with this scaffold revealed high fidelity between the docking results and the high resolution structures (RMSD 1.5 - 2.3 Å). Different substituents can be attached to this headgroup e.g. piperidine, adding a hydrophobic segment to the scaffold (e.g. ZINC336438345 (PDB 5RSE)), occupying most of the adenosine binding pocket as shown in **Fig. 3**A,B and **Fig. S7**A,B. In addition to hydrophobic variations, ZINC263392672 (PDB 5RSG) attaches an anionic substituent to the pyrrolopyrimidine scaffold, offering additional hydrogen bonds within the binding pocket (**Fig. 3**C). Interestingly, while docking predicted the carboxylic acid of compound ZINC263392672 to insert into the phosphate binding tunnel, forming a hydrogen bond to Val49, the crystal structure instead revealed hydrogen bonds to the backbone amines of Phe156 and Asp157 which we defined as the ‘oxyanion site’ within the adenosine sub-pocket. Interactions with this backbone-defined oxyanion site were also observed for many other hits from both the docking and the crystallographically screened libraries (e.g. **Fig. 3**F, **Fig. S7** E).

For a set of smaller, mainly adenine-like docking hits, modeled to only occupy the adenine subpocket of the targeted adenosine binding site (**Fig. 3**D,E, **Fig. S7**C,D), the comparison between docked and experimental poses revealed deviations between 1.3 and 4 Å. Making these somewhat larger deviations harder to interpret was that for several fragments the crystallographically observed pose, e.g. ZINC331945 (RMSD 3.97 Å, **Fig. 3**E) and ZINC763250 (RMSD 3.78 Å, **Fig. S7**D), is partially stabilized by interactions with the symmetry mate (*vide infra*).

Another group of docking hits was selected as they mimicked the adenosine scaffold more closely (see **Fig. 3**F,G and **Fig. S7**I-L). For these, the ultra-high resolution of the crystal structures was crucial, revealing that for four of these, initially the wrong purine isomer had been inadvertently synthesized, with alkyl derivatives from the N3 rather than the intended N9 nitrogen corresponding to the alkylation of adenine in ADP-ribose (**Fig. S7**I-L). Characterization of the original compound samples by HPLC/MS and NMR confirmed that the delivered compounds were >95% pure, mis-assigned positional isomers. For ZINC901381520 (**Fig. 3**F), both N3 (PDB 5RSK) and N9 (PDB 5S6W) isomers were synthesized in different batches and confirmed to bind to the targeted adenosine binding pocket forming equal hydrogen-bond interactions with the protein (**Fig. S7**I). ZINC3888754 (PDB 5RSC) (**Fig. 3**G) contains an adenine-like heterocycle extended by methyl-groups at the C7 and C8 positions, revealing opportunities for expanding purine scaffolds beyond the adenine subsite to achieve Mac1 selectivity over other adenine-binding proteins.

In addition to hydrogen-bonding with residues involved in the adenine recognition of ADP-ribose, several docking hits hydrogen bond to the backbone carbonyl group of Ala154 (see **Fig. 3**D,I and **Fig. S7**G), thereby revealing an intriguing accessory polar contact within this sub-pocket. While most residues surrounding the adenosine-binding subpocket adopted similar conformations in the fragment-bound crystal structures as in the ADP-ribose-bound template protein structure used for docking (PDB 6W02) (*25*), Asp22 and Phe156 had increased dynamics in the electron density. In most fragment-bound crystal structures, Phe156 rotated by approximately 90° degrees, enabling improved π-π stacking (face-to-face) against the aromatic moieties in the bound fragments, as shown in **Fig. 3** C-G. However, the docking template orientation of Phe156 was retained for other pyrimidine-containing fragment-bound crystal structures (**Fig. 3**B,H).

Overall, two characteristics stand out from the docking screen: first, despite some important differences, there was high fidelity between the docking-predicted poses and those observed, to ultra-high resolution, by crystallography. The docking hits explored the adenine subsite to which they were targeted. Second, these hits did so with relatively dense variations around several chemotypes, something afforded by the granularity of a >20 million fragment library. This density can be explored further, for example, 9,170 fragments (888 unique BM scaffolds) in the ZINC15 fragment library contained 7H-pyrrolo(2,3-d)pyrimidines, the functional group repeatedly observed in crystallographically confirmed docking hits (**Fig. 3**A-C).

### Analysis of key interactions between Mac1 and fragments from the crystallographic screen

#### Fragments binding to the adenine subsite

While docking was successful in targeting the adenine binding pocket, crystallographic fragment screening has the advantages of being binding site agnostic and the potential to identify novel chemotypes at that site. Crystallographic screening identified 96 adenine-site binding fragments that form subsets of the three hydrogen bonds found between Mac1 and ADP-ribose (**Fig. 4**A-C). Fragments that formed at least two hydrogen bonds to the adenine subsite were separated into nine classes based on the number, nature and connectivity of atoms involved in such hydrogen bonding (**Fig. 4**D). The most common class consisted of a 1,3-hydrogen bond donor/acceptor motif (**Fig. 4**D,E.I). This is reminiscent of the hinge binding motifs observed in kinases, with the difference being the engagement of a side chain oxygen rather than the backbone carbonyl oxygen (**Fig. S8**B) (*57*). While 7 out of 18 fragments in this class were 4-amino-pyrimidine derivatives, other moieties were featured, including two 2-amino-thiazole-based fragments and several purine derivatives (**Supplementary data set 2**). We also observed an unusual adenine-binding mode with an H-bond formed between Ile23 and N7 instead of N1 (**Fig. 4**D,E.II). The alternative binding mode can be explained by the N3 substitution of adenine on this fragment, which prevents formation of the canonical N1-Ile23 H-bond. This pattern of hydrogen bonds to the protein has not been previously observed in adenines linked through N9 (*58*).

**Figure 4.**
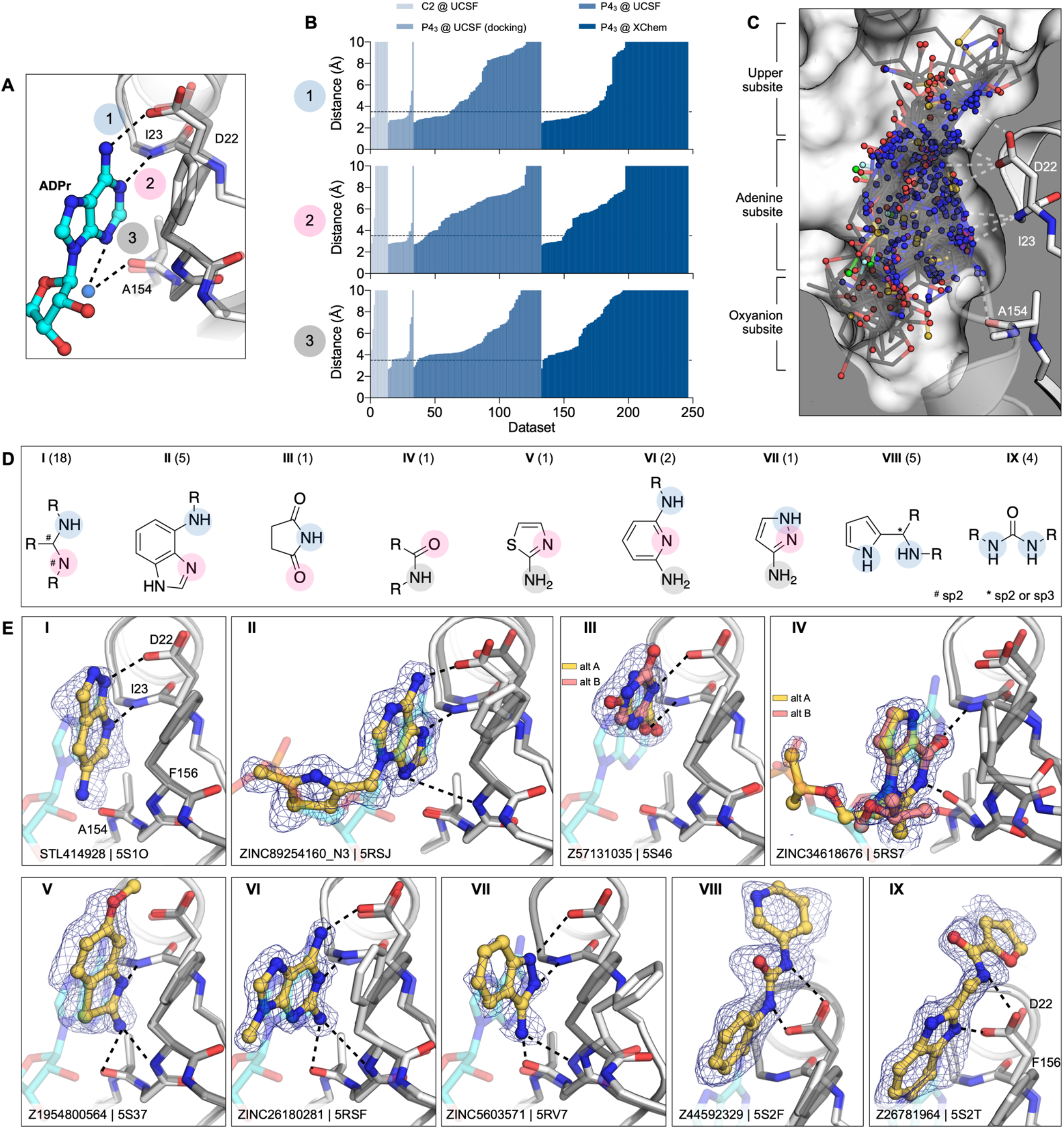
Fragments binding to the adenine subsite. **A)** Stick representation showing the interaction of the adenosine moiety of ADP-ribose with Mac1. The key interactions are shown as dashed lines. **B)** Plot of the distances shown in **(A)** for all fragment hits. The distances, truncated to 10 Å, are for the closest non-carbon fragment atom. **C)** Stick representation showing all fragments interacting with Asp22-N, Ile23-N or Ala154-O. The surface is ‘sliced’ down a plane passing through Asp22. **D)** Structures of the nine unique motifs that make at least two hydrogen bonds to the adenine site. Colored circles match the interactions listed in **(A)** and **(B)**. The number of fragments identified for each motif are listed in parentheses. **E)** Example for the nine structural motifs. The fragment is shown with yellow sticks and the PanDDA event map is shown as a blue mesh. ADP-ribose is shown as cyan transparent sticks. The apo structure is shown with dark gray transparent sticks.

We also observed diverse fragments without adenine-like motifs binding at this site, including succinimides, amides, thiazoles, diamino-pyridines, pyrazoles, pyrroles, and ureas (**Fig. 4** D,E.III-VIII). These exploited, separately and together, Asp22 and Ile23, Ala154, and occasionally all three adenine-defining hydrogen-bonding residues. Several fragments π-π stacked with Phe156, while those bearing a urea hydrogen-bonded with the carboxylate of Asp22 (**Fig. 4**D,E. VIII). These interactions were reproduced by a series of benzimidazole-based fragments (**Fig. 4**D,E. IX). These classes occupied what might be classified as an ‘upper subsite’, above that defined by the adenine-ribose axis, and may provide an opportunity to grow fragments away from the canonical site.

#### Fragments binding to the oxyanion subsite

As was true of several of the docking hits, 54 fragments formed interactions with an unexpected ‘oxyanion site’, defined by the backbone nitrogens of Phe156 and Asp157 adjacent to the adenine site (**Fig. 2** G). As suggested by its name, most of these fragments (48/56) were anionic (**Supplementary data set 1**). Intriguingly, the defining backbone nitrogens adopted a similar orientation to those defining the classic oxyanion hole of serine hydrolases such as acetylcholinesterase and trypsin (**Fig. S8**). In the ADPr-Mac1 structure, the C2 hydroxyl (2’OH) of the proximal ribose interacts with the oxyanion subsite via a bridging water (**Fig. 5**A). In total, 54 fragments formed at least one H-bond to the oxyanion subsite (**Fig. 5**B). Here too, the fragments were both geometrically (**Fig. 5**C) and chemically diverse (**Fig. 5**D): orienting groups either toward the phosphate tunnel, the lower site, or wrapped around toward the upper adenine site, providing multiple points of departure for further elaboration. Chemically, they interacted with the site using not only a carboxylate, but also sulfones, and isoxazole, α-keto acid, and a succinimide (**Fig. 5**E). We suspect that the presence of the oxyanion subsite explains the higher hit rate for the Enamine Essential library versus the other crystallographic fragment libraries screened (27% versus 6%), as this library contained a greater proportion of acids than the others (41% versus 4%) (**Fig. S5**).

**Figure 5.**
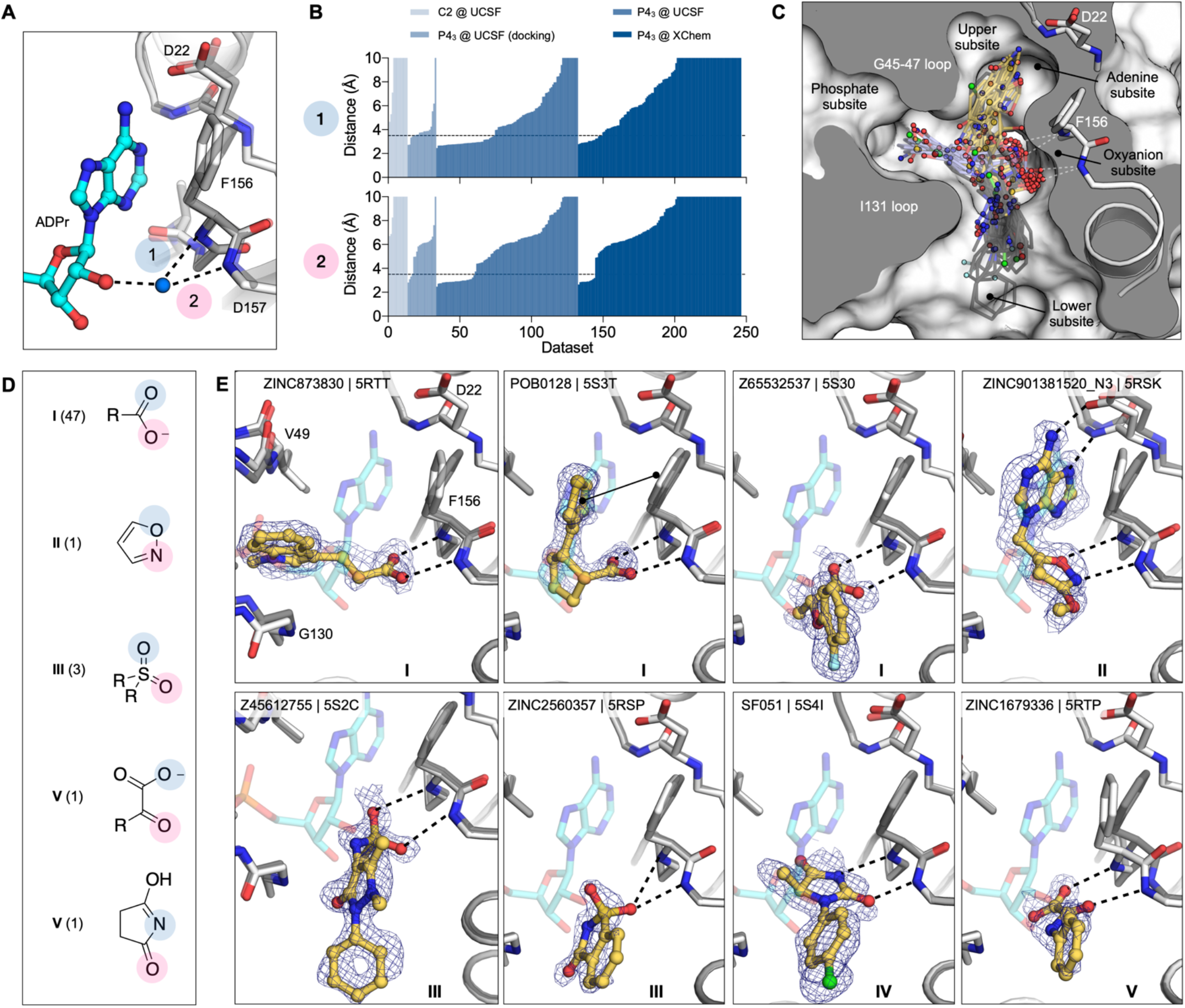
Fragments binding to the oxyanion subsite of the adenosine site. **A)** Stick representation showing the interaction of the adenosine portion of ADP-ribose bound to Mac1. The water molecule bridging the ribose moiety and the oxyanion site is shown as a blue sphere. **B)** Plot of the distances highlighted in **(A)**for all fragment hits. Distances were calculated as described for Figure 4B. **C)** Stick representation showing all fragments interacting with Phe156-N and Asp157-N. Fragments are colored by secondary binding site with blue = phosphate, black = lower and yellow = adenine. The surface is “sliced” across a plane passing through Phe156 (white surface, grey interior). **D)** Structures of the five unique motifs that bind the oxyanion site. **E)** Example for the five structural motifs. Three examples for motif I are shown, where the fragment also interacts with the phosphate, adenine or lower subsites. The fragment is shown with yellow sticks and the PanDDA event map is shown for reference as a blue mesh. ADPr is shown with transparent cyan sticks. The apo structure is shown with transparent gray sticks.

### Fragments binding to the catalytic and other potential allosteric sites

There were substantially fewer hits against the catalytic site (**Fig. 2**G) compared to the adenosine site (eight versus >100), though both appear to be accessible (**Fig. S3**). The catalytic site consists of three subsites: the phosphate tunnel, which is occupied by the diphosphate of the ADPr-diphosphate, the ribose site, which is occupied by the terminal ribose of the molecule, and the outer site, which sits between Asn40 and Asn99 (**Fig. 6**, **2**G). Of the eight fragments binding in the overall catalytic site, seven bound in the outer subsite and one bound in the phosphate tunnel. Binding to the outer site was often defined by hydrophobic packing between the Tyr42 and Lys102 side chains, although POB0135 (PDB 5S3W) and POB0128 (PDB 5S3T) formed a salt bridge to Lys102 (e.g. **Fig. 6**A.I). Interestingly, the latter fragment was also found to bind in the adenosine-binding site. Other molecules, including Z2234920345 (PDB 5S2L) and Z955123498 (PDB 5S4A) stabilize an alternative conformation of Lys102 (**Fig. 6**A.II). Three of the fragments, including Z85956652 (PDB 5S2U), positioned a halogen atom in the outer subsite (**Fig. 6**A.III). The only fragment identified in the phosphate subsite was ZINC84843283 (PDB 5RVI). This fragment was wedged between the Gly47/Ile131 loops, and increased the gap between the two loops by 1.6 Å (**Fig. 6**A.IV). The absence of fragments binding to the ribose subsite, and the sparsity of fragments in the phosphate subsite, means that designing a Mac1 inhibitor to occupy the catalytic site will rely more heavily on fragment growing than on fragment merging.

**Figure 6.**
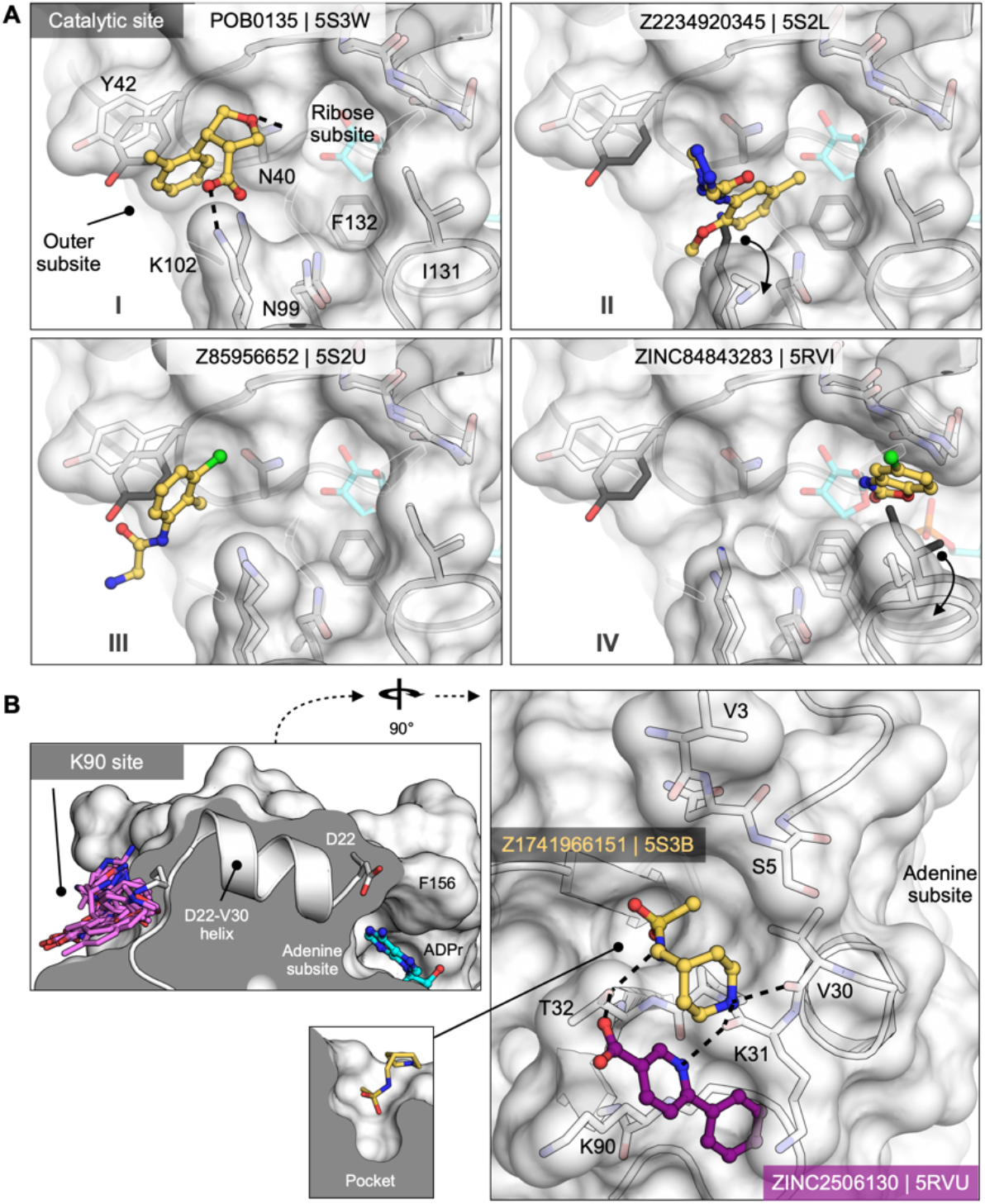
Fragments targeting the catalytic and potential allosteric sites are sparsely populated compared to the adenosine site. **A)**Surface representation showing fragments that bind near the catalytic site. The fragment POB0135 (PDB 5S3W) bridges the gap between Asn40 and Lys102 via a hydrogen bond and a salt bridge, respectively. Although eight fragments bind in the outer site, the fragment POB0135 makes the highest quality interactions. No fragments bind in the ribose subsite. The fragment in ZINC331715 (PDB 5RVI) inserts into the phosphate subsite between Ile131 and Gly47. **B)**Left: the K90 site is connected to the adenosine site via the D22-V30 alpha-helix. Right: surface representation showing two fragments that bind to the K90 site. Hydrogen bonds are shown as dashed black lines. The fragment in Z1741966151 (PDB 5S3B) is partially inserted in a nearby pocket (insert).

Both crystallographic screens also identified fragments binding to the ‘K90 site’, which is formed by a cleft between Lys31, Thr32 and Lys90 (**Fig. 6**B). We identified seven fragments from the C2 crystal form and six from the P4_3_ crystal form; intriguingly, none of the C2-derived fragments were found again when the UCSF libraries were rescreened in the P4_3_ crystal condition. Although the K90 site is 15 Å from the adenosine subsite, it is connected to that subsite via a single alpha-helix (**Fig. 6**B). Although there is no biochemical evidence for allosteric communication between these sites, the fragments provide starting points for designing chemical probes to test this possibility.

### Fragment binding exploits protein conformational flexibility

To identify Mac1 flexibility associated with molecular recognition, we calculated the root-mean-square fluctuation (RMSF) of side-chain atoms across the P4_3_ fragment-bound structures. Residues lining the adenosine site, especially Asp22 and Phe156, are the most flexible (**Fig. 7**A,B). The flexibility of both residues is paralleled in previously reported crystal structures (**Fig. 7**C, **Fig. S4**A) and also in the 0.77 Å apo structure, where multiple alternative conformations are clearly defined in electron density maps (**Fig. 7**D and **Fig. S1**A-C). In the high resolution structure, residues 155-159 are modeled as a combination of two distinct backbone conformations that diverge substantially at Phe156, which requires three distinct conformations of this residue to satisfy the observed density (**Fig. 7**D, **Fig. S1**C). Despite this flexibility, hydrogen bonds to Asp22 are present in many fragments, even docking compounds that were chosen based on interactions with a static receptor. Similarly, the flexibility of the aromatic side-chain of Phe156 enables adaptable stacking interactions with fragments (**Fig. 7**F), with 46 fragments binding within 4 Å to Phe156. As with Asp22, the nature and geometry of these interactions are maintained for many soaked and docked fragments even as the residue moves relative to the rest of the protein.

**Figure 7.**
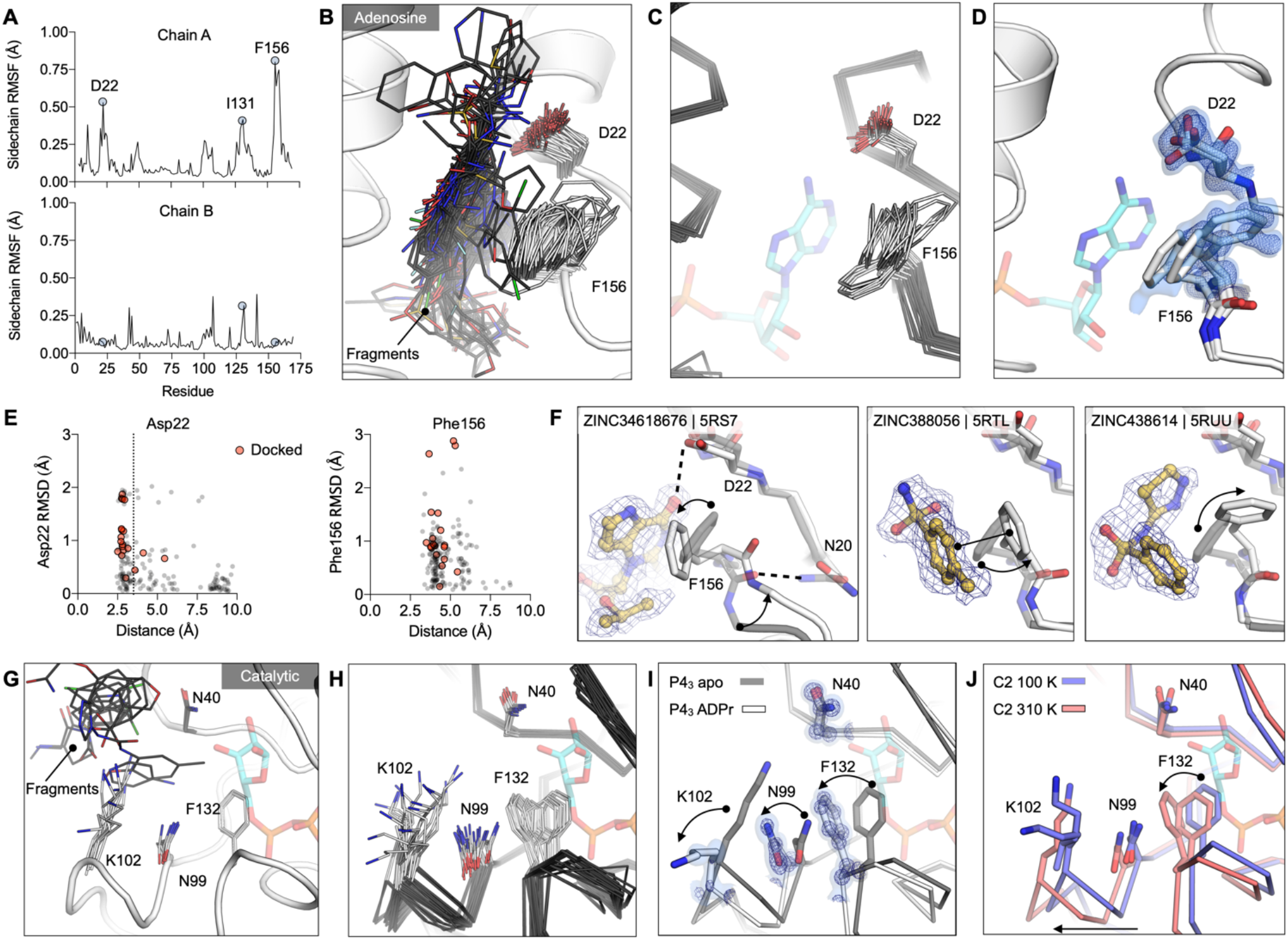
Experimentally observed conformational heterogeneity is sampled by various fragments. **A)** Plots of side-chain RMSF for the 117 fragment structures from the UCSF screen using P4_3_ crystals. **B)** Stick representation showing all fragments (black sticks) within 3.5 Å of the Asp22 carboxylate and 4 Å of the Phe156 ring (white sticks). **C)** Structural heterogeneity in the previously reported Mac1 structures. **D)** The Phe156 side-chain is captured in three conformations in C2 apo structure. Electron density maps (2mFO-DFC) are contoured at 0.5 σ (blue surface) and 1 σ (blue mesh). For reference, ADP-ribose is shown with blue sticks. **E)** Plots of side-chain RMSD for Asp22 and Phe156 from the Mac1 apo structure as a function of ligand-protein distance. Structures were aligned by their Cα atoms, before RMSDs were calculated for the Asp22 carboxylate and the Phe156 aromatic carbons. **F)** Fragment binding exploits preexisting conformational heterogeneity in the Phe156 side-chain. The apo structure is shown with dark transparent gray sticks in each panel and the conformational changes are annotated with arrows. **G)** Stick representation showing all fragments (black sticks) in the outer subsite of the catalytic site. **H)** Conformational heterogeneity of residues in the catalytic site of the previously reported Mac1 crystal structures. **I)** ADP-ribose binding induces a coupled conformational change in the Phe132, Asn99 and Lys102 side-chain, as well as a 2 Å shift in the Phe132 loop. Electron density maps (2mFO-DFC) are contoured at 1.5 σ (blue surface) and 4 σ (blue mesh). **J)** Mac1 structures determined at 100 K and 310 K.

In contrast to the adenosine site, little conformational heterogeneity is observed at the catalytic site, with only minimal changes in Lys102 and Tyr42 conformations (**Fig. 7**G). Still, even in this site, there is more conformational heterogeneity observed in previously published structures (**Fig. 7**H, **Fig. S4**A). In particular, a network of flexible side-chains encompassing Phe132, Asn99, and Lys102 is stabilized in a distinct conformation upon ADP-ribose binding (**Fig. 7**I). To further probe the flexibility of the Phe132-Asn99-Lys102 network, we determined structures of Mac1 using the C2 crystal at human physiological temperature (37°C, 310 K) to 1.5 Å resolution (**Fig. 7**J). As observed in other systems (*59, 60*), we noticed that the cryogenic structure appeared more compact than the structure at higher temperatures. Specifically, we observed substantial loop displacements near the ribose-binding pocket of the active site, which are coupled to a global hinge-bending motion involving correlated motion of helices about the central β-sheet (**Fig. S4**G). The structure at physiological temperature more closely resembles the structure with ADPr-bound, with the backbone adopting a more open conformation (**Fig. 7**J). However, the side-chain rotamers of Phe132, Asn99 and Lys102 do not undergo the larger rearrangements. This temperature-dependent change in the width of the active site cleft can provide alternative, potentially more relevant, conformations for future ligand-discovery efforts targeting the catalytic site around the distal ribose.

### Changes in water networks upon fragment binding

To assess the role of water networks in fragment binding, we examined changes in water networks upon ADP-ribose binding. In the 0.85 Å P4_3_ apo structure, the catalytic site contains 14 water molecules arranged in an ordered network that connects the Gly47 loop and the Ile131 loop, with an arc formed around the Phe132 side-chain (**Fig. 8**A). In contrast, waters were more disordered in the adenosine site, with more diffuse electron density and with waters modelled with higher B-factors (**Fig. 8**E). Upon ADP-ribose binding, six waters were displaced from the catalytic site and the water network was disrupted (**Fig. 8**B). This disruption is partly caused by altered conformation of the Phe132 and Asn99 side-chains, which break the network between residues Asn40 and Asn99. Conversely, the network in the adenosine site was stabilized in the Mac1-ADPr complex (**Fig. 8**B). The average B-factor decreased from 24 to 10, and two networks connect the phosphate tunnel with the adenine/oxyanion subsites (**Fig. 8**C). Although the adenine moiety only forms two direct hydrogen bonds to protein, it has four additional contacts via bridging water molecules (**Fig. 8**B). Similar bridging waters were observed for fragments binding in the adenosine site including ZINC2055 (PDB 5RSV), which forms only one direct H-bond to the protein, but has an extensive H-bond network via water molecules (**Figure 8**D). Visualizing all water molecules within 3.5 Å of fragment atoms shows clusters near protein hydrogen bond acceptors and donors (**Fig. 8**E). Of particular interest is the cluster near the backbone carbonyl of Ala154 (**Fig. 8**). This site is occupied by a water molecule in the ADPr-Mac1 structure, and is bridged by adenine derivatives such as ZINC340465 (PDB 5RSJ) (**Fig. 8**D). In addition, four fragments occupy this site directly (**Fig. 4**E), including the C2-amino-substituted adenine present in ZINC26180281 (PDB 5RSF, **Fig. 3**D). Extending fragments to displace the water molecules at other frequently populated sites could help to quantify the contribution of water networks to Mac1, and to provide a test set for computational methods that seek to exploit solvent dynamics for ligand optimization (*61*–*63*).

**Figure 8.**
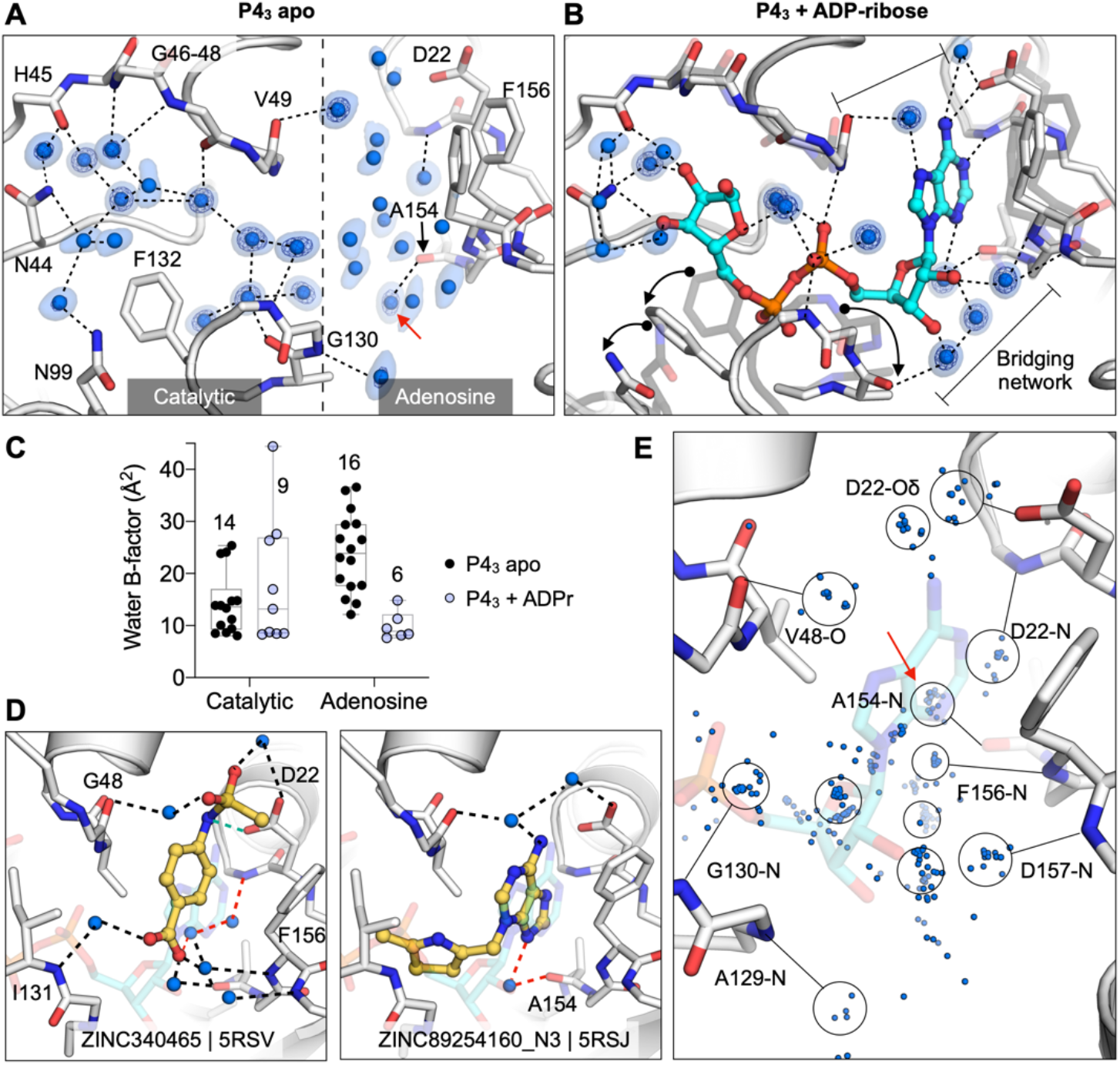
Water networks in the active site are displaced as well as used by fragments for bridging interactions. **A)** Water networks in the apo enzyme (P4_3_ crystal form). Waters are shown as blue spheres, with electron density contoured at 5 σ (blue mesh) and 1.5 σ (blue surface). Hydrogen bonds are shown as dashed lines (distances are 2.6-3 Å). **B)** Water networks in the Mac1-ADPr complex. ADP-ribose is shown as cyan sticks. Conformational changes upon ADP-ribose binding are highlighted with black arrows. **C)** Comparison of crystallographic B-factors of water molecules in the catalytic site and adenosine site. The range and 95% confidence interval are shown. **D)** Examples of the role of water networks in fragment binding. Right: ZINC340465 (PDB 5RSV) makes a single hydrogen bond to the protein (green dashed line), but makes five hydrogen bonds via water molecules. Although few fragments hydrogen bond directly to the backbone oxygen of Ala154 (Figure 4E), several fragments interact with this residue via bridging water molecules (red dashed line) including ZINC89254160_N3 (PDB 5RSJ). **E)** Plot showing all water molecules that lie within 3.5 Å of a non-carbon fragment atom. Water molecules are shown as blue spheres, with the major clusters circled. The cluster in a red circle bridges fragments and the Ala154 backbone oxygen.

### Characterization of in-solution binding of fragment hits

To buttress the crystallographic studies, selected compounds were biophysically screened using differential scanning fluorimetry (DSF) (*64*–*66*), isothermal titration calorimetry (ITC) (*67*–*69*), and a homogeneous time-resolved fluorescence (HTRF) ADPr-peptide displacement assay (**Fig. 9, Supplementary Table 1, Supplementary data set 2**). Because of their ready availability in useful amounts, most of these experiments focused on the docking hits. For DSF, in agreement with previous reports for this domain (*24*), we observed substantial elevation of the apparent melting temperature (Tm_a_) of Nsp3 Mac1 upon addition of its known binding partner, ADP-ribose (**Fig. 9**C,D,G). When tested in concentration-response from 0.188 to 3 mM, 10 of 54 docked fragments also induced small, but statistically significant and dose-responsive Tm_a_ elevation (**Fig. 9**C,D,G, **Supplementary Table 1, Supplementary data set 2**). All 10 of these were also observed to bind in the crystallographic studies, providing relatively good agreement between these assays. However, the correlation was incomplete, as the remaining fragments observed in crystallography either decreased the Tm_a_ or had no significant effect (**Supplementary Table 1**). Despite this, the results suggest that DSF is a rapid way to corroborate the fragment hits from crystallography and docking.

**Figure 9.**
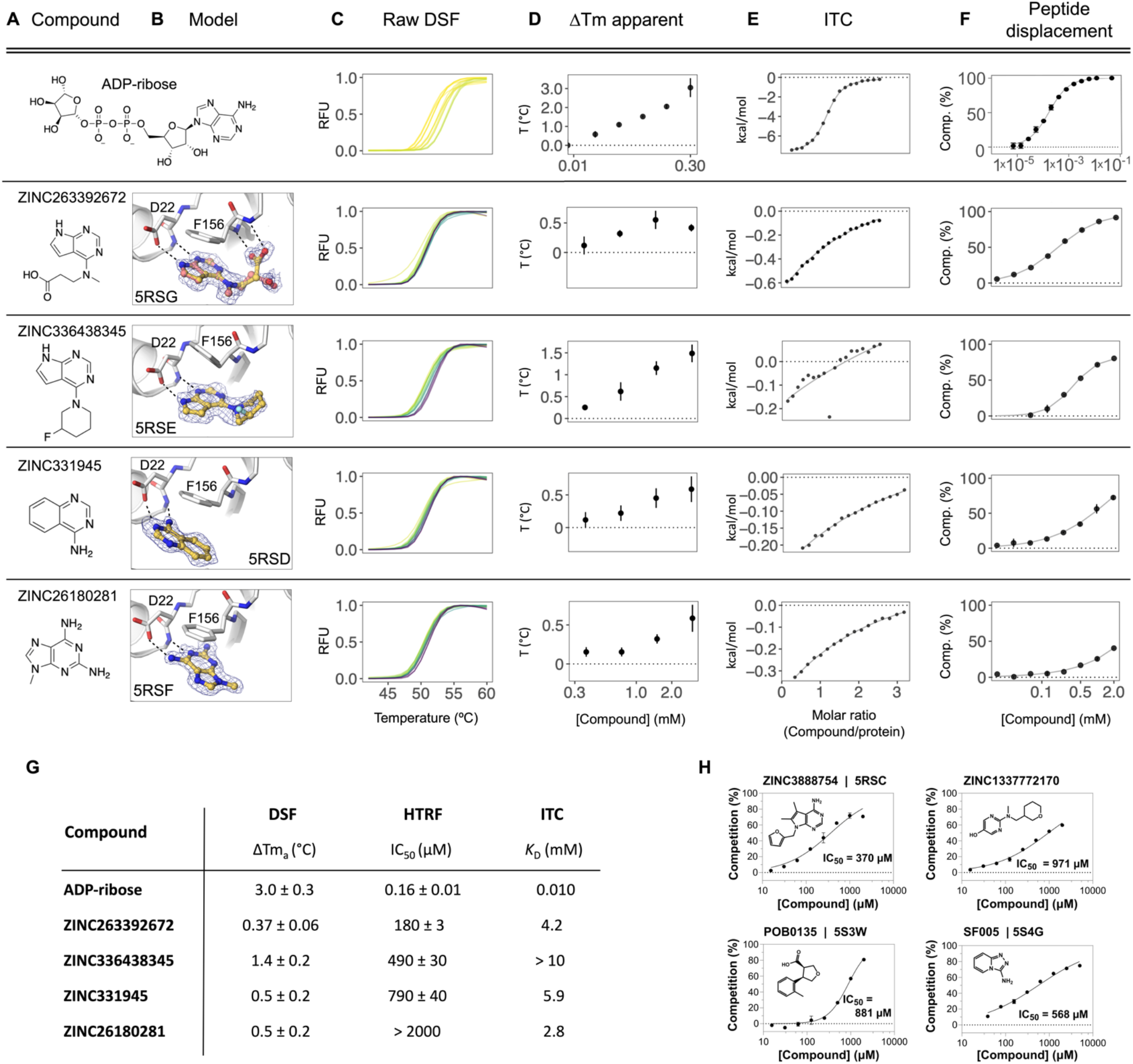
Biophysical corroboration of solution binding of crystallographic fragment hits by DSF, ITC and ADPr-peptide displacement assay. Top panel **(A-F)** shows performance of the most potent fragment hits in DSF, ITC, and ADPr-peptide displacement assay compared to ADP-ribose. **C,D)** Normalized raw DSF RFU demonstrates canonical unfolding curves and minimal compound-associated curve shape aberrations. Tm_a_ elevation reveals Mac1 stabilization through fragment binding. Gradient color scale: 0 mM = yellow; 3 mM = purple. **E)**Integrated heat peaks measured by ITC as a function of binding site saturation. The black line represents a non-linear least squares (NLLS) fit using a single-site binding model. **F)** Peptide displacement assay measures ADPr-peptide displacement (i.e. % competition) from Mac1 by ligand. **G)** Summary of solution binding data for fragments from top panel. ΔTm_a_ are given for the highest compound concentration in this assay. **H)** Additional fragment hits showing Mac1 peptide competition.

To identify fragments with the most promising binding affinity for optimization, we tested the 19 crystallographically observed docking hits using ITC. Due to their small size, most of these fragments have low binding affinity and release little heat upon binding versus ADP-ribose. Thus, we only observed reliable thermodynamic measurements for four of the 19 fragments. These could be fit to a 1:1 binding stoichiometry with affinities in the low mM range (**Fig. 9**E, **Supplementary Table 1, Supplementary data set 2**), consistent with the DSF results. Furthermore, the compounds measured by ITC that released the greatest amount of heat also induced the most significant Tm_a_ shifts in DSF, highlighting the complementarity between these two biophysical assays.

Finally, we tested 57 docking-derived fragments and 18 crystallographic hits from the XChem library in a HTRF-based peptide displacement assay, which monitors displacement of a fluorescently labeled ADPr-conjugated peptide from the active site of Mac1 by the fragment (**Fig. 9**F,G, **Supplementary Table 1, Supplementary data set 2**). Eight of 57 docking hits (14%) and three of 18 crystallographic hits (17%) inhibited the enzyme with IC_50_ values generally between 200 μM-1 mM, with the most potent fragment being the docking-derived ZINC263392672 (PDB 5RSG) with an IC_50_ of 183 μM in this assay. Only five (ZINC3888754 (PDB 5RSC), ZINC331945 (PDB 5RSD), ZINC263392672 (PDB 5RSG), ZINC336438345 (PDB 5RSE) and ZINC6180281(PDB 5RSF), **Fig. 3**) of the 10 docking hits that stabilized the macrodomain observed by DSF proved also inhibitory in the ADPr-peptide displacement assay. Interestingly, two docking hits which were not identified as binders by DSF or crystallography, ZINC1337772170 (IC_50_ = 971 μM) and pterin (IC_50_ = 790 μM), were found to be macrodomain inhibitors in this assay (**Fig. 9**H). This result might be explained with the use of a detergent-based buffer in the peptide displacement assay that could increase compound solubilization for what appear to be low-solubility compounds. With its ability to detect specific inhibition of the macrodomain, this ADPr-peptide displacement assay proved to be a sensitive and complementary strategy for further characterization of the fragment hits obtained from the docking and crystallographic screens. Assuming that the HTRF-based peptide displacement assay produced the most reliable affinity data, we estimated ligand efficiencies from IC_50_ values for hits for which we obtained reasonable dose-response curves. ADP-ribose with an IC_50_ of 161 nM and 36 heavy atoms indicates a ligand efficiency (LE) of 0.26 kcal/mol per non-hydrogen atom. The docking hits ZINC3888754 (PDB 5RSC, LE=0.26), ZINC336438345 (PDB 5RSE, LE=0.28), ZINC263392672 (PDB 5RSG, LE=0.32) and ZINC331945 (PDB 5RSD, LE=0.38) reveal similar or slightly improved ligand efficiencies, while the highest LE was calculated for the XChem library hit SF005 (PDB 5S4G, **Fig. 9**H) with 0.44 kcal/mol per heavy atom.

In summary, all crystallographically confirmed docking hits were tested using three complementary in-solution binding techniques - DSF, ITC, and an HTRF-based peptide displacement assay (**Fig. S9, Supplementary Table 1, Supplementary data set 2**) (*70*). ZINC336438345 (PDB 5RSE), ZINC331945 (5RSD), ZINC263392672 (5RSG) and ZINC26180281 (5RSF) were the only four fragment hits for which binding data could be obtained by all three techniques (**Fig. 9**). All of these fragments have key hydrogen bonds in the adenine site and display π-π stacking with Phe156. Furthermore, ZINC263392672 (PDB 5RSG) can additionally interact via its carboxyl group with the oxyanion site of Mac1. Finally, we note that crystallography, DSF, ITC all monitor binding, but do not measure function. The peptide displacement assay is thus of particular value for fragment characterization, since it measures specific displacement of an analog of the natural Mac1 substrate.

### Opportunities for fragment linking and merging to optimize Mac1 inhibitors

Typically, one might be reluctant to speculate on optimization from fragment structures alone, but the unusually large number of structures, perhaps supports some cautious inference here. Prior to modifying, linking, or merging fragments, it is important to consider the crystalline environment. In the macrodomain crystal, the active site forms a bipartite enclosed pocket with a symmetry mate (**Fig. 10**A). In particular, 24 fragments only hydrogen bond to Lys11 of the symmetry mate, and do not hydrogen bond with any residues in the adenosine site, indicating that these molecules should not be considered for fragment elaboration (**Fig. 10**C,D). Based on the solved binding poses of remaining compounds, fragment pairs were linked into hypothetical scaffolds. These were used as templates to search the make-on-demand chemical space of the Enamine REAL database employing the smallworld similarity (http://sw.docking.org) and arthor substructure (http://arthor.docking.org) search engines (**Fig. 10**E,F) (*71*). In a second approach, fragments with overlapping binding poses were merged into larger scaffolds, e.g. the purine of ZINC89254160_N3 (PDB 5RSJ) interacting in the adenine binding pocket was replaced by ZINC26180281 (PDB 5RSF) adding an additional hydrogen bond to Ala154 (**Fig. 10**F). Whereas it remains speculative whether the suggested linked or merged molecules are indeed active against the macrodomain, the scaffolds observed here, and the key interactions they make with the enzyme, indicate a fruitful chemical space to further explore. Naturally, many of the fragments described here also merit investigation by alternative fragment growing or analoging strategies.

**Figure 10.**
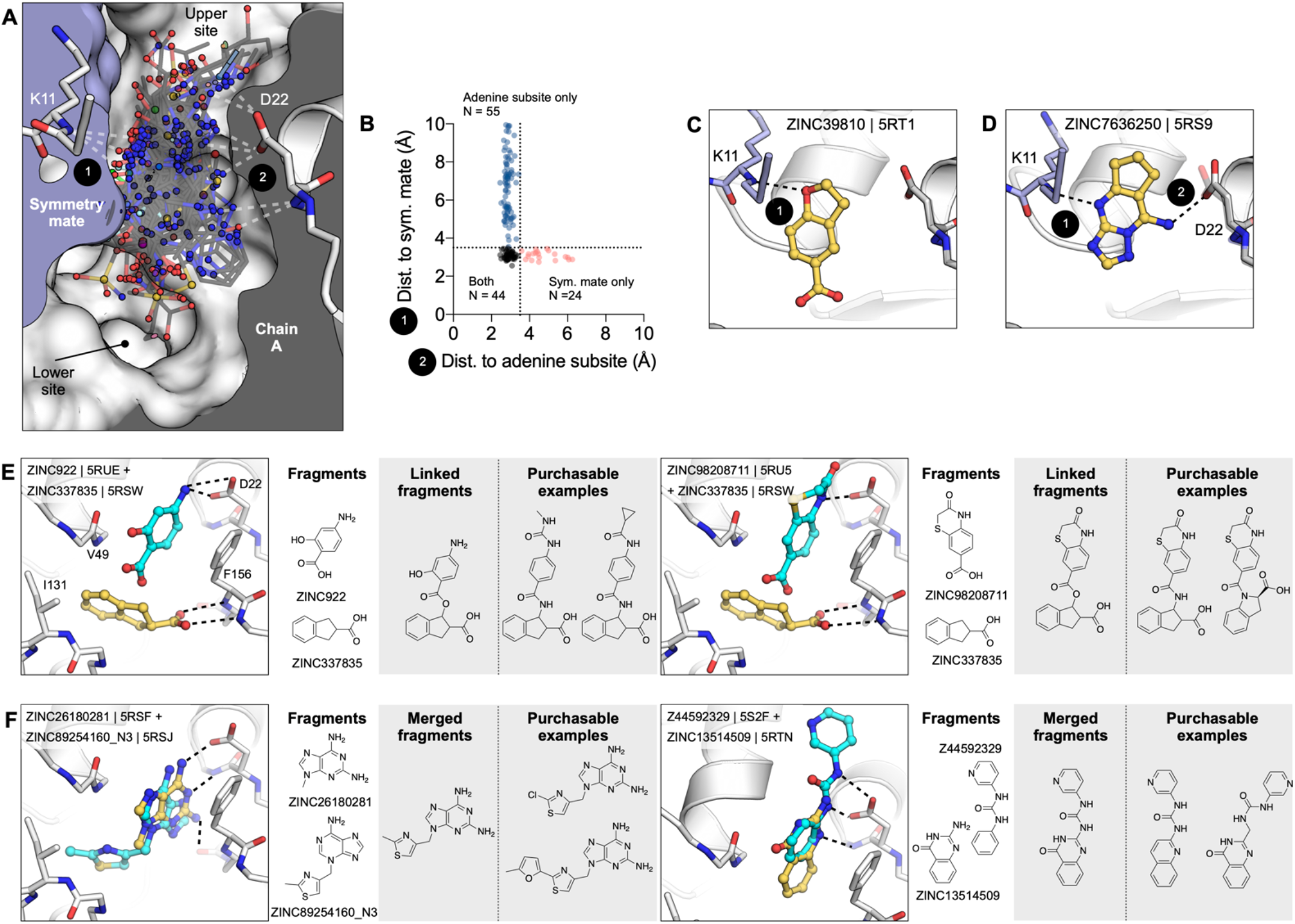
Fragments bridging multiple adenosine subsites provide direct merging opportunities. **A)** Sliced view of the adenosine site (white surface, grey interior) and a symmetry mate (blue surface and interior) showing the deep pocket created by crystal packing in the P4_3_ crystals. The 66 fragments that H-bond with the Lys11 backbone nitrogen are shown as sticks. **B)** Plot showing distances between the symmetry mate (Lys11-N) and the adenine subsite (Asp22-Oδ, Ile23-N, Ala154-O) for all fragments identified in the adenosine site. Dashed lines show the 3.5 Å cut-off used to classify H-bonds. **C)** An example showing one of the 24 fragments that bound in the adenosine site, yet only formed an H-bond with the symmetry mate. **D)** An example of one of the fragments that bridged the 9-11 Å gap between the adenine subsite and the symmetry mate. **I, J)** Opportunities for fragment linking and merging. Adjacent or overlapping fragments were initially merged into a single new compound. Examples of readily available make-on-demand compounds are shown.

## DISCUSSION

Three key observations emerge from this study. Most noteworthy is the sheer number and the unusually high resolution of the 234 fragment-bound Mac1 structures reported here, including 192 fragments identified in the active site. The fragments cover both stereotyped interactions (such as adenine-like hydrogen bonding to the Asp22 side chain/ Ile23 backbone and stacking interaction with Phe156) as well as chemical diversity exploiting flexibility in the active site (for instance targeting of the oxyanion-subsite). This abundance and diversity afford robust starting points for future elaboration into lead-like molecules. Second, the high fidelity of docked poses to the subsequent crystallographic results supports the use of docking to explore the adenine recognition site, and importantly, demonstrates an ability of docking to prioritize fragments in this case, an approach still debated in the field. Finally, with 234 diverse fragment structures determined, it should be possible to exploit the fortuitous juxtaposition of pairs of fragments to design joined ligands that combine the affinities of both, leading to robust starting points with the low micromolar affinity needed for initiating hit-to-lead optimization of inhibitors. One clear strategy involves extending molecules bound to the adenine site and with biophysically measurable binding affinities into the phosphate and ribose recognition regions.

In contrast to the large number of chemically diverse hits binding to the adenine site, the lack of fragments bound to the catalytic site is notable and may inform models of how the ADP-ribosylated peptide may bind to the macrodomain. This result also suggests that this is not likely to be a productive region for growing fragments which are binding in the adenine site. This paucity of fragments is especially surprising given that three crystal environments (the A and B chains in the P4_3_ crystal and the C2 crystal) were screened and that the site appears accessible in all lattices (**Fig. S2**). The two major models for peptide-macrodomain interactions are either that the peptide binds along the widened cleft defined by Tyr42 and Lys102, or that it extends into solution through the flexible Gly46-48 loop (*72*). Indeed, we observe fragments that bind in both locations (**Fig. 6**A). Regardless of the binding mode, which could be distinct depending on the identity of the modified residue and target substrate, the lack of binding at this site suggests that the binding energy comes almost exclusively from the ADP-ribose and not from the amino acids on the ADPr-conjugated protein. This hypothesis is also supported by the fact that Mac1 can hydrolyse a wide range of ADP-ribosylated substrates (*2, 3*). Docking of larger ‘lead-like’ molecules, perhaps enabled by the expanded catalytic site revealed by the physiological temperature structure, and detailed description of solvent may help to identify molecules exploiting this site.

The success of the fragment docking campaign contrasts, perhaps, with expectations of the field that fragments have too few functional-group handles to accurately dock or prioritize (*73, 74*). Not only were hit rates high (33%), so too was the fidelity of most docking poses to the crystallographic results. Even judged by potency, the most active fragment to emerge from this study, the 183 μM inhibitor ZINC263392672 (PDB 5RSG) (**Fig. 3 and 9**), was a docking hit. Also, it was the docking hits that were most readily available for such functional testing, as they were sourced as 10 mg of dry powder, while the crystallographic screening compounds were often in short supply. This is a purely mechanical advantage of docking, and it is counterbalanced by the small numbers tested versus the crystallographic screens; still, having substantial material to work with is a pragmatic advantage. Admittedly, weaknesses also emerged from the docking. Intriguingly, the oxyanion site that featured so prominently among the crystallographic screening hits were not to be found among the docking predictions. This gap reflects both a failure of the docking scoring function to prioritize anions binding to this site (as they were at least sampled), and to some extent a failure of the docking group to pick the few molecules that did dock well to this site as likely candidates. More broadly, as we docked against a single rigid structure of the protein, the subsequent conformational changes that the protein underwent, and the changes in the water network, were not captured in the docking predictions, and this was sometimes reflected in the larger RMSD differences between predicted and observed fragment poses. These caveats, important as they are, should not obscure a central observation from this study: the docking hit rate was not only high, but the hits were typically right for the right reasons; this may be something to build on for the field.

From the docked compounds, the most promising hits identified by in-solution binding experiments were also crystallographically confirmed. However, as expected, the majority of soaking hits did not show appreciable activity in the orthogonal biophysical assays within the tested concentration range (up to 10 mM in ITC, **Supplementary Table 1**). The macrodomain ADPr-peptide displacement assay also identified two docking hits not previously observed in soaking (ZINC1337772170 and pterin), which suggests that the crystal environment limited the ability of some fragments to bind. Yet, between solution experiments good consensus across all assays was observed for ZINC263392672 (PDB 5RSG), ZINC336438345 (PDB 5RSE) and ZINC331945 (PDB 5RSD). While we are aware that obtaining high-quality binding data remains particularly challenging to determine for weak binders such as fragments, the dose-response results obtained in the complementary assays for many of the identified hits provided convincing evidence for their true binding to the macrodomain. The inconsistency of fragment binding is apparent when comparing fragments that resulted in high quality data sets in both the P4_3_ and C2 crystal systems. Surprisingly, only five of 59 possible fragments were observed in both systems, with three fragments binding with equivalent poses in the adenine subsite. This observation points to the value of having multiple measurements, and even multiple crystal systems when they are available, in fragment-based drug discovery approaches.

Overall, this study has three main implications for inhibitor discovery of SARS-CoV-2 Nsp3 macrodomain, and for antiviral efforts targeting this domain more broadly. First, we describe not only the first new chemical matter for this target, but map its hot spots at high resolution. This provides a template for future inhibitor discovery and development against this enzyme. Such efforts will need to navigate selectivity over human macrodomains but also ATP-binding proteins including kinases (**Fig. S8**) and consider breadth across other viral macrodomains (*16*) (**Fig. S4**). Second, the specific fragments that we describe may lend themselves directly to optimization: several examples are discussed explicitly, amenable to make-on-demand chemistry (**Fig. 10**); and the 234 structures should provide inspiration for countless other molecules. Finally, important technical advances emerged from this study: a crystal form that lends itself to ready structure determination, the creation of a reliable peptide-displacement assay for the macrodomain, and evidence supporting the ability of structure-based screening, such as molecular docking, to predict effective fragments. The ability to position explicit hydrogen atoms experimentally, and to resolve electron density on a subatomic scale, makes Mac1 an attractive candidate for in-depth computational dissection of its catalytic mechanism using approaches that integrate both classical and quantum calculations. Taken together, these advances will speed progress throughout the community to help validate this target and create effective antivirals.

## MATERIALS AND METHODS

### Fragment libraries

We screened 2,126 molecules from the XChem facility at Diamond Light Source and 411 molecules from UCSF against Mac1 crystallised in P4_3_ crystal form (**Supplementary data set 1**). The fragment library at XChem combined molecules from multiple fragment libraries: the Diamond, SGC and iNEXT (DSI)-poised Library (768 molecules, (*34*)), the Edelris fragment collection (280 molecules, (*75*)), the MiniFrags Probing Library (80 molecules, (*76*)), the FragLites collection (31 compounds, (*77*)), the PepLite library (25 molecules, (*38*)), the SpotFinder (~100 compounds, (*78*)), and the York3D (106 molecules, (*79*)) and the EU Open screen (969 molecules, (*52*)).

The UCSF fragment library was composed of Enamine’s Essential Fragment library (320 compounds) and 91 additional compounds from an in-house library (UCSF_91). To assemble the UCSF_91 library, we selected topologically diverse molecules having over 10,000 commercially available analogs in at least three points of substitution, allowing for rapid and extensive analog-by-catalog without having to resort to flask synthesis. We picked molecules that were also Bemis-Murcko scaffolds (*80*), stripped of acyclic terminal substituents. We thought simple, unsubstituted frameworks would be easier to optimize by adding chemical matter during analoging. From among these, we prioritized by eye scaffolds with various ring sizes and combinations including fused rings, spiro systems, with linkers of varying lengths between rings, in an attempt to sample a diverse range of compact shapes and properties. We added anions where the anionic moiety was a small acyclic substituent on the scaffold, again picking by eye for shape diversity. We chose molecules with 11-21 heavy atoms, with molecular weights between 200-300 amu and with a logP < 2.5 for solubility. Physical properties of all screened libraries are shown in **Fig. S5**.

Analyses of scaffolds and specific chemotypes in the employed chemical libraries are shown in **Fig. S5**E. Bemis-Murcko (BM) scaffold analysis was performed with the molinspiration mib engine (http://www.molinspiration.com) (*81*). Pyrimidines were identified using RDKit (http://www.rdkit.org) (*82*) and molecular charges at pH 7.4 were approximated using ChemAxon Jchem version 2019.15 (http://www.chemaxon.com) to identify anionic fragments (*83*).

### C2 crystals at UCSF

#### Protein expression and purification

SARS-CoV-2 Nsp3 Mac1 (residues 2-170) was cloned into a pET22b(+) expression vector with an N-terminal His_6_-tag and a TEV protease recognition site for removal of the tag (GenScript; Piscataway, NJ). In addition, a short linker (Asn-Ala-Gly) was included between the TEV recognition site and the Mac1 gene (**Supplementary data set 1**). To express Mac1, plasmid DNA was transformed into BL21(DE3) *E. coli*. After overnight growth on LB agar supplemented with carbenicillin (100 ug/ml), starter cultures (10 ml LB) were grown at 37°C for 8 hours. Large scale cultures (1 l TB) were grown at 37°C until an optical density of 0.8. Cultures were cooled at 4°C for 15 minutes, before protein expression was induced with 1 mM IPTG, and the cultures were shaken at 20°C for 12 hours. Cells were collected by centrifugation and frozen at −80°C. All purification steps were performed at 4°C using an AKTA FPLC system (Cytiva). Cells were resuspended in Ni-NTA binding buffer (50 mM Tris-HCl pH 8.0, 500 mM NaCl, 10 mM imidazole, 5% glycerol, 2 mM βME supplemented with 5 units/ml TurboNuclease (Sigma, T4330)) and lysed by sonication. Cell debris was collected by centrifugation and the lysate was applied to a 5 ml HisTrap HP column (Cytiva, 17524802). The column was washed with 25 ml binding buffer followed by 25 ml 5% Ni-NTA elution buffer (50 mM Tris-HCl pH 8.0, 500 mM NaCl, 500 mM imidazole, 5% glycerol, 2 mM β-ME), and then eluted with 100% elution buffer. Eluted protein was exchanged into TEV reaction buffer (50 mM Tris pH 8.0, 100 mM NaCl, 1 mM DTT and 1% glycerol) using a HiPrep 26/10 desalting column (Cytiva, 17508701). To cleave the His_6_ tag, Mac1 was diluted to 1.5 mg/ml using TEV reaction buffer and incubated with recombinant TEV protease (*84*) at a 1:20 ratio (Mac1:TEV) for 16 hours at 4°C. Cleaved Mac1 was separated from the uncleaved protein and TEV protease by re-running the sample over a HisTrap HP column (pre-equilibrated with TEV reaction buffer) and collecting the flow-through. The flow-through was supplemented with 10 mM DTT and concentrated to 2.5 ml using a 10 kDa molecular weight cut-off (MWCO) centrifugal concentrator (Amicon, UFC901024). The sample was further purified by size-exclusion chromatography (SEC) using a HiLoad 16/600 Superdex 75 pg column (Cytiva, 28989333) and SEC buffer (20 mM Tris pH 8.0, 150 mM NaCl, 5% glycerol, 2 mM DTT). Eluted fractions were concentrated to 15 mg/ml and flash-frozen in liquid nitrogen, and stored at −80°C. Protein used for ITC was purified in the same manner, but SEC was run with 150 mM NaCl, 20 mM Tris pH 8.0. Protein was concentrated to 10.8 mg/ml prior to flash freezing in liquid nitrogen and storage at −80°C.

#### Crystallization

Crystals were grown at 19°C using sitting-drop vapor diffusion with a reservoir solution containing 100 mM Tris pH 8.5, 100 mM sodium acetate and 28% PEG 4000. Crystallization drops were set up with 200 nl protein and 200 nl reservoir. Initially, crystals were grown in MRC 2-well plates (SwissCI, MRC96TUVP) with a reservoir volume of 40 μl. Crystals grew to a maximum size after 1-2 days, and were vitrified in liquid nitrogen without additional cryoprotection. For diffraction experiments at physiological temperatures, crystals were mounted using ALS-style goniometer bases (Mitegen, GB-B3S) and sealed with plastic capillary and vacuum grease (Mitegen, RT-T1). The capillary contained 4 μl reservoir solution to prevent crystal dehydration.

Fragment soaking was performed using crystals grown with SwissCI 3-well plates (SwissCi, 3W96T-UVP). Microseeding was required to achieve consistent nucleation. Several large crystals grown in 100 mM Tris pH 8.5, 100 mM sodium acetate and 28% PEG 4000 were transferred to a drop containing 5 μl seed storage buffer (100 mM Tris pH 8.5, 100 mM sodium acetate, 32% PEG 4000, 2 mM DTT) on a silicon coverslip (Hampton Research, HR3-233). Crystals were crushed using a flattened glass rod and transferred to 200 μl of seed storage buffer, before being serially diluted 1:10 with seed storage buffer. Consistent nucleation was achieved with seeds at a 1:100 dilution, with crystallization drops containing 200 nl reservoir, 100 nl seed stock and 300 nl protein with 30 μl in each reservoir.

#### Crystal dehydration and fragment soaking

Fragments were added to crystallization drops using acoustic dispensing with an Echo 650 liquid handler (Labcyte) (*35*). Two libraries were soaked at UCSF: the Enamine Essential fragment library (Enamine, 320 fragments), and the UCSF_91 library (91 fragments) (**Supplementary data set 1**). To limit DMSO-induced crystal damage, fragments were targeted to crystallization drops as far away from crystals as possible (*35*). Initial DMSO tolerance tests indicated that the C2 crystals were sensitive, rapidly disintegrating upon soaking with 10% DMSO (**Fig. S2**). To enhance DMSO tolerance, 300 nl of a solution containing 35% PEG 400, 100 mM Tris pH 8.5 and 50 mM sodium acetate was added to drops containing crystals using the Echo. Plates were resealed and incubated at 19°C for 6 hours. Fragment solutions (120 nl, 10% of the drop volume) were added using the Echo, and plates were re-sealed and incubated at 20°C for 3-8 hours. Crystals were vitrified directly from crystallization drops without additional cryoprotection..

#### Lysine methylation

Lysine methylation is a routine strategy for altering the crystallization properties of a protein (*51*). All reagents were added with the protein on ice and incubation steps were performed at 4°C with gentle shaking. First, 20 mg Mac1 was exchanged into lysine methylation buffer (50 mM HEPES pH 7.5, 150 mM NaCl, 5% glycerol) using a HiPrep 26/10 desalting column (Cytiva, 17508701). The protein was diluted to 1 mg/ml with lysine methylation buffer, and 400 μl 1 M dimethylamine borane (DMAB, prepared in water) (Sigma, 180238) and 800 μl 1 M formaldehyde (prepared in water) (Sigma, F8775) were added to initiate the methylation reaction. The reaction was left to proceed for 2 hours, then 400 μl 1 M DMAB and 800 μl 1 M formaldehyde was added. After an additional 2 hours, 200 μl 1 M DMAB was added and the reaction was left for a further 16 hours. To consume any remaining formaldehyde, and to cleave any intermolecular disulfide bonds, 2.5 ml of 1 M glycine (prepared in water) and 2.5 ml of 50 mM DTT (prepared in water) was added and the reaction was incubated for an additional 2 hours. Next, the sample was concentrated to 2.5 ml using a 10 kDa MWCO concentrator, and purified by SEC. The methylated protein was concentrated to 15 mg/ml before flash freezing in liquid nitrogen and storage at −80°C.

To test the extent of lysine methylation, the purified sample was analysed by liquid contromatogrpahy-mass spectrometry (LC-MS), using a Waters Acquity LC connected to a Waters TQ detector with electrospray ionization. The sample was separated on a C4 column held at 40°C using water with 0.1% formic acid as solvent A and acetonitrile with 0.1% formic acid as solvent B. After sample injection (5 μl at 10 μM diluted in 150 mM NaCl, 20 mM Tris pH 8.0), an isocratic elution was run with 95% solvent A and 5% solvent B for 1.5 min. Then, a linear gradient elution was run for 6.5 min to 95% solvent B. Finally, an isocratic elution was run with 95% solvent B for 2 min. The flow rate was 0.2 ml/min.

#### Crystallization of methylated Mac1

Crystals grew readily in the same conditions as the non-methylated protein (100 mM Tris pH 8.5, 100 mM sodium acetate, 28% PEG 4000). Consistent nucleation was achieved using microseeding with the same protocol as the non-methylated protein. Crystallization drops were set up with 100 nl reservoir, 100 nl seed stocks and 200 nl protein using SwissCI 3-well plates. The methylated crystals displayed increased DMSO tolerance, so DMSO/fragment soaks were performed directly with 40 nl DMSO (10% of the drop volume).

#### Ultra high resolution data collection, refinement and modelling

To measure the diffraction at such high resolution, we employed a multi-pass, multi-crystal data collection strategy. We collected ultra high resolution X-ray diffraction data for Mac1 (C2 crystal form) by performing sequential high-energy (17000 eV) and low-energy (11111 eV) runs to accurately measure reflection intensities at high and low scattering angles respectively. The same data collection strategy (wedge, oscillation angle, exposure) was implemented for multiple crystal specimens, each held in different orientations relative to the X-ray beam and phi rotation axis.

The data sets were individually indexed and integrated with XDS (*85*). During data processing, we merged the high and low resolution datasets from multiple crystals in different orientations to maximize our coverage of reciprocal space given a square detector surface. A low-resolution cutoff of 2.5 Å was applied to the high-resolution (high energy) data sets, because this cutoff simultaneously excludes potentially overlapping reflections at low scattering angles and allows for a significant number of shared observations between high and low resolution data sets, which facilitates robust scaling. Scaling and merging were performed using XSCALE, and the merged intensities were converted to structure factor magnitudes using XDSCONV (*85*). We calculated phases by the method of molecular replacement, using the program Phaser (*86*) and a previous structure of Mac1 (PDB 6WCF) as the search model. The model was manually adjusted in Coot to fit the electron density map calculated from molecular replacement, followed by automated refinement of coordinates, atomic displacement parameters, and occupancies using phenix.refine (*87*) with optimization of restraint weights. Following two initial rounds of iterative model building and refinement using the aforementioned strategy, we began introducing additional parameters into the model, enabled by the extraordinarily high resolution of our diffraction data. First we implemented anisotropic atomic displacement parameters for heavy atoms (C,N,O,S), followed by refinement of explicit hydrogen atom positions. During early rounds of model building, we noticed mFo-DFc difference density peaks appearing between heavy atom positions, suggesting that we are able to resolve covalent bonding densities (**Fig. S1**). Indeed, atomic refinement that included a model for inter-atomic scatterers (IAS) was able to account for these densities (*88*). Final refinement was performed without geometry or ADP weights (unrestrained).

#### Data collection at physiological temperature, refinement and modelling

We used a low-dose X-ray data collection strategy to acquire diffraction data from macrodomain crystals (C2 crystal form) at human physiological temperature (37°C, 310 K), which is the temperature most relevant to studies of SARS-CoV infection. Using this strategy, we acquired data sets using an X-ray exposure of only 50 kGy - less than 1% of the total dose used at 100K, which is essential to mitigate the rapid rate of radiation damage at 310 K compared to 100 K. The lower overall X-ray dose resulted in data with a lower overall resolution, extending to 1.5 Å.

Diffraction data from multiple crystals were merged using xia2 (*89*), implementing DIALS (*90*) for indexing and integration, and Aimless (*91*) for scaling and merging. We calculated phases by the method of molecular replacement, using the program Phaser (*86*) and our high resolution 100K structure as the search model. The model was manually adjusted in Coot (*92*) to fit the electron density map calculated from molecular replacement, followed by automated refinement of coordinates, atomic displacement parameters, and occupancies using phenix.refine (*87*) with optimization of restraint weights.

#### Fragment data collection, refinement and modelling

Diffraction data was collected at ALS beamline 8.3.1 and SSRL beamlines 12-1 and 12-2. The data collection strategy is summarized in **Supplementary data set 1**. Fragment datasets were indexed, integrated and scaled using XDS (*85*) run through xia2 (*89*). Based on the space group and unit cell dimensions, six crystal forms were present (**Fig. S2**. For each of the three C2 isoforms with one molecule in the ASU (isoform A, B and C), a single, high resolution dataset was selected to create a representative model for each isoform. Phases were obtained via molecular replacement with Phaser (*86*), using the ultra-high resolution C2 coordinates as the search model (PDB 7KR0). Coordinates were refined with iterative rounds of manual model building in Coot (*92*) and refinement with phenix.refine (*87*). Default refinement parameters were used, except five refinement macrocycles carried out per iteration and water molecules were automatically added to peaks in the 2mFO-DFC electron density map higher than 3.5 σ. The minimum model-water distance was set to 1.8 Å and a maximum model-water distance to 6 Å. For later rounds of refinement, hydrogens were added to riding positions using phenix.ready_set, and B-factors were refined anisotropically for non-hydrogen and non-water atoms. Although these datasets were obtained from crystals soaked with fragments, three was no evidence for fragment binding in the mFO-DFC difference density maps, therefore the datasets were deemed acceptable as representative DMSO-only models for each isoform.

For the fragment datasets, molecular replacement was performed with Phaser (*86*) and initial refinement with Refmac (*93*), both run through the DIMPLE pipeline (*91*). The search model used for molecular replacement was selected to match the isoform of the dataset. Waters were included in the initial refinement by changing the HOH records in the PDB file to “WWW”. After refinement, waters were stripped from models and electron density maps were analyzed for fragment binding using PanDDA (*54*). Electron density maps from 31 datasets were used to calculate the background electron density map for the A isoform, and 24 datasets were used for isoforms B and C (**Supplementary data set 1**). Datasets selected for background map calculation had the highest resolution and lowest *R*free values. After PanDDA was run with default parameters, the threshold used to classify a hit was decreased by adjusting the Z-map analysis settings (contour_level = 2, min_blob_volume = 5, min_blob_z_peak = 2.5). Although there was a substantial increase in false positives, the decreased threshold allowed an additional seven fragments to be identified. Fragments were modelled into PanDDA event maps with COOT, using restraints generated by phenix.elbow from a SMILES string. Changes in protein conformation and solvation were also modeled. Because PanDDA can identify fragments binding with low occupancies, any changes in protein coordinates will have similar, low occupancies. If un-restrained refinement is performed on these low occupancy models, changes supported by PanDDA event maps are often reverted to the ground state model. In the past, this has been overcome by refining both ground-state (apo) and changed-state (fragment bound) structures simultaneously, with the changed state coordinates restrained. However, these multi-state models can be difficult to interpret. As an alternative, we modeled and refined the changed-state only. To prevent reversion of the model into ground state density, coordinate refinement was switched off after fragments were modelled. Hydrogens were added with phenix.ready_set, waters were updated automatically and B-factors were refined anisotropically for non-hydrogen and non-water atoms. After one round of refinement, waters added into ground state electron density were removed. This was achieved by aligning the DMSO-only model to the refined model, and removing any water molecules within 2.2 Å of the DMSO-only model. A final round of refinement was performed without updating water molecules.

### P4_3_ crystals at UCSF

#### Protein expression and purification

The C2 sequence in pET22b(+) was converted into the P4_3_ sequence by removal of Glu170 and replacement of the N-terminal Asn-Ala-Gly-Glu motif with a methionine. Additionally, a Ser-Ser-Gly-Val-Asp-Leu-Gly-Thr linker was introduced between the His_6_ tag and the TEV recognition sequence **(Supplementary data set 1**). All cloning steps were performed by PCR with overlapping primers and Gibson assembly (*94*). Protein was purified using the same protocol as the C2 protein, except that after SEC, the protein was concentrated to 40 mg/ml prior to flash freezing in liquid nitrogen.

#### Crystallization

Initially, crystals were grown by hanging-drop vapour diffusion with a reservoir solution containing 34% PEG 3000 and 100 mM CHES pH 9.5. Screens were performed using pre-greased VDX plates (Hampton Research, HR3-142) with 0.5 ml reservoir solution in each well. Crystallization drops were set up on silicon coverslips (Hampton Research, HR3-233) with 2 μl Mac1 at 10 mg/ml and 2 μl reservoir. Crystals grew after 2-4 days at 19°C. As with the C2 crystals, microseeding was required to achieve consistent nucleation. Seed stocks were prepared as described previously, except the seed storage buffer used was 35% PEG 3000, 100 mM CHES pH 9.5 and 2 mM DTT. Crystals for fragment soaking were grown using SwissCI 3-well sitting drop plates with reservoirs containing 30 μl 28% PEG 3000, 100 mM CHES pH 9.5. Crystallization drops were set-up with 100 nl reservoir solution, 100 nl seed stocks (1:100,000 dilution) and 200 nl Mac1 at 40 mg/ml. Crystals were grown at 19°C and reached a maximum size after 24 hours.

#### Fragment and ADP-ribose soaking

Fragment soaks were performed using the same protocol as the C2 crystals, with soak times between 2-6 hours. ADPr soaks were performed similarly, except that ADP-ribose was prepared in water to 100 mM, and crystals were soaked with 80 nl ADPr (20 mM final concentration). Crystals were vitrified directly after soaking using a Nanuq cryocooling device (Mitegen).

#### Fragment data collection, processing, modelling and refinement

Diffraction data was collected at ALS beamline 8.3.1, SSRL beamline 12-1 and NSLS-II beamline 17-ID-2. The data collection summary is summarized in **Supplementary data set 1**. Fragment datasets were indexed, integrated and scaled using XDS (*85*) and merged with Aimless (*91*). In addition to the fragment soaks, we collected diffraction data for 40 crystals soaked only with DMSO. To generate a DMSO-only model, a single high resolution dataset was selected and phases were obtained by molecular replacement using the 0.77 Å C2 structure as a search model. Refinement and model building was performed as described previously for the C2 crystals. The fragment datasets were prepared for PanDDA analysis using the DIMPLE pipeline. Fragments were identified using PanDDA, with the background electron density map generated using 35 DMSO-only datasets (**Supplementary data set 1**). As with the analysis of C2 electron density maps, PanDDA was re-run with a decreased Z-map threshold (contour_level = 2.5, min_blob_volume = 5, min_blob_z_peak = 2.5). This strategy identified an additional 24 fragments. Fragment modeling and refinement was carried out using the same protocol as the experiment with C2 crystals.

### P4_3_ crystals at Oxford/XChem

#### Protein expression and purification

SARS-CoV-2 Nsp3 Mac1 (residues 3-169) was cloned into a pNIC28-Bsa4 expression vector which adds an N-terminal His_6_-tag and a TEV protease recognition site for removal of the tag. For expression of protein used for crystallisation, the constructs was transformed into the *E. coli* Rosetta strain BL21(DE3)-R3 and cells were grown at 37°C in LB medium (Miller) supplemented with 50 μg/ml of kanamycin and 35 μg/ml of chloramphenicol. After reaching an OD_600_ of 0.5–0.6, the temperature was lowered to 18°C prior to induction of protein expression overnight by adding 0.5 mM isopropyl 1-thio-D-galactopyranoside. Harvested cells were resuspended in lysis buffer (50 mM HEPES (pH 7.5), 500 mM NaCl, 5% glycerol, 20 mM imidazole, 10 mM βME, cOmplete EDTA-free protease inhibitors (Roche)) and stored at −20°C until purification. For protein purification, pellets were gently thawed in lukewarm water and lysed by high-pressure homogenisation. DNA was digested using Benzonase. Proteins were purified by immobilised metal affinity chromatography (IMAC) using Ni-Sepharose resin (GE Healthcare) and eluted stepwise in binding buffer containing 40–500 mM imidazole. A high salt wash with 1 M NaCl was combined with the first elution step including 40 mM imidazole. Removal of the hexahistidine tag was carried out by addition of recombinant TEV protease during overnight dialysis into buffer without imidazole, followed by purification on a second IMAC column and finally by size-exclusion chromatography (SEC) (Superdex 75, GE Healthcare) in a buffer consisting of 20 mM HEPES (pH 8.0), 250 mM NaCl and 2 mM DTT. Macrodomain protein used for HTRF assay was not subjected to TEV cleavage and purified after the IMAC step by SEC in a buffer consisting of 25 mM HEPES pH 7.4, 300 mM NaCl, 5% glycerol and 0.5 mM TCEP. Proteins were characterised by SDS-PAGE, then flash frozen in liquid nitrogen and stored at −80°C until required.

#### Crystallographic fragment screening

SARS-CoV-2 Nsp3 Mac1 was concentrated to a final concentration of 47 mg/ml and *apo* crystals were grown in crystallization solution containing 100 mM CHES pH 9.5 and 30% PEG3000. Fragments were soaked into crystals as previously described (*35*) by adding dissolved compounds directly to the crystallisation drops using an ECHO liquid handler (final concentration 10% DMSO); drops were incubated for approximately 1-3 hours prior to mounting and flash freezing in liquid nitrogen.

Data was collected at the beamline I04-1 at 100K and automatically processed with Diamond Light Source’s auto-processing pipelines using XDS (*85*) and either xia2 (*95*), autoPROC (*96*) or DIALS (*90*) with the default settings. Most Nsp3 macrodomain data processed to a resolution of approximately 1.1 Å. Further analysis was performed with XChemExplorer, (*36*), electron density maps were generated with Dimple (*97*) and ligand-binding events were identified using PanDDA (*54*). Ligands were modelled into PanDDA-calculated event maps using Coot (*98*), restraints were calculated with AceDRG (*99*) or GRADE (*100*), and structures were refined with BUSTER (*101*). Coordinates, structure factors and PanDDA event maps for the structures discussed are deposited in the Protein Data Bank. Data collection and refinement statistics are summarised in the Supplementary Data File 1.

### Molecular Docking Screens

Docking was performed against the crystal structure of SARS-CoV-2 Nsp3 Mac1 bound to ADP-ribose (PDB 6W02). Chain B and all water molecules except for HOH324, HOH344, HOH384, and HOH406 were removed. These water molecules were included in the docking template structure since they were buried within the ADP-ribose binding site and formed bridging hydrogen bonds between ADP-ribose and the protein. The protein structure in complex with ADP-ribose and the 4 selected water molecules was capped at N- and C-termini and prepared for docking following the prepwizard protocol in Maestro (Schrödinger; (*102*)). Accordingly, protons were added using Epik and protonation states were optimized with PropKa at pH 7. Finally, the structure was energetically minimized using the OPLS3e force field, thereby, the maximum heavy-atom deviation from the initial structure was 0.3 Å (*102*).

Docking was performed with DOCK3.7 using pre-calculated scoring grids for rapid evaluation of docked molecules (*103*). AMBER united atom charges (*104*) were assigned to the minimized protein structure and water molecules. Partial atomic charges of backbone amides for residues Ile23 and Phe156 were increased by 0.2 elementary charge units without changing the net charge of the residues, as described previously (*105*). The low dielectric constant of the protein environment was extended outwards from the protein surface by 1.9Å using spheres generated by SPHGEN. Electrostatic potentials at the ligand-binding pocket were calculated by numerical solution of the Poisson-Boltzmann equation using QNIFFT (*106*), scoring grids for van der Waals potentials were generated with CHEMGRID. Ligand desolvation scoring grids were calculated by SOLVMAP (*107*), thereby, the volume of the low protein dielectric was extended out 0.4 Å from the protein surface, as described previously (*55*).

Since we specifically targeted the adenosine binding site of the full ADP-ribose binding pocket, atomic coordinates of adenosine rather than the whole ADP-ribose molecule were used to generate 45 matching spheres, representing favorable positions for placing ligand atoms with docking (*103*).

As ADP-ribose was the only known ligand for the viral macrodomain when we started the docking campaign, the generated scoring grids and matching spheres were judged for their ability to place and score adenosine, adenine and ribose at the adenosine binding site of the ligand binding pocket compared to 250 property-matched decoys, generated following the DUD-EZ method (*108*). Decoys share similar physical properties as the control molecules but are topologically different, hence unlikely to ligate the binding pocket. Furthermore, an “extrema” set (*108*) of approximately 500,000 molecules including anionic, neutral and cationic compounds with molecular weights ranging from 250-350 Da was screened to ensure similar enrichments for monovalent anions and neutral molecules. We note that the lack of experimentally confirmed ligands for the macrodomain did not allow exhaustive control calculations.

Virtual compound libraries were downloaded from ZINC15 (www.zinc15.docking.org) (*53*). From the set of 722,963 in-stock fragments, 696,092 compounds were successfully docked, exploring on average 2,355 orientations and 63 conformations per compound in the binding pocket. Roughly 58 billion complexes were sampled in 88 core hours, or roughly 10 minutes on a 500 core cluster. Screening the entire 20 million ZINC15 fragment library resulted in the evaluation of ca. 4.4 trillion complexes within 2,342 core hours, or 4.7 hours on 500 cores. In that screen, 19,130,798 compounds were scored and sampled in ca. 2,145 orientations and 180 conformations each. From the relatively small “in-human” library, containing 20,726 molecules, 17,362 compounds were scored, and sampling was increased to roughly 16,615 orientations per compound. 84 billion complexes were evaluated within 27 core hours.

Compounds with DOCK scores < −20 (top 500,000 compounds from the entire fragment screen), were subsequently filtered for those with strained conformations, and inspected for their ability to form hydrogen bonds to residues Asp22, Ile23, Gly48, Val49, Gly130 or Phe156. Compounds with unsatisfied hydrogen bond donors or more than three unsatisfied hydrogen bond acceptors were deprioritized. From both fragment screens, 17 in-stock compounds (8 selected from the ZINC15 in-stock library docking screen) were purchased, and 45 make-on-demand fragments were ordered of which 33 were successfully synthesized, both from Enamine. The following compounds were selected from the “in-human” collection docking screen and purchased from different vendors: Pterin (Sigma-Aldrich, P1132), Verdiperstat (MedChem Express, HY-17646), Kinetin (Cayman Chemical, 20712), Irsogladine (Cayman Chemical, 30223), Diaveridine (Cayman Chemical, 29427), N6-Benzyladenine (Cayman Chemical, 21711), PP2 (Cayman Chemical, 13198), Temozolomide (Cayman Chemical, 14163), Chrysophanol (Cayman Chemical, 19870), Isoxanthopterin (Cayman Chemical, 17564).

### Fragment linking and merging

Fragment mergers and linkers were generated using Fragmenstein (https://github.com/matteoferla/Fragmenstein) (*109*), a python module that automatically joins inspiration hits or places compounds based on inspiration hits in way that is as faithful to the positions of the inspiration hits as possible in a conformation that is energy acceptable. For merging, using RDKit (*110*) rings are temporarily collapsed into pseudo-atoms, one-to-one spatial overlapping atoms are identified, pseudo-atoms expanded with appropriate bonds to nearby atoms and various chemical corrections applied. For the constrained energy minimisation, Pyrosetta is used (*111*). Interactive online summary of mergers was made in https://michelanglo.sgc.ox.ac.uk (*112, 113*).

### Purity and structure determination of initial fragment samples ZINC901381520, ZINC82473428 and ZINC89254160 provided by Enamine

Samples of ZINC901391520, ZINC82473428 and ZINC89254160 obtained from Enamine were expected to be N_9_-alkylated isomers but electron density of compounds in X-ray co-crystal structures indicated these samples were N_3_-alkylated isomers instead (ZINC901391520_N3, ZINC82473428_N3 and ZINC89254160_N3, see Fig. S7I-L). The original samples of ZINC901391520, ZINC82473428 and ZINC89254160 used in fragment screening by X-ray crystallography were analyzed by HPLC-MS and 1H NMR to confirm sample purity and corroborate structure.

There is no reported characterization data to be used as reference for structure confirmation for N_9_- or N_3_-alkylated compounds ZINC901391520 and ZINC89254160. The N_9_-alkylated structure ZINC82473428 is a previously prepared compound with tabulated NMR data reported by Rad *et al.* (*114*).

A re-supplied sample of ZINC901391520 from a new batch synthesized at Enamine was confirmed by ^1^H NMR to be >95% purity and a different isomer than the original sample of ZINC901391520. The X-ray crystal structure of this fragment in complex with Mac1 revealed the fragment to be N_9_-alkylated isomer (**Fig. S7**I).

Original samples of ZINC901391520, ZINC82473428 and ZINC89254160 from Enamine used in fragment screen were evaluated for purity by HPLC on an Agilent 1200 Binary SL system with diode array detection and mass spectrometric detection on an Agilent 6135B Quadrupole system in electrospray ionization mode (positive ion detection). One of two HPLC Methods A or B were used to determine sample purity using mobile phase linear gradients of acetonitrile with 0.1% TFA in water with 0.1% TFA detailed below at 1.000 ml/min flow rate through a Phenomenex Gemini 3 mm C18 110 Å LC column (4.6 mm dia. x 150 mm length).

#### HPLC Method A mobile phase gradient

Gradient time points (minutes): 1.0-1.5-10.5-11.0-12.5-13.0-15.0; % acetonitrile at gradient time points: 5-5-20-95-95-5-5

#### HPLC Method B mobile phase gradient

Gradient time points (minutes): 1.0-7.0-8.0-10.0-10.5-12.0; % acetonitrile at gradient time points: 5-30-95-95-5-5

#### NMR Experiments for Samples ZINC901391520, ZINC82473428 and ZINC89254160

Original samples of ZINC901391520, ZINC82473428 and ZINC89254160 from Enamine used in the fragment screen were dissolved in *d*_*6*_-DMSO and analyzed by ^1^H and ^13^C NMR on a Bruker 400 MHz instrument with Avance III electronics. Data was obtained at ambient temperature (ca. 25°C) collecting 64 scans for proton experiments and 1024 scans for carbon experiments. Raw data was processed and reports created using ACD Spectrus software.

#### Original sample ZINC901391520

A sample of 5.5 mg ZINC901391520 was dissolved in 0.75 ml *d*_*6*_-DMSO for NMR analysis and from this solution 50 μl was diluted in 0.45 ml acetonitrile to make up the analytical sample for HPLC-MS using HPLC Method A. The sample chromatogram from HPLC revealed a single peak with UV absorbance at both 214 and 254 nm at *t*_*R*_ = 5.272 minutes. Aside from a very strong UV_214_ peak at *t*_*R*_ = 2.00 minutes attributed to DMSO co-solvent in the sample, no other peaks were observed at these UV wavelengths and sample purity estimated >98% based on UV peak area. ^1^H NMR (400 MHz, *d*_*6*_-DMSO, 25 °C) δ ppm 8.49 (s, 1H), 7.91-8.26 (br d, 2H), 7.76 (s, 1H), 6.30 (s, 1H), 5.63 (s, 2H), 3.86 (s, 3H). ^13^C NMR (101 MHz, *d*_*6*_-DMSO, 25 °C) δ ppm 172.05, 167.57, 155.01, 152.46, 149.39, 143.55, 120.18, 94.85, 57.13, 44.32. LRMS (ESI^+^) for peak at *t*_*R*_ = 5.272 minutes: observed *m/z* = 247.3 [MH]^+^ for C_10_H_10_N_6_O_2_ exact mass = 246.09.

#### Second batch sample ZINC901391520

A sample was dissolved in 0.75 ml *d*_*6*_-DMSO for NMR analysis. ^1^H NMR (400 MHz, *d*_*6*_-DMSO, 25 °C) δ ppm 8.24 (s, 1H), 8.16 (s, 1H), 7.31 (br s, 2H), 6.22 (s, 1H), 5.49 (s, 2H), 3.86 (s, 3H).

#### Sample ZINC82473428

A sample of 3.9 mg ZINC82473428 was dissolved in 0.75 ml *d*_*6*_-DMSO for NMR analysis and from this solution 50 μl was diluted in 0.45 ml acetonitrile to make up the analytical sample for HPLC-MS using HPLC Method B. The sample chromatogram from HPLC revealed a single peak with UV absorbance at both 214 and 254 nm at *t*_*R*_ = 3.766 minutes. Aside from a very strong UV_214_ peak at *t*_*R*_ = 2.00 minutes attributed to DMSO cosolvent in the sample no other peaks were observed at these UV wavelengths and sample purity estimated >98% based on UV peak area. ^1^H NMR (400 MHz, *d*_*6*_-DMSO, 25 °C) δ ppm 8.31 (s, 1H), 8.01 (br s, 2H), 7.86 (s, 1H), 4.44 (dd, *J*=13.18, 3.39 Hz, 1H), 4.31-4.40 (m, 1H), 4.20-4.30 (m, 1H), 3.75-3.87 (m, 1H), 3.58-3.70 (m, 1H), 1.93-2.07 (m, 1H), 1.75-1.92 (m, 2H), 1.58-1.73 (m, 1H). ^13^C NMR (101 MHz, *d*_*6*_-DMSO, 25 °C) d ppm 154.78, 151.53, 149.56, 144.30, 75.35, 67.24, 52.54, 40.44, 28.23, 25.03. LRMS (ESI^+^) for peak at *t*_*R*_ = 3.766 minutes: observed *m/z* = 220.3 [MH]^+^ for C10H13N5O exact mass = 219.11.

Reported NMR data for compound ZINC82473428_N9 from Rad et al., 2015 (*114*): ^1^H NMR (400 MHz, *d*_*6*_-DMSO, 25 °C) δ ppm 7.91 (s, 1H), 7.83 (s, 1H), 7.01 (br s, 2H), 3.87-3.99 (m, 3H), 3.34-3.52 (m, 2H), 1.30-1.54 (complex m, 4H). ^13^C NMR (101 MHz, *d*_*6*_-DMSO, 25 °C) δ ppm 156.6, 152.9, 149.2, 144.7, 117.2, 80.6, 67.9, 57.8, 29.1, 25.1.

#### Sample ZINC89254160

A sample of 3.2 mg ZINC89254160 was dissolved in 0.75 ml *d*_*6*_-DMSO for NMR analysis and from this solution 50 μl was diluted in 0.45 ml acetonitrile to make up the analytical sample for HPLC-MS using HPLC Method A. The sample chromatogram from HPLC revealed a major peak and a minor peak with UV absorbances at both 214 and 254 nm: major peak *t*_*R*_ = 6.530 minutes and minor peak *t*_*R*_ = 6.751 minutes. Relative peak area calculated as percentage of combined UV peak area at 254 nm was 93.3% major peak and 6.7% minor peak (corresponds to ca. 14:1 ratio). Aside from a very strong UV_214_ peak at *t*_*R*_ = 2.00 minutes attributed to DMSO cosolvent in the sample no other peaks were observed at these UV wavelengths. Tabulated NMR data reported here for major peaks only. ^1^H NMR (400 MHz, *d*_*6*_-DMSO, 25 °C) δ ppm 8.47 (s, 1H), 7.95 (br s, 2H), 7.73 (s, 1H), 7.47 (s, 1H), 5.55 (s, 2H), 2.60 (s, 3H). ^13^C NMR (101 MHz, *d*_*6*_-DMSO, 25 °C) δ ppm 166.21, 154.93, 152.47, 149.63, 149.51, 143.63, 120.43, 117.69, 48.08, 18.66. LRMS (ESI^+^) for major peak at *t*_*R*_ = 6.530 minutes: observed *m/z* = 247.3 [MH]^+^ for C10H10N6S exact mass = 246.07. LRMS (ESI^+^) for minor peak at *t*_*R*_ = 6.751 minutes: observed *m/z* = 247.3 [MH]^+^ for C_10_H_10_N_6_S exact mass = 246.07. Major peak and minor peak have the same observed mass peak in LRMS and are presumed to be different N-alkylated isomers.

#### Conclusions Based on HPLC-MS and NMR Characterization of Samples ZINC901391520, ZINC82473428 and ZINC89254160

HPLC-MS data confirmed that samples ZINC901391520 and ZINC82473428 are single compounds >98% purity with mass peak corresponding to either N_9_- or N_3_-alkylated isomers. Both ^1^H and ^13^C NMR data corroborated initial samples ZINC901391520 and ZINC82473428 are >98% single compound. The very high purity determined for these two samples strongly rules out the possibility that X-ray co-crystals obtained were the result of protein complexed to trace amount of alternative isomer in the samples. For ZINC89254160, HPLC-MS data confirmed that there was a 13:1 ratio of isomers in this sample and it is possible the X-ray co-crystal obtained with ZINC89254160 was the result of protein complexed to trace/minor amounts of the alternative isomer (N_3_-alkylated).

The NMR data obtained for sample ZINC82473428 used in crystallographic fragment screen does not match NMR data reported in the literature for the N_9_-alkylated ZINC82473428 and thus this sample is presumed not to be N_9_-alkylated isomer. NMR data is not sufficient to unambiguously assign N_3_- or N_9_-alkylated structures for ZINC901391520, ZINC82473428 or ZINC89254160 and the unambiguous structure assignment of ZINC901391520, ZINC82473428 and ZINC89254160 as N_3_-alkylated isomers in this work was provided by the electron density observed for these fragments in X-ray co-crystal structures obtained. The crystal structure with ZINC400552187 additionally revealed the N_3_-alkylated structure instead of the requested N_9_-alkylated form. Using DSF and ITC, ZINC901391520, ZINC82473428, ZINC89254160, ZINC400552187 were initially screened as the N_3_-alkylated isomer (ZINC901391520_N3 (PDB 5RSK), ZINC82473428_N3 (PDB 5RVF), ZINC89254160_N3 (PDB 5RSJ), ZINC400552187_N3 (PDB 5RVG)). In addition the N_9_-alkylated ZINC901391520 (PDB 5S6W) was tested in DSF and the peptide-competition assay (HTRF) (**Supplementary Table 1**).

### Differential Scanning Fluorimetry (DSF)

#### Compound handling

Compounds were dissolved in DMSO to a final concentration of 100 mM, and placed in a 384-well Echo source plate (Labcyte PP0200). Using a LabCyte Echo. Each compound was dispensed into a 384-well storage plate (Greiner BioOne 781280) in five stock concentrations in two-fold serial dilutions (compounds: 6.25-100 mM; ADP-ribose: 0.625-10 mM) and a final volume of 750 nL in triplicate. Two identical plates were created, with the second plate used to provide protein-free controls for all tested conditions. Echo dispensing instructions were created by an in-house script (https://gestwickilab.shinyapps.io/echo_layout_maker/).

#### Experimental set-up

DSF buffer was prepared by adding 10 μl of SYPRO Orange (Thermo Scientific, Product #S6650) to 10 ml buffer (50 mM Tris HCl pH 7.50, 150 mM NaCl, 1 mM EDTA, 1 mM DTT, 0.01% Triton X-100), for a final dye concentration of 5X (10 μM) SYPRO Orange. A compound plate (see above) was resuspended by the addition of 20 μl of DSF buffer, and set aside for 20 minutes in the dark. Purified Nsp3 Mac1 domain was diluted to 10 μM in DSF buffer, and 2 μl of either protein solution or protein-free buffer was added to each well a 384-well white PCR plate (Axygen PCR-384-LC480WNFBC, Lot #23517000) using an E100 ClipTip p125 Matrix Pipette. 8 μl of resuspended compound was transferred to each well of the protein- and buffer-containing PCR plate using an Opentrons OT-2 liquid handling system, yielding the following final conditions: 2 μM Nsp3 Mac1, 5X (10 μM) SYPRO Orange, 3% DMSO, 0.1-3 mM fragments, and 0.1-1 mM ADP-ribose. The PCR plate was spun briefly in a salad spinner to remove bubbles, and sealed with optically clear film (Applied Biosystems, MicroAmp Optical Adhesive Film, Product #4311971). In an Analytik Jena qTower 384G qPCR instrument, plate was continuously heated from 25 - 94ºC at a rate of 1ºC/minute, and fluorescence was measured at each degree in the TAMRA channel (535 nm / 580 nm). 53 of 54 fragments could be tested up to 3 mM without assay interference in these conditions (**Supplementary data set 2, Supplementary Table 1**).

Raw DSF data for the Nsp3 Mac1 construct used in this work was characterized by a major transition at 50.8 +/− 0.3ºC, with a minor second transition at 67.0 +/− 3.6ºC (**Fig. 9**, **Supplementary data set 2, Supplementary Table 1**); results described refer to the major transition. Significance was defined as compounds with ANOVA p-values < 0.005 for Tm_a_ over the tested concentration regime.

### Isothermal Titration Calorimetry (ITC)

All ITC titrations were performed on a MicroCal iTC 200 instrument (GE Healthcare, IL). All reactions were performed in 20 mM Tris pH 7.5, 150 mM NaCl using 300 - 600 μM of Mac1 at 25°C. Titration of 4 mM ADP-ribose (Sigma-Aldrich A0752) or 4-10 mM fragment contained in the stirring syringe included a single 0.2 μl injection, followed by 18 consecutive injections of 2 μl. Data analysis of thermograms was analyzed using one set of binding sites model of the <u>RITC package</u> to obtain all fitting model parameters for the experiments.

### Homogeneous Time-Resolved Fluorescence (HTRF)-based Peptide Displacement Assay

Fragment inhibitory activity on Mac1 was assessed by the displacement of an ADP-ribose-conjugated biotin peptide from the His_6_-tagged Nsp3 Mac1 domain using HTRF with a Eu^3+^-conjugated anti-His_6_ antibody donor and streptavidin-conjugated acceptor. Compounds were dispensed into ProxiPlate-384 Plus (PerkinElmer) assay plates using an Echo 525 Liquid Handler (Labcyte). Binding assays were conducted in a final volume of 16 μl with 12.5 nM Nsp3 Mac1 domain, 400 nM peptide ARTK(Bio)QTARK(Aoa-RADP)S, 1:125 Streptavidin-XL665, 1:20000 Anti-His_6_-Eu^3+^ cryptate in assay buffer (25 mM HEPES pH7.0, 20 mM NaCl, 0.05% BSA, 0.05% Tween20). Assay reagents were dispensed into plates using a Multidrop combi (Thermo Scientific) and incubated at room temperature for 1 h. Fluorescence was measured using a PHERAstar microplate reader (BMG) using the HTRF module with dual emission protocol (A = excitation of 320 nm, emission of 665 nm, and B = excitation of 320 nm, emission of 620 nm). Raw data were processed to give an HTRF ratio (channel A/B × 10,000), which was used to generate IC_50_ curves by nonlinear regression using GraphPad Prism v8 (GraphPad Software, CA, USA).

## Supporting information

Supplementary Data Set 1

Supplementary Data Set 2

## ACKNOWLEDGMENTS

J. Fraser is supported by NIH GM123159, NSF Rapid 2031205, and a TMC Award from the UCSF Program for Breakthrough Biomedical Research, funded in part by the Sandler Foundation. Work in the Ahel laboratory is funded by the Wellcome Trust (grants 101794 and 210634), BBSRC (BB/R007195/1) and Cancer Research UK (C35050/A22284). BKS is supported by NIH R35GM122481 and DARPA HR0011-19-2-0020. The crystallographic screen at Oxford was supported by the XChem facility at Diamond Light Source (proposal ID MX27001). We thank all the staff of Diamond Light Source for providing support and encouragement which allowed us to carry out this work during the COVID-19 lockdown. We also thank Gemma Davison, Selma Dormen, James Sanderson, Matthew Martin, Mike Waring and Martin Noble (CRUK Newcastle Drug Discovery Unit, Newcastle University), Thomas Downes, Paul Jones, Hanna Klein, James Firth and Peter O’Brien (York University), David Bajusz and Gyorgy Keseru (Hungarian Academy of Sciences) for providing fragment libraries. We also acknowledge EU-OPENSCREEN ERIC for providing its fragment library for the presented scientific work. EU-OPENSCREEN ERIC has received funding from European Union’s Horizon 2020 research and innovation programme under grant agreement No 823893 (EU-OPENSCREEN-DRIVE).The SGC is a registered charity (number 1097737) that receives funds from AbbVie, Bayer Pharma AG, Boehringer Ingelheim, Canada Foundation for Innovation, Eshelman Institute for Innovation, Genome Canada, Innovative Medicines Initiative (EU/EFPIA) [ULTRA-DD grant no. 115766], Janssen, Merck KGaA Darmstadt Germany, MSD, Novartis Pharma AG, Ontario Ministry of Economic Development and Innovation, Pfizer, São Paulo Research Foundation-FAPESP, Takeda, and Wellcome [106169/ZZ14/Z]. R. Díaz and T. Wu were supported by NSF GRFP. R. Díaz is a Howard Hughes Medical Institute Gilliam Fellow. I. Young was supported by NIH F32GM133129. M. Ferla is supported by the Wellcome Trust 203141/Z/16/Z and the NIHR Biomedical Research Centre Oxford

## QCRG

The QCRG Structural Biology Consortium has received support from: Quantitative Biosciences Institute, Defense Advanced Research Projects Agency HR0011-19-2-0020 (to D.A.A. and K.A.V.; B. Shoichet PI), FastGrants COVID19 grant (K.A. Verba PI), Laboratory For Genomics Research (O.S. Rosenberg PI) and Laboratory for Genomics Research LGR-ERA (R.M. Stroud PI).

## Beamlines

Beamline 8.3.1 at the Advanced Light Source is operated by the University of California Office of the President, Multicampus Research Programs and Initiatives grant MR-15-328599, NIH (R01 GM124149 and P30 GM124169), Plexxikon Inc., and the Integrated Diffraction Analysis Technologies program of the US Department of Energy Office of Biological and Environmental Research. The Advanced Light Source (Berkeley, CA) is a national user facility operated by Lawrence Berkeley National Laboratory on behalf of the US Department of Energy under contract number DE-AC02-05CH11231, Office of Basic Energy Sciences.

Use of the Stanford Synchrotron Radiation Lightsource, SLAC National Accelerator Laboratory, is supported by the U.S. Department of Energy, Office of Science, Office of Basic Energy Sciences under Contract No. DE-AC02-76SF00515. The SSRL Structural Molecular Biology Program is supported by the DOE Office of Biological and Environmental Research, and by the National Institutes of Health, National Institute of General Medical Sciences (including P41GM103393). Extraordinary SSRL operations were supported in part by the DOE Office of Science through the National Virtual Biotechnology Laboratory, a consortium of DOE national laboratories focused on response to COVID-19, with funding provided by the Coronavirus CARES Act.

This research used beamline 17-ID-2 of the National Synchrotron Light Source II, a U.S. Department of Energy (DOE) Office of Science User Facility operated for the DOE Office of Science by Brookhaven National Laboratory under Contract No. DE-SC0012704. The Center for BioMolecular Structure (CBMS) is primarily supported by the National Institutes of Health, National Institute of General Medical Sciences (NIGMS) through a Center Core P30 Grant (P30GM133893), and by the DOE Office of Biological and Environmental Research (KP1605010).

## Contributions

M. Schuller designed and cloned the construct that yielded the P4_3_ crystals at Oxford/XChem; expressed, purified and establishing crystallization conditions for P4_3_ crystals for Oxford/XChem; assisted with data processing/analysis at XChem; set up the HTRF functional assay and performed data interpretation; prepared the manuscript. G. Correy cloned, expressed and purified the P4_3_ construct at UCSF; crystallized, performed fragment soaking, vitrified crystal, collected X-ray diffraction data and processed data for fragment screens at UCSF; modelled, refined and analyzed fragment structures at UCSF; purified, methylated and crystalized the C2 construct; performed ADP-ribose soaks; refined the apo and ADP-ribose bound structures determined using the P4_3_ crystals; prepared the manuscript. S. Gahbauer performed docking screens against Mac1; performed the chemoinformatic analysis of fragment libraries; assisted with fragment-linking and - merging; prepared the manuscript. D. Fearon crystallized, prepared samples, collected X-ray diffraction data, refined and analyzed fragment structures at XChem; prepared the manuscript. T. Wu performed and analyzed DSF experiments; prepared the manuscript. R. Díaz designed the construct that yielded the C2 crystals; expressed the C2 construct; performed and analyzed ITC experiments; prepared the manuscript. I. Young collected X-ray diffraction data and processed the diffraction data for crystals screened at UCSF. L. Martins assisted with docking screens against Mac1. D. Smith assisted with DSF experiments. U. Schulze-Gahmen crystallized the C2 construct at UCSF. T. Owens purified the C2 construct at UCSF. I. Deshpande collected X-ray diffraction data for C2 crystals at UCSF. G. Merz purified the C2 construct at UCSF. A. Thwin purified the C2 construct at UCSF. J. Biel supported the fragment soaking experiments performed at UCSF; supported the fragment modeling and refinement at UCSF. J. Peters purified C2 construct at UCSF. M. Moritz purified C2 construct at UCSF. N. Herrera supported crystallization of the C2 construct at UCSF. H. Kratochvil supported crystallization of the C2 construct at UCSF. QCRG Structural Biology Consortium provided infrastructure and support for experiments performed at UCSF. A. Aimon prepared samples and collected X-ray diffraction data at XChem. J. Bennett set up the HTRF functional assay; analyzed and collected data for the HTRF assay. J. Neto collected X-ray diffraction data at XChem. A. Cohen supported X-ray diffraction experiments at the SSRL. A. Dias collected X-ray diffraction data at XChem. A. Douangamath crystallized, prepared samples, collected X-ray diffraction data, refined and analyzed fragment structures at XChem. L. Dunnett collected X-ray diffraction data at XChem. O. Fedorov set up the HTRF functional assay; analyzed and collected data for the HTRF assay. M. Ferla performed fragment merging and linking. M. Fuchs supported X-ray diffraction experiments at the NSLS-II. T. Gorrie-Stone deposited fragments structures at XChem. J. Holton supported X-ray diffraction experiments at the ALS. M. Johnson analyzed and assigned structures to the N_3_- and N_9_-alkylated fragments. T. Krojer analyzed structural data at XChem. G. Meigs supported X-ray diffraction experiments at the ALS. A. Powell collected X-ray diffraction data at XChem. J. Rack provided feedback on the manuscript. V. Rangel refined fragment structures at XChem. S. Russi supported X-ray diffraction experiments at the SSRL. R. Skyner deposited fragments structures at XChem. C. Smith supported X-ray diffraction experiments at the SSRL. A. Soares supported X-ray diffraction experiments at the NSLS-II. J. Wierman supported X-ray diffraction experiments at the SSRL. K. Zhu expressed and crystallized the P4_3_ construct at XChem. N. Jura supervised work. A. Ashworth supervised work; prepared the manuscript. J. Irwin designed the UCSF_91 fragment library; prepared the manuscript. M. Thompson vitrified C2 crystals at UCSF; collected X-ray diffraction data for C2 crystals at UCSF; refined the ultra-high-resolution structure determined using C2 crystals; prepared the manuscript. J. Gestwicki supervised work. F. Delft provided the XChem facility; prepared the manuscript; supervised work. B. Shoichet guided and evaluated the docking work; prepared the manuscript; supervised work. J. Fraser supervised work; prepared the manuscript; arranged funding. I. Ahel supervised work; prepared the manuscript; arranged funding.

## QCRG Structural Biology Consortium

This work was supported by the QCRG (Quantitative Biosciences Institute Coronavirus Research Group) Structural Biology Consortium. Listed below are the contributing members of the consortium listed by teams. Within each team the team leads are italicized (responsible for organization of each team, and for the experimental design utilized within each team), then the rest of team members are listed alphabetically.

### Bacterial expression team

*Amy Diallo, Meghna Gupta, Erron W. Titus*, Jen Chen, Loan Doan, Sebastian Flores, Mingliang Jin, Huong T. Kratochvil, Victor L. Lam, Yang Li, Megan Lo, Gregory E. Merz, Joana Paulino, Aye C. Thwin, Zanlin Yu, Fengbo Zhou, Yang Zhang.

### Protein purification team

*Daniel Asarnow, Michelle Moritz, Tristan W. Owens, Sergei Pourmal*, Caleigh M. Azumaya, Cynthia M. Chio, Bryan Faust, Meghna Gupta, Kate Kim, Joana Paulino, Jessica K. Peters, Kaitlin Schaefer, Tsz Kin Martin Tsui.

### Crystallography team

*Nadia Herrera, Huong T. Kratochvil, Ursula Schulze-Gahmen, Iris D. Young*, Justin Biel, Ishan Deshpande, Xi Liu.

#### CryoEM grid freezing/collection team

*Caleigh M. Azumaya, Axel F. Brilot, Gregory E. Merz, Cristina Puchades, Alexandrea N. Rizo, Ming Sun*, Julian R. Braxton, Meghna Gupta, Fei Li, Kyle E. Lopez, Arthur Melo, Gregory E. Merz, Frank Moss, Joana Paulino, Thomas H. Pospiech, Jr., Sergei Pourmal, Amber M. Smith, Paul V. Thomas, Feng Wang, Zanlin Yu.

### CryoEM data processing team

*Axel F. Brilot, Miles Sasha Dickinson, Gregory E. Merz, Henry C. Nguyen, Alexandrea N. Rizo,* Daniel Asarnow, Julian R. Braxton, Melody G. Campbell, Cynthia M. Chio, Un Seng Chio, Devan Diwanji, Bryan Faust, Meghna Gupta, Nick Hoppe, Mingliang Jin, Fei Li, Junrui Li, Yanxin Liu, Joana Paulino, Thomas H. Pospiech, Jr., Sergei Pourmal, Smriti Sangwan, Raphael Trenker, Donovan Trinidad, Eric Tse, Kaihua Zhang, Fengbo Zhou.

### Mammalian cell expression team

*Christian Billesboelle, Melody G. Campbell, Devan Diwanji, Carlos Nowotny, Amber M. Smith, Jianhua Zhao*, Caleigh M. Azumaya, Alisa Bowen, Nick Hoppe, Yen-Li Li, Phuong Nguyen, Cristina Puchades, Mali Safari, Smriti Sangwan, Kaitlin Schaefer, Raphael Trenker, Tsz Kin Martin Tsui, Natalie Whitis.

### Infrastructure team

David Bulkley, Arceli Joves, Almarie Joves, Liam McKay, Mariano Tabios, Eric Tse. **Leadership team:***Oren S Rosenberg, Kliment A Verba*, David A Agard, Yifan Cheng, James S Fraser, Adam Frost, Natalia Jura, Tanja Kortemme, Nevan J Krogan, Aashish Manglik, Daniel R. Southworth, Robert M Stroud.

## DATA AND MATERIALS AVAILIBILITY

All data generated or analyzed during this study are included in this article and its Supplementary Information. Crystallographic coordinates and structure factors for all structures have been deposited in the Protein Data Bank with the following accessing codes: 7KR0, 7KR1, 7KQW, 7KQO, 7KQP, 5RVJ, 5RVK, 5RVL, 5RVM, 5RVN, 5RVO, 5RVP, 5RVQ, 5RVR, 5RVS, 5RVT, 5RVU, 5RVV, 5RS7, 5RS8, 5RS9, 5RSB, 5RSC, 5RSD, 5RSE, 5RSF, 5RSG, 5RSH, 5RSI, 5RSJ, 5RSK, 5RSL, 5RSM, 5RSN, 5RSO, 5RSP, 5RSQ, 5RSR, 5RSS, 5RST, 5RSU, 5RSV, 5RSW, 5RSX, 5RSY, 5RSZ, 5RT0, 5RT1, 5RT2, 5RT3, 5RT4, 5RT5, 5RT6, 5RT7, 5RT8, 5RT9, 5RTA, 5RTB, 5RTC, 5RTD, 5RTE, 5RTF, 5RTG, 5RTH, 5RTI, 5RTJ, 5RTK, 5RTL, 5RTM, 5RTN, 5RTO, 5RTP, 5RTQ, 5RTR, 5RTS, 5RTT, 5RTU, 5RTV, 5RTW, 5RTX, 5RTY, 5RTZ, 5RU0, 5RU1, 5RU2, 5RU3, 5RU4, 5RU5, 5RU6, 5RU7, 5RU8, 5RU9, 5RUA, 5RUC, 5RUD, 5RUE, 5RUF, 5RUG, 5RUH, 5RUI, 5RUJ, 5RUK, 5RUL, 5RUM, 5RUN, 5RUO, 5RUP, 5RUQ, 5RUR, 5RUS, 5RUT, 5RUU, 5RUV, 5RUW, 5RUX, 5RUY, 5RUZ, 5RV0, 5RV1, 5RV2, 5RV3, 5RV4, 5RV5, 5RV6, 5RV7, 5RV8, 5RV9, 5RVA, 5RVB, 5RVC, 5RVD, 5RVE, 5RVF, 5RVG, 5RVH, 5RVI, 5S6W, 5S18, 5S1A, 5S1C, 5S1E, 5S1G, 5S1I, 5S1K, 5S1M, 5S1O, 5S1Q, 5S1S, 5S1U, 5S1W, 5S1Y, 5S20, 5S22, 5S24, 5S26, 5S27, 5S28, 5S29, 5S2A, 5S2B, 5S2C, 5S2D, 5S2E, 5S2F, 5S2G, 5S2H, 5S2I, 5S2J, 5S2K, 5S2L, 5S2M, 5S2N, 5S2O, 5S2P, 5S2Q, 5S2R, 5S2S, 5S2T, 5S2U, 5S2V, 5S2W, 5S2X, 5S2Y, 5S2Z, 5S30, 5S31, 5S32, 5S33, 5S34, 5S35, 5S36, 5S37, 5S38, 5S39, 5S3A, 5S3B, 5S3C, 5S3D, 5S3E, 5S3F, 5S3G, 5S3H, 5S3I, 5S3J, 5S3K, 5S3L, 5S3M, 5S3N, 5S3O, 5S3P, 5S3Q, 5S3R, 5S3S, 5S3T, 5S3U, 5S3V, 5S3W, 5S3X, 5S3Y, 5S3Z, 5S40, 5S41, 5S42, 5S43, 5S44, 5S45, 5S46, 5S47, 5S48, 5S49, 5S4A, 5S4B, 5S4C, 5S4D, 5S4E, 5S4F, 5S4G, 5S4H, 5S4I, 5S4J, 5S4K.

