## Supplementary Information for "Fragment Binding to the Nsp3 Macrodomain of SARS-CoV-2 Identified Through Crystallographic Screening and Computational Docking"

**Affiliations**
[1] Sir William Dunn School of Pathology, University of Oxford, South Parks Road, Oxford OX1 3RE, UK
[2] Department of Bioengineering and Therapeutic Sciences, University of California San Francisco, CA, USA
[3] Department of Pharmaceutical Chemistry, University of California San Francisco San Francisco, CA, USA
[4] Diamond Light Source Ltd., Harwell Science and Innovation Campus, Didcot OX11 0DE, United Kingdom
[5] Institute for Neurodegenerative Disease, University of California San Francisco, CA, USA
[6] Chemistry and Chemical Biology Graduate Program, University of California San Francisco, CA, USA
[7] Tetrad Graduate Program, University of California San Francisco, CA, USA
[8] Quantitative Biosciences Institute (QBI) Coronavirus Research Group Structural Biology Consortium, University of California San Francisco, CA, USA
[9] Biochemistry Department, Institute for Biological Sciences, Federal University of Minas Gerais. Belo Horizonte, Brazil
[10] Helen Diller Family Comprehensive Cancer, University of California San Francisco, CA, USA
[11] Centre for Medicines Discovery, University of Oxford, South Parks Road, Headington, OX3 7DQ, UK
[12] Stanford Synchrotron Radiation Lightsource, SLAC National Accelerator Center, Menlo Park, CA 94025, USA
[13] Wellcome Centre for Human Genetics, University of Oxford, Old Road Campus, Oxford OX3 7BN, UK
[14] National Synchrotron Light Source II, Brookhaven National Laboratory, Upton, NY, USA
[15] Department of Biochemistry and Biophysics, University of California San Francisco, CA, USA
[16] Department of Molecular Biophysics and Integrated Bioimaging, Lawrence Berkeley National Laboratory, Berkeley, CA, USA
[17] ChemPartner Corporation, South San Francisco, CA, USA
[18] Structural Genomics Consortium, University of Oxford, Old Road Campus, Roosevelt Drive, Headington OX3 7DQ, UK
[19] School of Pharmaceutical Sciences of Ribeirao Preto, University of Sao Paulo, São Paulo, Brazil
[20] Photon Sciences, Brookhaven National Laboratory, Upton, NY, USA
[21] Department of Cellular and Molecular Pharmacology, University of California San Francisco, CA, USA
[22] Department of Chemistry and Chemical Biology, University of California Merced, CA, USA
[23] Department of Biochemistry, University of Johannesburg, Auckland Park, 2006, South Africa

† These authors contributed equally
‡ Full listing at the end of the manuscript

### SUPPLEMENTARY FIGURES

**Supplementary Figure 1.** Ultra-high resolution features in Mac1 electron density maps.

**Supplementary Figure 2.** Comparison of isomorphism and DMSO tolerance of the C2 and $P4_3$ crystals.

**Supplementary Figure 3.** Crystal packing determines active site accessibility.

**Supplementary Figure 4.** Structure and sequence comparison of Mac1 with related viral and human macrodomains.

**Supplementary Figure 5.** Physical properties, scaffold and chemotype analysis of screened fragment libraries.

**Supplementary Figure 6.** Overview of fragment binding to protomer A and protomer B of the $P4_3$ crystals.

**Supplementary Figure 7.** Additional soaking hits from docking and adenine-N3 vs -N9-alkylated isomers.

**Supplementary Figure 8.** Mac1 subsites compared to the adenine binding site in kinases and the oxyanion binding site in carboxylesterases.

**Supplementary Figure 9.** Comparison of DSF, HTRF, and ITC results for compounds tested in all assays.

### SUPPLEMENTARY TABLE

**Supplementary Table 1**. Characterisation of docking and crystallographic hits including docking scores and solution binding data (spreadsheet, **SI_table1**)

### SUPPLEMENTARY DATA SETS

**Supplementary data set 1.** Crystallographic fragment screening information including sequences, data collection strategy, data reduction and refinement stats, summary of all fragment datasets, subsite classification (spreadsheet, **SI_dataset_1**)

**Supplementary data set 2.** Selected fragment structures. DSF, ITC, and peptide displacement data for all tested compounds (PDF, **SI_dataset_2**).

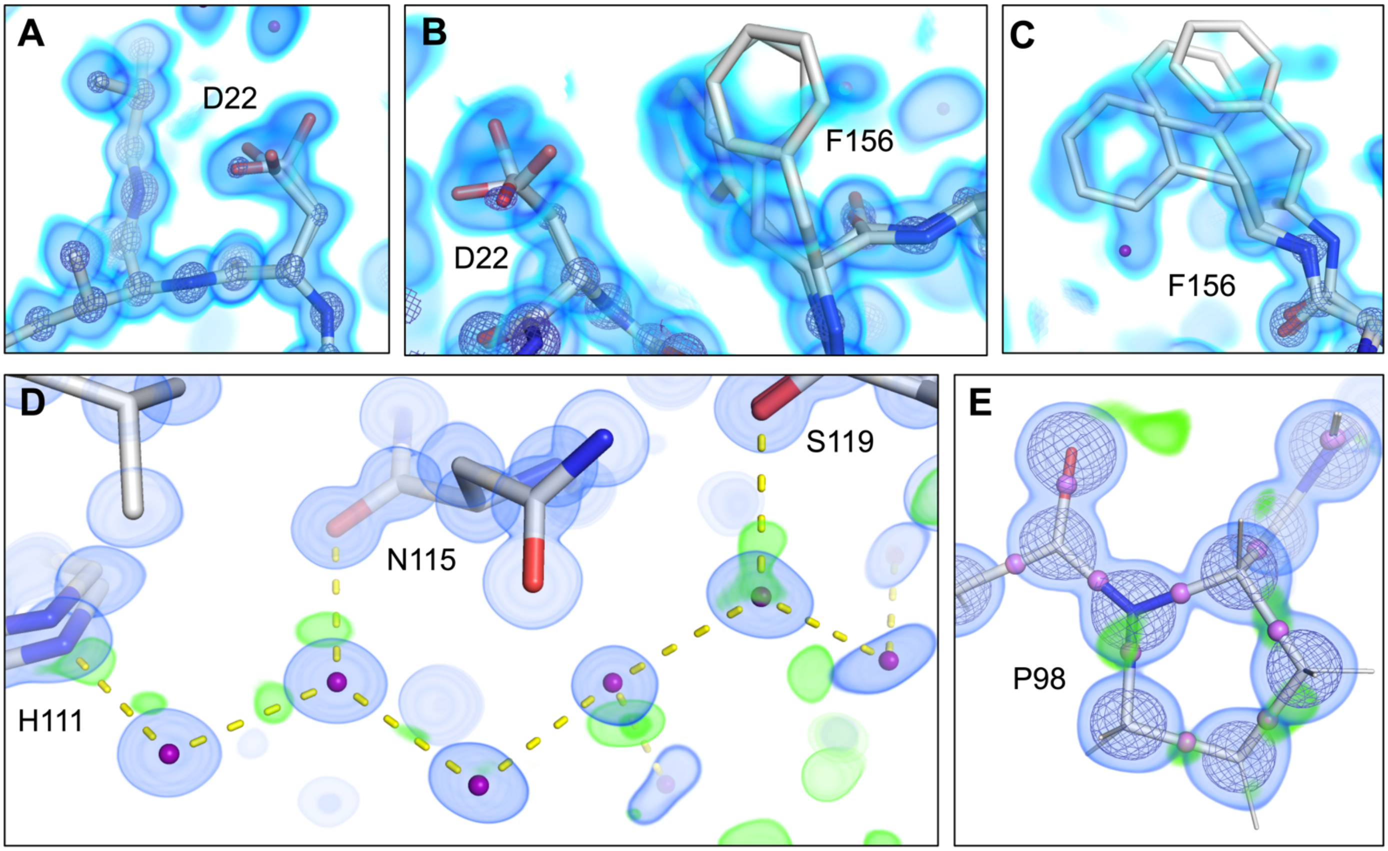


**Figure S1**. **Ultra-high resolution features in Mac1 electron density maps.**
**A,B,C)** Residues Asp22 and Phe156 both display conformational heterogeneity in high-resolution electron density maps of apo Mac1. The ultra-high resolution nature of the electron density is evident in (A), where atoms belonging to residues Ile23 and Leu24, as well as the backbone, appear as separate spheres at high contour. In (**A,B,C**) three 2mFo-Dfc maps are shown, contoured at 4.0 σ (blue mesh), 1.0 σ (blue volume), and 0.5 σ (cyan volume). **D)** Mac1 electron density reveals ordered water networks, including difference density that suggests the positions of hydrogen atoms within the water network. In (**D**), a $2mF_O$-$Df_C$ map is shown, contoured at 1.5 σ (blue volume), and a mFo-DFc map is also shown, contoured at 2.5 σ (green volume). Hydrogen bonding interactions are depicted as dashed yellow lines. **E)** Two electron density maps are shown for P98, a $2mF_O$-$Df_C$ ma, contoured at 1.5 σ (blue volume), and a $mF_O$-$Df_C$ map, contoured at 2.0 σ (green volume). The green peaks of positive difference density between heavy atom positions suggest covalent bonding densities, and we refined a model of Mac1 that included interatomic scatterers (IAS), shown in violet in the figure.

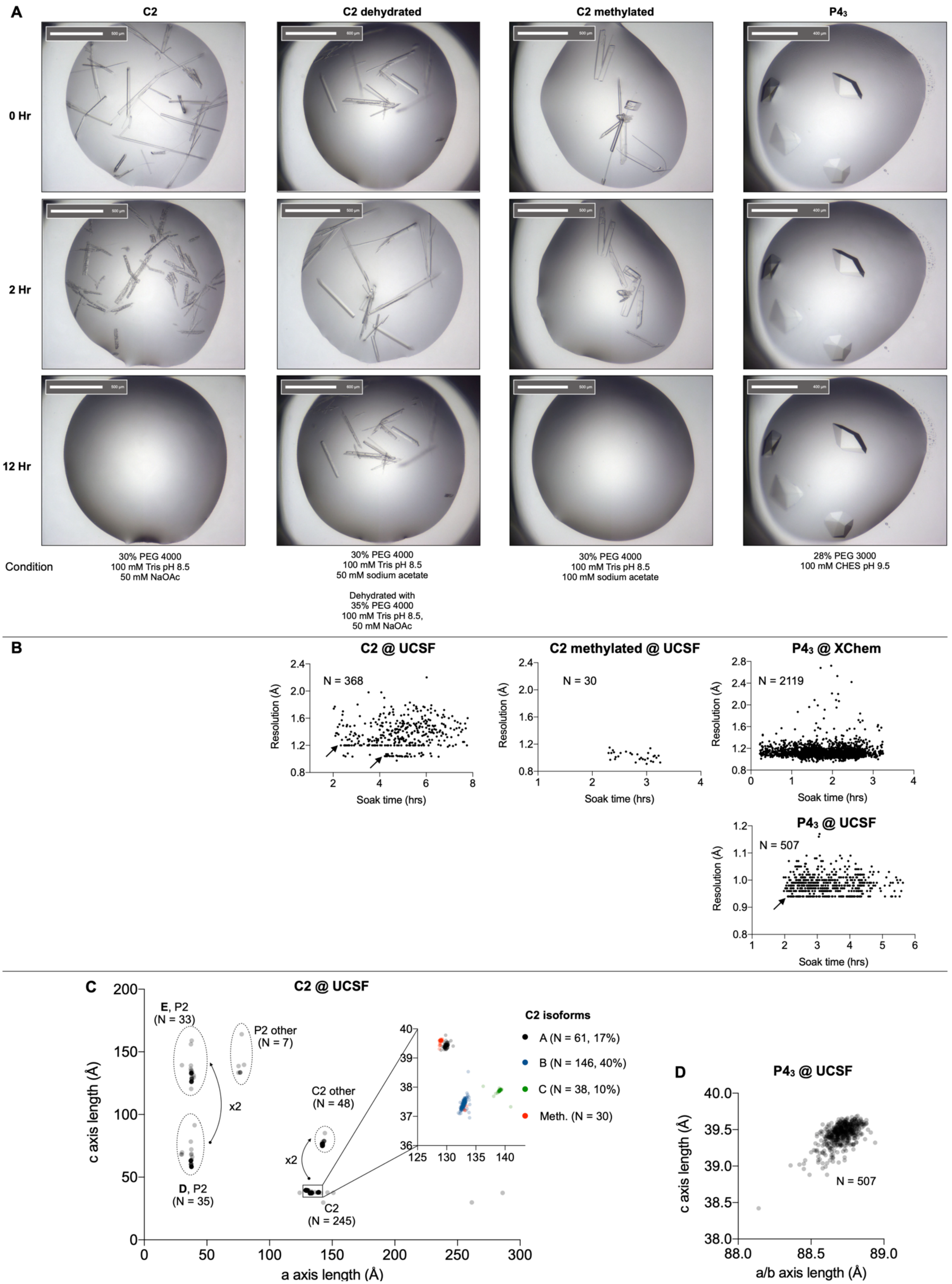
A
C2
C2 dehydrated
C2 methylated
P4₃
0 Hr
2 Hr
12 Hr
Condition
30% PEG 4000
100 mM Tris pH 8.5
50 mM NaOAc
30% PEG 4000
100 mM Tris pH 8.5
50 mM sodium acetate
Dehydrated with
35% PEG 4000
100 mM Tris pH 8.5,
50 mM NaOAc
30% PEG 4000
100 mM Tris pH 8.5
100 mM sodium acetate
28% PEG 3000
100 mM CHES pH 9.5
B
C2 @ UCSF
N = 368
C2 methylated @ UCSF
N = 30
P4₃ @ XChem
N = 2119
P4₃ @ UCSF
N = 507
Resolution (Å)
Soak time (hrs)
C
C2 @ UCSF
E, P2
(N = 33)
P2 other
(N = 7)
x2
C2 other
(N = 48)
D, P2
(N = 35)
C2
(N = 245)
C2 isoforms
A (N = 61, 17%)
B (N = 146, 40%)
C (N = 38, 10%)
Meth. (N = 30)
c axis length (Å)
a axis length (Å)
D
P4₃ @ UCSF
N = 507
c axis length (Å)
a/b axis length (Å)

**Figure S2. Comparison of isomorphism and DMSO tolerance of the C2 and $P4_3$ crystals. A)** Images of crystals after soaking with 10% DMSO for 0, 2 and 12 hours. **B)** Resolution of the three crystal forms as a function of soak time for the datasets collected at XChem and UCSF. The arrows indicate where the measurement of high resolution reflections was limited by the experimental setup. **C)** Multiple isoforms were observed for the C2 crystals after dehydration. Isoforms were distinguished based on differences in the a and c unit cell lengths. Arrows indicate where doubling of the a or c axis occurred. Inset: the majority of the datasets that were indexed in C2 (245, 84%) could be clustered into three isoforms (A, B and C). Of the 30 datasets collected for crystals grown from methylated protein, the majority (28) were similar to the A isoform. **D)** The $P4_3$ crystals were isomorphous.

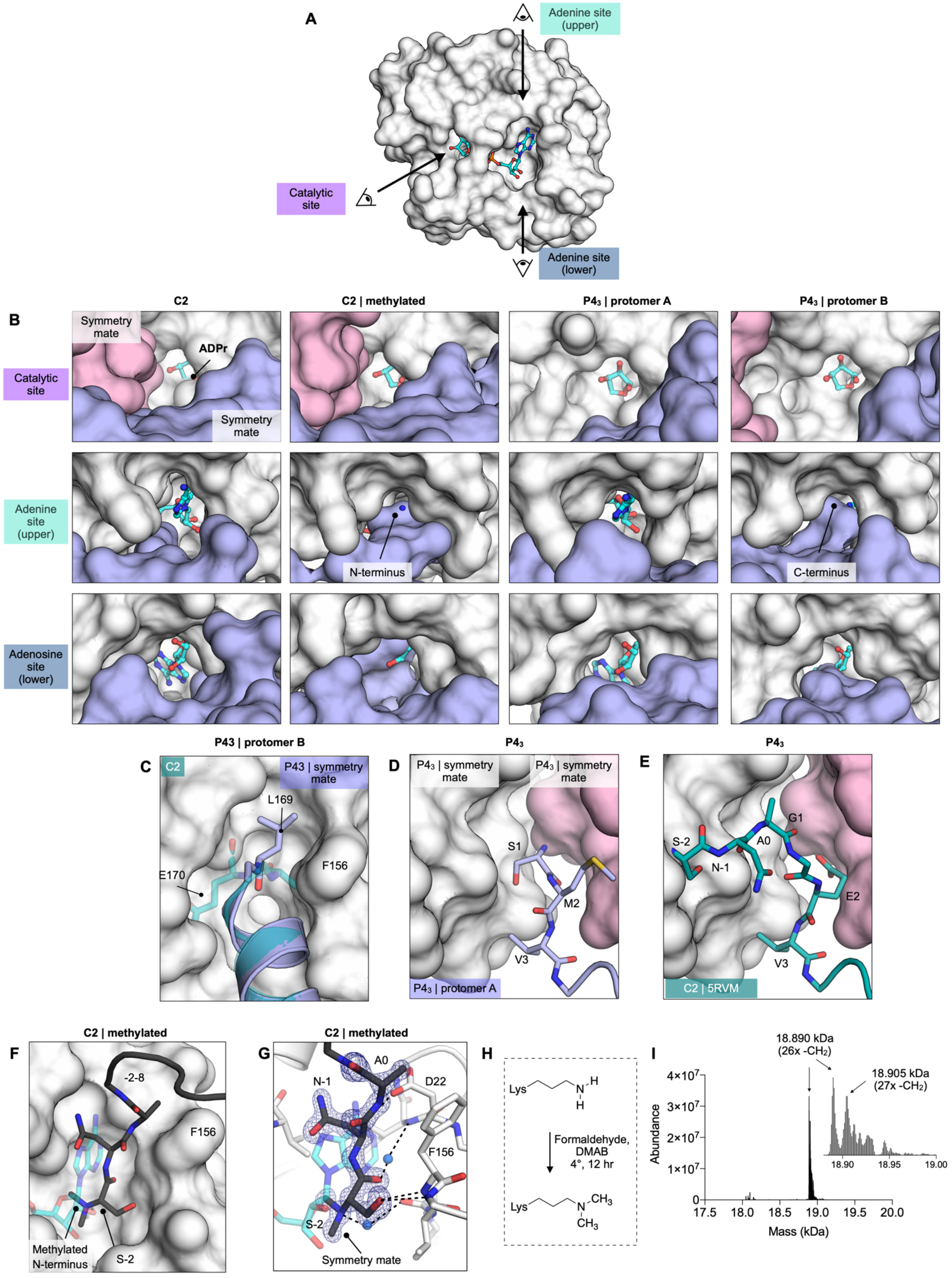

A
Adenine site (upper)
Catalytic site
Adenine site (lower)
B
C2
C2 | methylated
P4$_3$ | protomer A
P4$_3$ | protomer B
Symmetry mate
ADPr
Catalytic site
Symmetry mate
Adenine site (upper)
N-terminus
C-terminus
Adenosine site (lower)
P43 | protomer B
C
C2
P43 | symmetry mate
L169
E170
F156
P4$_3$
D
P4$_3$ | symmetry mate
P4$_3$ | symmetry mate
S1
M2
V3
P4$_3$ | protomer A
P4$_3$
E
S-2
N-1
A0
G1
E2
V3
C2 | 5RVM
C2 | methylated
F
-2-8
F156
Methylated N-terminus
S-2
C2 | methylated
G
A0
N-1
D22
F156
S-2
Symmetry mate
H
Lys
Formaldehyde, DMAB 4°, 12 hr
I
18.890 kDa (26x -CH$_2$)
18.905 kDa (27x -CH$_2$)
Abundance
Mass (kDa)

**Figure S3. Crystal packing in Mac1 crystals determines active site accessibility. A)** Active site access in the C2 and $P4_3$ crystals. Mac1 is shown as a white surface with ADP-ribose bound in the active site shown as cyan sticks. The three access points are indicated with arrows. **B)** Crystal packing defines the three access points. The catalytic site is partially obstructed in the C2 crystals, but open in both protomers of the $P4_3$ crystals. In both the methylated C2 crystals and protomer B of the $P4_3$ crystals, the adenosine site is obstructed. **C)** The C-terminal leucine (Leu169) of the $P4_3$ construct occupies the adenosine site of a symmetry mate. The adenosine site is shown as a white surface and the C-terminal residues with blue sticks/cartoon. The C2 sequence (transparent teal cartoon/sticks) has an additional residue at the C-terminus (Glu170), and is therefore incompatible with the $P4_3$ crystal packing. **C)** The N-terminal residues of the $P4_3$ sequence (blue sticks) pack between two symmetry mates (white and pink surface). Compared to the $P4_3$ sequence, the C2 sequence contains a substitution (Met2Glu) and a three residue insertion (Asn-Ala-Gly). These residues were typically disordered, however, they were resolved in one of the fragment structures (ZINC157088 | 5RVM) (shown aligned to the $P4_3$ protomer A in **(E)**). Like differences in the C-termini, differences in the N-termini may have contributed to the distinct crystal packing seen for the two Mac1 structures reported in this work. **F)** The adenosine site was obstructed by a symmetry mate in the structure determined from crystals grown using methylated C2 protein. **G)** In the structure of methylated Mac1, the side-chain hydroxyl of Ser-2 occupies the oxyanion subsite. Electron density ($2mF_O$-$DF_C$) is shown as a blue mesh, contoured at 1.5 σ. **H)** Free amines were methylated using formaldehyde and dimethylamine borane (DMAB). The reaction is shown for lysine, however, based on the electron density shown in **(G)**, the N-terminal amine was methylated as well. The methylated amines would be protonated at the pH used to grow crystals (pH 8.5). **I)** LC/MS analysis of methylated Mac1 (C2 construct). The mass spectrum was deconvoluted using MaxEnt1. The major peak (18.89 kDa) is consistent with the methylation of 13 lysine residues (26x -CH2). The minor peak (18.905 kDa + 15 Da) suggests that methylation was not 100% complete.

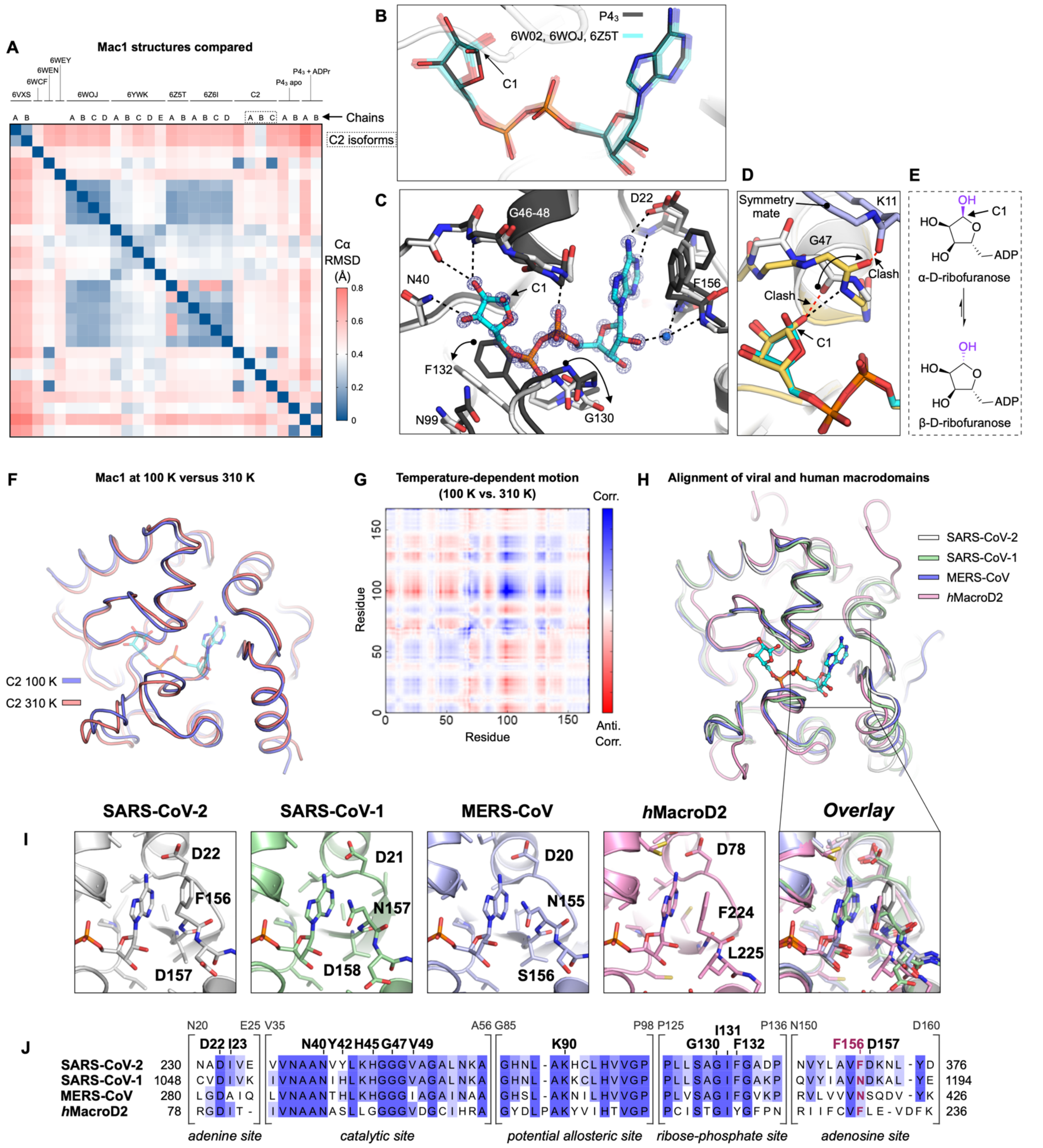


**Figure S4. Structure and sequence comparison of Mac1 with related viral and human macrodomains. A)** Heatmap showing the Cα RMSD values after Cα alignment for 10 previously reported SARS-CoV-2 Mac1 structures (6VXS, 6W02, 6W6Y, 6WCF, 6WEN, 6WOJ, 6WEY, 6YWK, 6Z5T, 6Z6I) and the new structures reported in this work. **B)** Stick representation showing the previously reported Mac1-ADPr structures (cyan sticks) and the new structure determined using $P4_3$ crystals (grey sticks). The agreement between ADP-ribose is excellent, despite different anomers of the terminal ribose being present (*α* in the $P4_3$ structure, *β* in the previously reported structures). **C)** The structural changes previously reported to occur upon ADPr binding are captured by the Mac1-ADPr structure determined using $P4_3$ crystals. The apo $P4_3$ structure is shown with dark gray sticks, with arrows indicating the changes in protein conformation upon ADP-ribose binding (white sticks). Electron density ($2mF_O$-$DF_C$) is contoured at 4 σ (blue mesh). **D)** The *α*-anomer of the terminal ribose of ADP-ribose was observed in the $P4_3$ crystal form (cyan and white sticks). In previously reported structures (e.g. PDB 6W02, yellow sticks), a flip in Gly47 allows the β-anomer to bind by removing a steric block (red dashed line) and forming a new H-bond (black dashed line). However, the Gly47 flip is incompatible with the $P4_3$ crystal form because it would clash with the Lys11 carbonyl of a symmetry mate (blue sticks). In *α*-anomer, the anomeric hydroxyl is orientated away from Gly47, and binding can proceed without the peptide flip. **E)** Interconversion between ribose anomers in solution. **F)** Comparison of SARS-CoV-2 Nsp3 Mac1 structures at 100 K (blue) and 310 K (red). The adenosine diphosphoribose ligand shown in the figure (cyan) is modeled according to its position in PDB 6W02. **G)** Correlation plot showing structural differences between the 100 K and 310 K structures. To generate the plot, the 100 K and 310 K structures were aligned and difference vectors were calculated between identical Cα atoms in the two structures. The plot shows all pairwise dot products between these difference vectors, revealing the extent to which temperature-dependent changes are correlated across the structure. Positive dot products (positive correlations) are colored blue, while dot products (negative correlations) are shown in red. The pattern of positive and negative correlations is characteristic of a hinge-bending motion. **H)** Alignment of three coronavirus macrodomain structures with a human macrodomain (hMacroD2). ADPr from the SARS-CoV-2 structure is shown with cyan sticks. **I)** Comparison of the adenosine binding site highlighting key residues involved in adenine and fragment interaction. The adenine coordination by Phe156 is unique to SARS-CoV-2 amongst betacoronaviruses and replaced in SARS-CoV-1 (PDB 2FAV) and MERS-CoV (PDB 5HOL) with asparagine. Human macrodomains including MacroD2 (PDB 4IQY) interact with adenine as SARS-CoV-2 Nsp3 Mac1 with a phenylalanine in this position which needs to be considered for achieving inhibitor selectivity for viral over human macrodomains. **J)** Sequence alignment showing conservation of residues in the ADP-ribose, catalytic and potential allosteric sites which are targeted by the fragments. Residue numbers on top refer to the construct residue numbering of SARS-CoV-2 Nsp3 Mac1. Numbers on either end of the alignment are residue numbers in the full-length proteins. The adenine-coordinating Phe156 is highlighted in red.

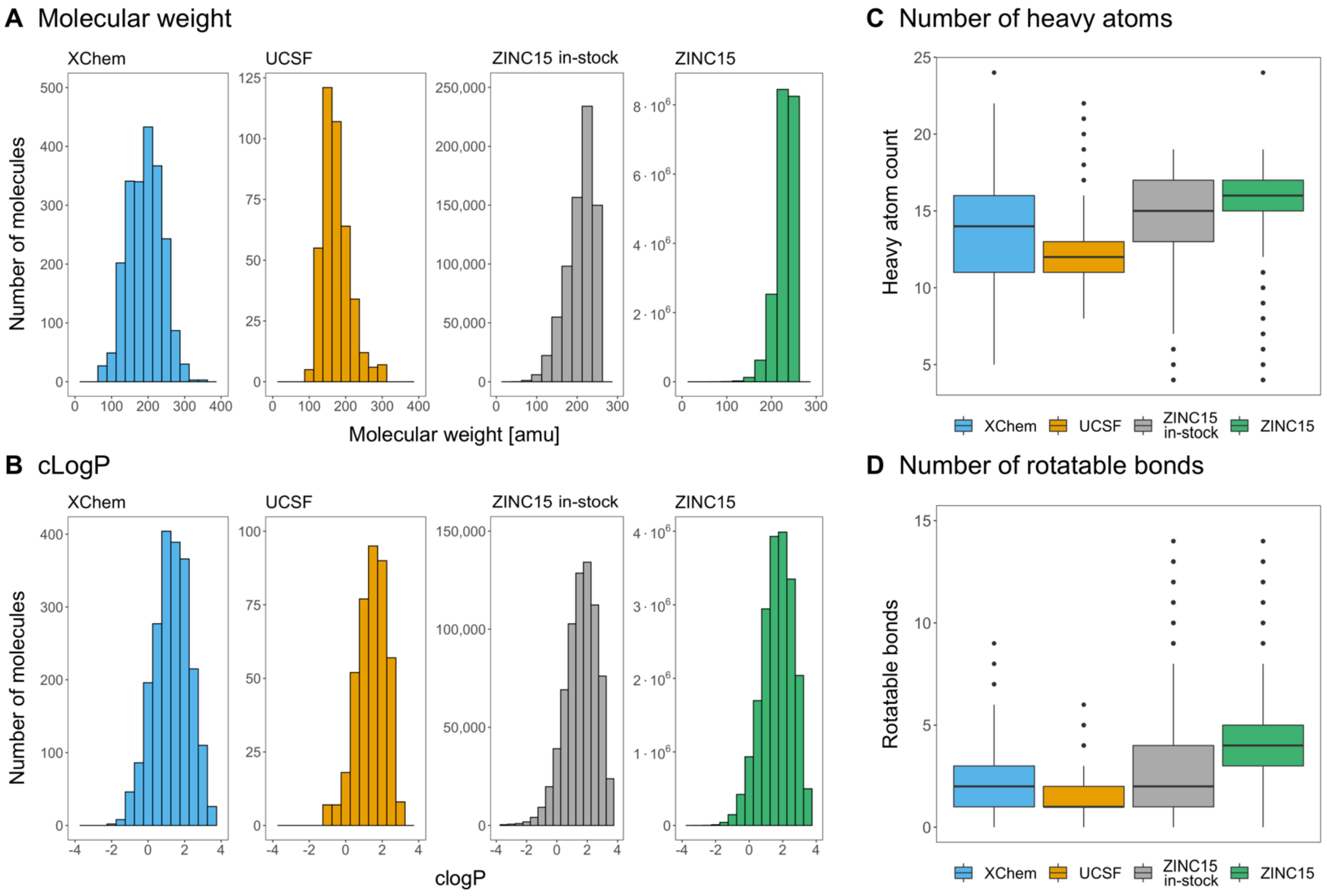


**E** Scaffold and chemotype analysis

| Fragment Library | Number of molecules | Distinct Bemis-Murcko scaffolds | Number of pyrimidines | Number of anions |
|---|---|---|---|---|
| XChem | 2,126 | 809 | 72 (3.39%) | 115 (5.4%) |
| UCSF | 411 | 179 | 12 (2.92%) | 144 (35%) |
| ZINC15 (in-stock) | 722,963 | 69,244 | 41,531 (5.74%) | 85,398 (11.8%) |
| ZINC15 | 20,006,175 | 803,333 | 890,199 (4.44%) | 739,184 (3.7%) |

**Figure S5. Physical properties, scaffold and chemotype analysis of screened fragment libraries.**

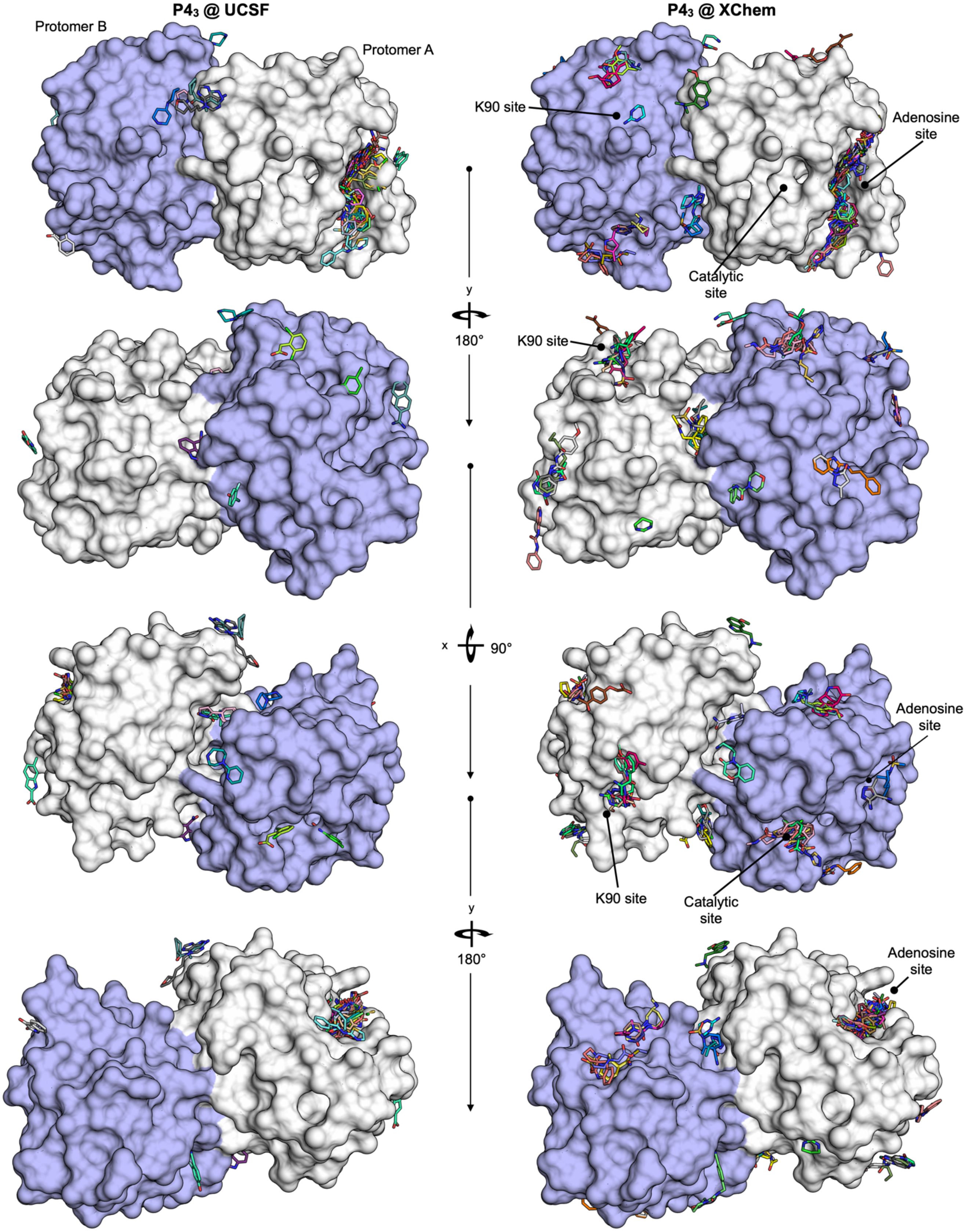


**Figure S6. Overview of fragment binding to protomer A (white surface) and protomer B (blue surface) of the P4$_3$ crystals.**

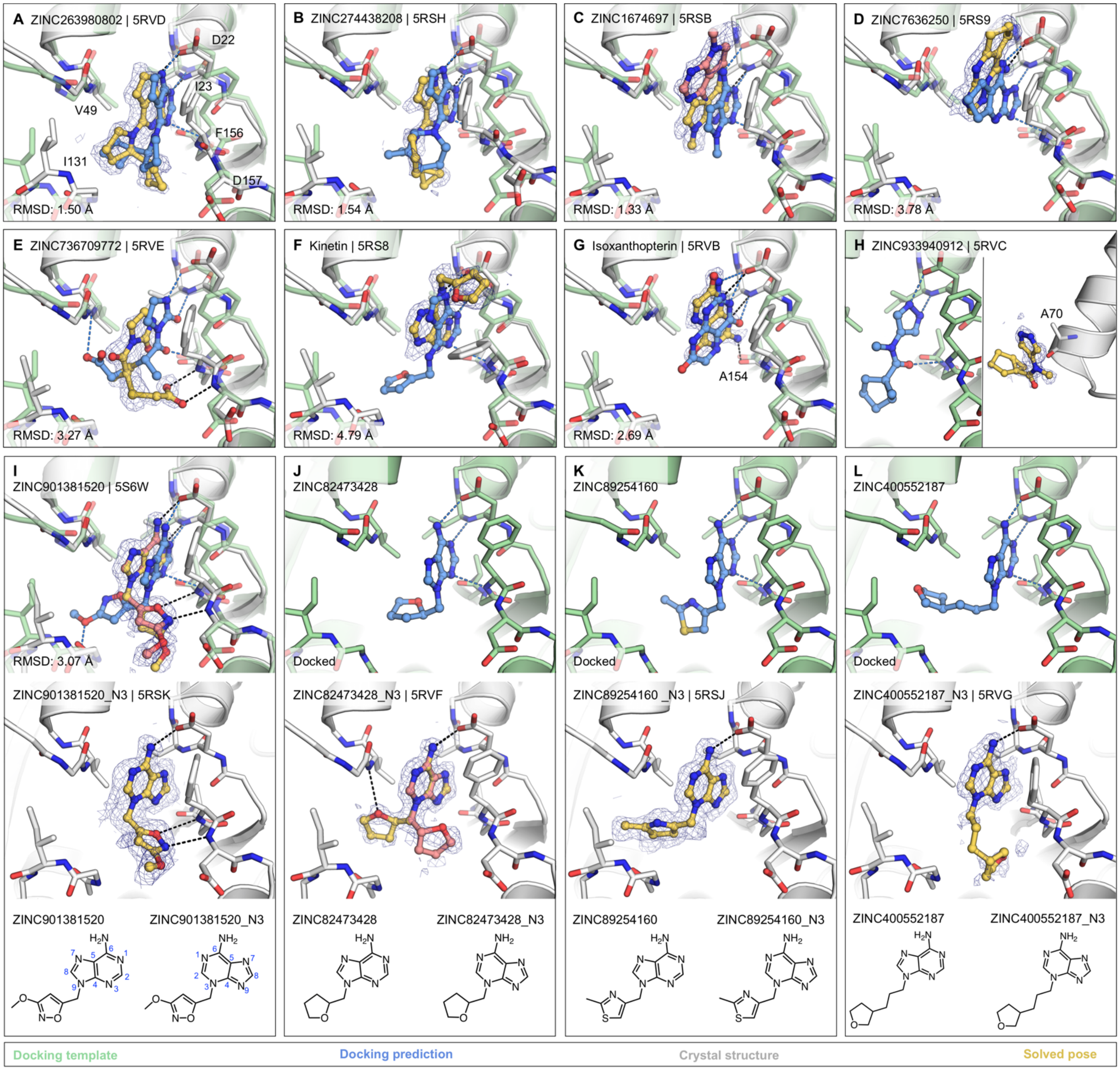


**Figure S7. Additional soaking hits from docking and adenine-N3 vs -N9-alkylated isomers.**

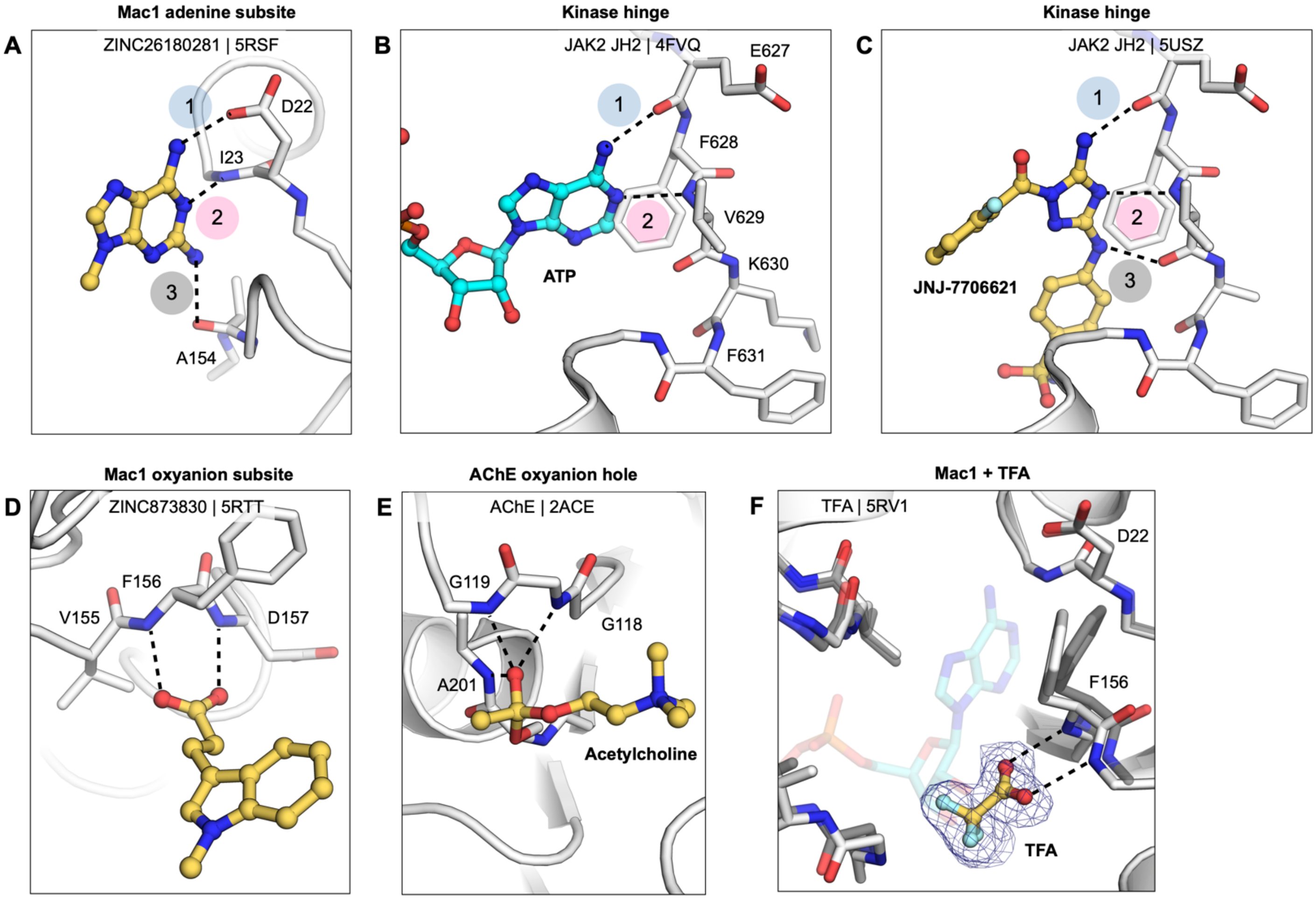


**Figure S8. Mac1 subsites compared to the adenine binding site in kinases and the oxyanion binding site in carboxylesterases. A)** Key features of the Mac1 adenine subsite are illustrated by the structure of ZINC26180281 (PDB 5RSF). Hydrogen bonds are formed between the C6-amine of the adenine scaffold and the backbone nitrogen of Ile23, and between N1 of the adenine scaffold and the side-chain carboxylate of Asp22. The C2 amine of ZINC26180281 forms a non-canonical H-bond to the backbone carbonyl oxygen of Ala154. **B)** Adenine recognition is similar in the pseudokinase domain of JAK2, however, the C6-amine forms an H-bond to a backbone carbonyl oxygen rather than a side-chain carboxylate. Adenine binding occurs at the hinge residues that connect the N- and C-terminal lobes of the catalytic domain. Interactions that mimic adenine binding to the hinge residues are conserved in the majority of kinase inhibitors (*57*). Like ZINC2618028, kinase inhibitors exploit non-canonical H-bonds. The 1,2,4-triazole derived inhibitor shown in **(C)** forms an H-bond to the backbone carbonyl oxygen of Lys630. **D)** The fragment screens against Mac1 identified 47 oxyanions binding to the backbone nitrogens of Phe156 and Asp157. A comparable oxyanion recognition motif is present in acetylcholinesterase (AChE) **(E).** In AChE, this motif stabilizes negative charge on the oxyanion transition state. **F)** Trifluoroacetic acid (TFA), present as a counter ion for ZINC3860798, was clearly defined in PanDDA event maps binding to the oxyanion subsite. TFA was also observed binding to the oxyanion subsite for fragments ZINC35185198 and ZINC51658946. The docking fragment ZINC263392672 also contained TFA, but no TFA was observed in the oxyanion subsite (PDB 5RSG).

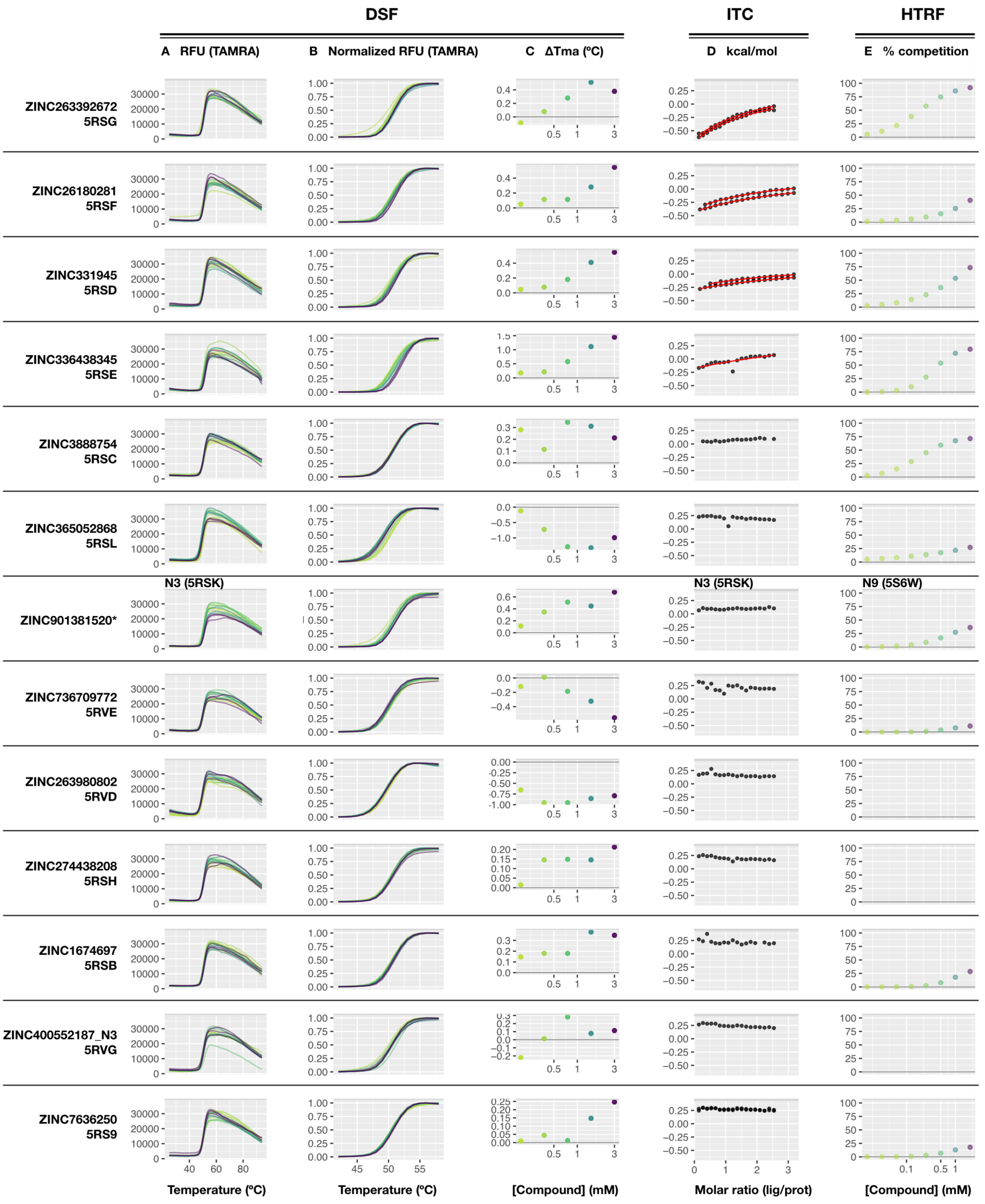

DSF
ITC
HTRF
A RFU (TAMRA)
B Normalized RFU (TAMRA)
C ΔTma (ºC)
D kcal/mol
E % competition
ZINC263392672
5RSG
ZINC26180281
5RSF
ZINC331945
5RSD
ZINC336438345
5RSE
ZINC3888754
5RSC
ZINC365052868
5RSL
N3 (5RSK)
N3 (5RSK)
N9 (5S6W)
ZINC901381520*
ZINC736709772
5RVE
ZINC263980802
5RVD
ZINC274438208
5RSH
ZINC1674697
5RSB
ZINC400552187_N3
5RVG
ZINC7636250
5RS9
Temperature (ºC)
Temperature (ºC)
[Compound] (mM)
Molar ratio (lig/prot)
[Compound] (mM)

**Figure S9. Comparison of DSF, HTRF, and ITC results for compounds tested in all assays. A)** Raw, un-normalized DSF data for the full measured temperature range (25 - 94ºC) demonstrates the absence of confounding changes in curve shape for all compounds. **B)** Normalized raw DSF data, enlarged to visualize compound-induced thermal shifts. **C)** Changes in $Tm_a$ observed in the presence of fragments (0-3 mM fragment). **D)** Integrated heat peaks as a function of binding site saturation shown as black dots. The red line represents a non-linear least squares (NLLS) fit using a single-site binding model. **E)** Dose-response curves showing competition of the fragments with an ADPr-conjugated peptide for Mac1 binding. (*) ZINC901381520_N3 was tested in DSF and ITC, ZINC901381520_N9 was tested in HTRF.
