## Supplementary Data Set 2 for "Fragment Binding to the Nsp3 Macrodomain of SARS-CoV-2 Identified Through Crystallographic Screening and Computational Docking"

**Structures of fragments bound to Mac1.**

Only fragments binding in the adenosine site, catalytic site or K90 site are shown. Fragments are shown as yellow sticks. Mac1 is shown in the fragment-bound state (white sticks) and the apo state (transparent teal sticks). Hydrogen bonds are shown as dashed black lines. The Mac1 subsites are defined in Figure 2. The dataset code, PDB code and compound ID are given for each fragment.

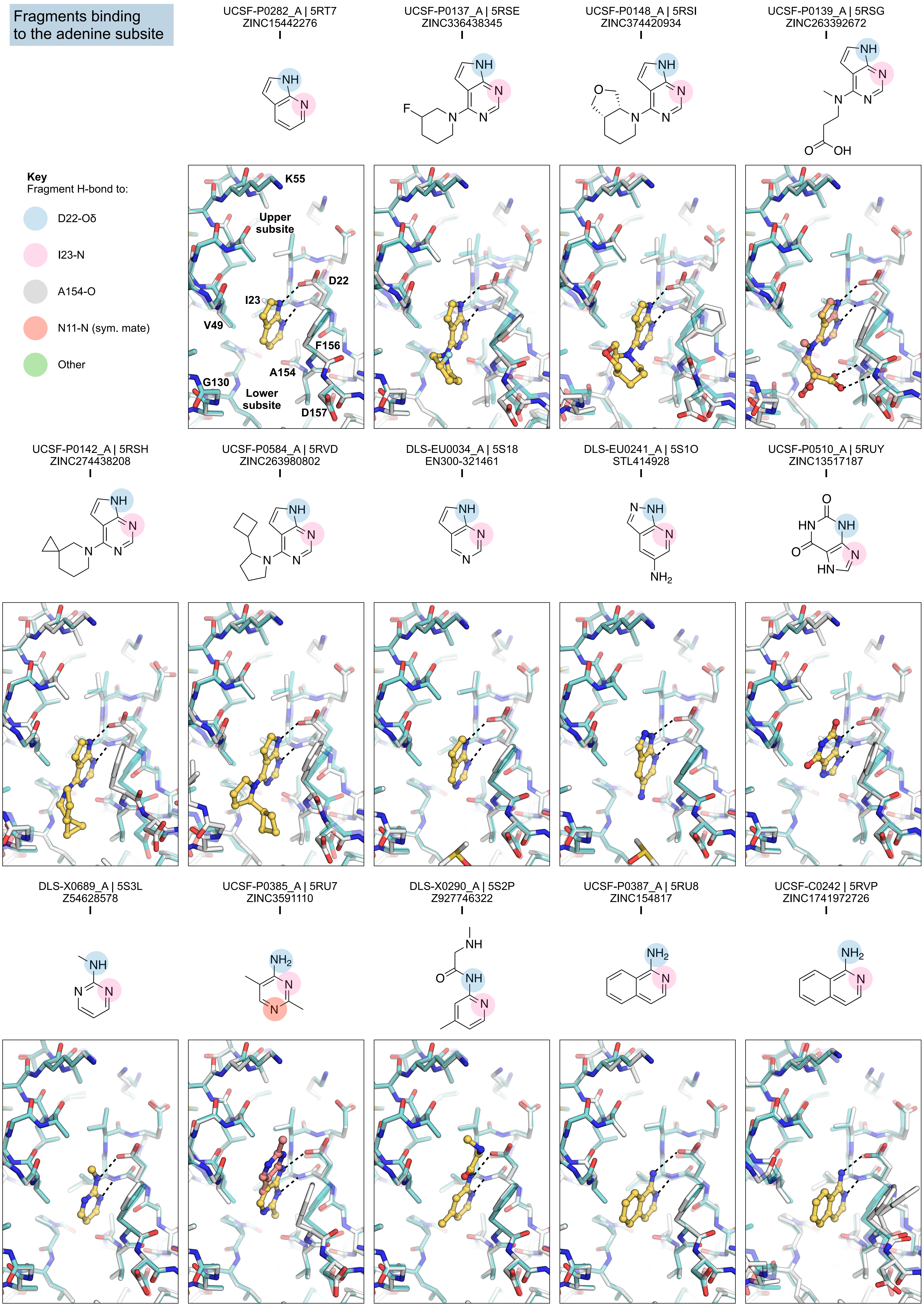
Fragments binding to the adenine subsite
UCSF-P0282_A | 5RT7
ZINC15442276
UCSF-P0137_A | 5RSE
ZINC336438345
UCSF-P0148_A | 5RSI
ZINC374420934
UCSF-P0139_A | 5RSG
ZINC263392672
Key
Fragment H-bond to:
D22-Oδ
I23-N
A154-O
N11-N (sym. mate)
Other
K55
Upper subsite
D22
I23
V49
F156
A154
G130
Lower subsite
D157
UCSF-P0142_A | 5RSH
ZINC274438208
UCSF-P0584_A | 5RVD
ZINC263980802
DLS-EU0034_A | 5S18
EN300-321461
DLS-EU0241_A | 5S1O
STL414928
UCSF-P0510_A | 5RUY
ZINC13517187
DLS-X0689_A | 5S3L
Z54628578
UCSF-P0385_A | 5RU7
ZINC3591110
DLS-X0290_A | 5S2P
Z927746322
UCSF-P0387_A | 5RU8
ZINC154817
UCSF-C0242 | 5RVP
ZINC1741972726

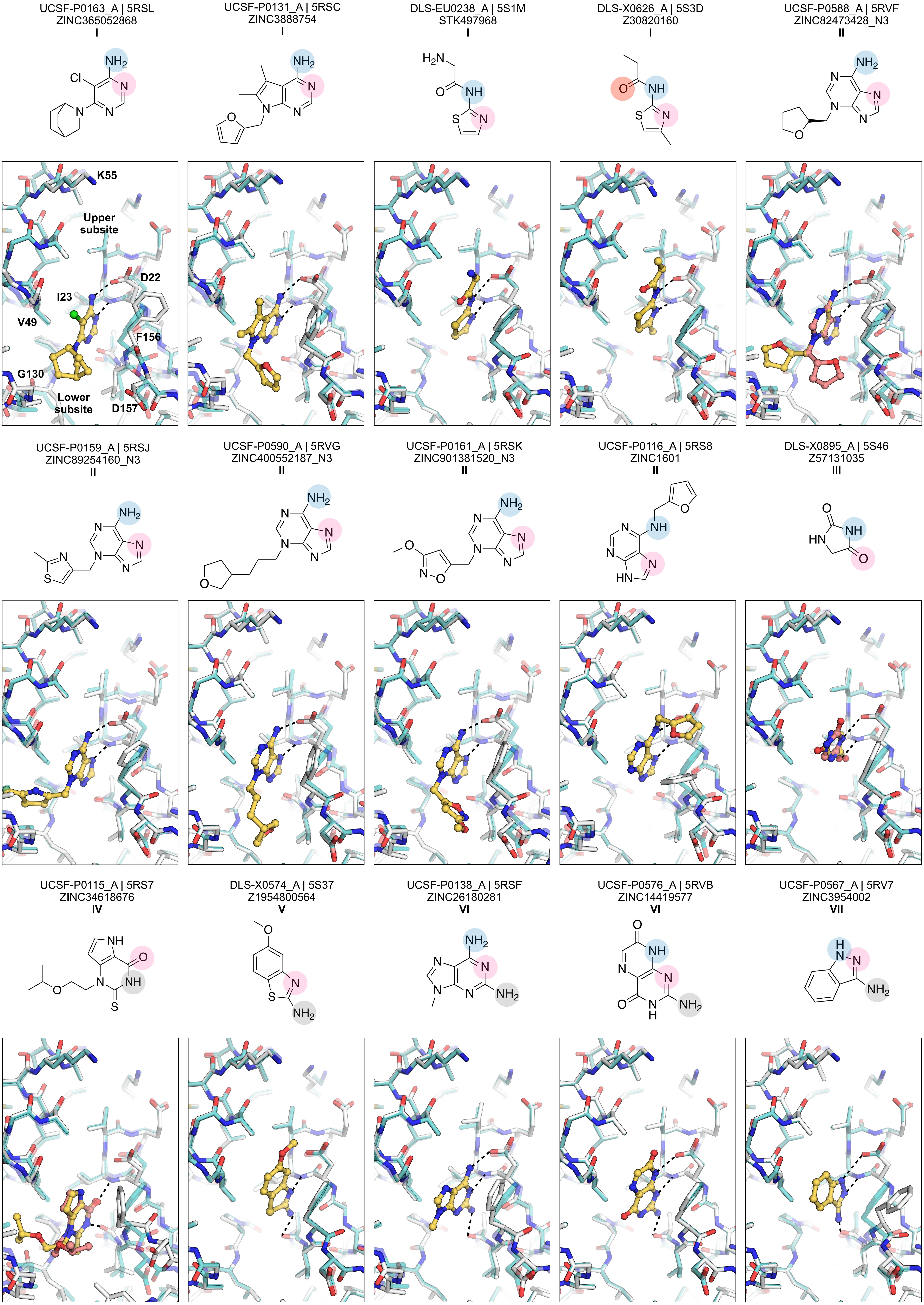
UCSF-P0163_A | 5RSL
ZINC365052868
I
NH2
Cl
N
N
N
K55
Upper subsite
D22
I23
V49
F156
G130
Lower subsite
D157
UCSF-P0131_A | 5RSC
ZINC3888754
I
NH2
N
N
N
O
DLS-EU0238_A | 5S1M
STK497968
I
H2N
O
NH
S
N
DLS-X0626_A | 5S3D
Z30820160
I
O
NH
S
N
UCSF-P0588_A | 5RVF
ZINC82473428_N3
II
NH2
N
N
N
N
O
UCSF-P0159_A | 5RSJ
ZINC89254160_N3
II
NH2
N
N
N
N
N
S
UCSF-P0590_A | 5RVG
ZINC400552187_N3
II
NH2
N
N
N
N
O
UCSF-P0161_A | 5RSK
ZINC901381520_N3
II
NH2
N
N
N
N
O
N-O
UCSF-P0116_A | 5RS8
ZINC1601
II
O
NH
N
N
N
HN
DLS-X0895_A | 5S46
Z57131035
III
O
HN
NH
O
UCSF-P0115_A | 5RS7
ZINC34618676
IV
NH
O
O
N
NH
S
DLS-X0574_A | 5S37
Z1954800564
V
O
N
S
NH2
UCSF-P0138_A | 5RSF
ZINC26180281
VI
NH2
N
N
N
N
NH2
UCSF-P0576_A | 5RVB
ZINC14419577
VI
O
NH
N
N
O
N
H
NH2
UCSF-P0567_A | 5RV7
ZINC3954002
VII
H
N-N
NH2

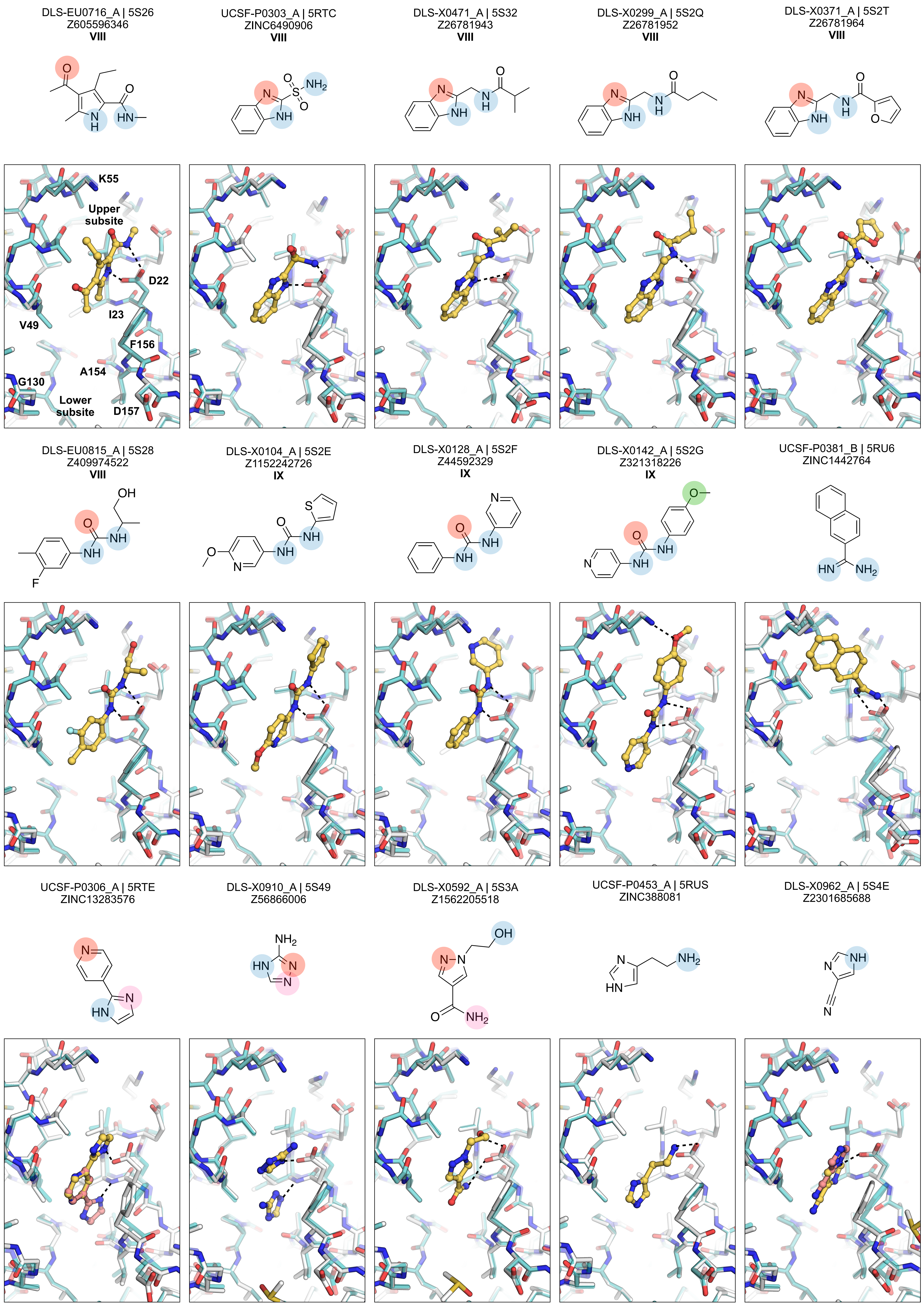
DLS-EU0716_A | 5S26
Z605596346
VIII
UCSF-P0303_A | 5RTC
ZINC6490906
VIII
DLS-X0471_A | 5S32
Z26781943
VIII
DLS-X0299_A | 5S2Q
Z26781952
VIII
DLS-X0371_A | 5S2T
Z26781964
VIII
K55
Upper subsite
D22
I23
V49
F156
A154
G130
Lower subsite
D157
DLS-EU0815_A | 5S28
Z409974522
VIII
DLS-X0104_A | 5S2E
Z1152242726
IX
DLS-X0128_A | 5S2F
Z44592329
IX
DLS-X0142_A | 5S2G
Z321318226
IX
UCSF-P0381_B | 5RU6
ZINC1442764
UCSF-P0306_A | 5RTE
ZINC13283576
DLS-X0910_A | 5S49
Z56866006
DLS-X0592_A | 5S3A
Z1562205518
UCSF-P0453_A | 5RUS
ZINC388081
DLS-X0962_A | 5S4E
Z2301685688

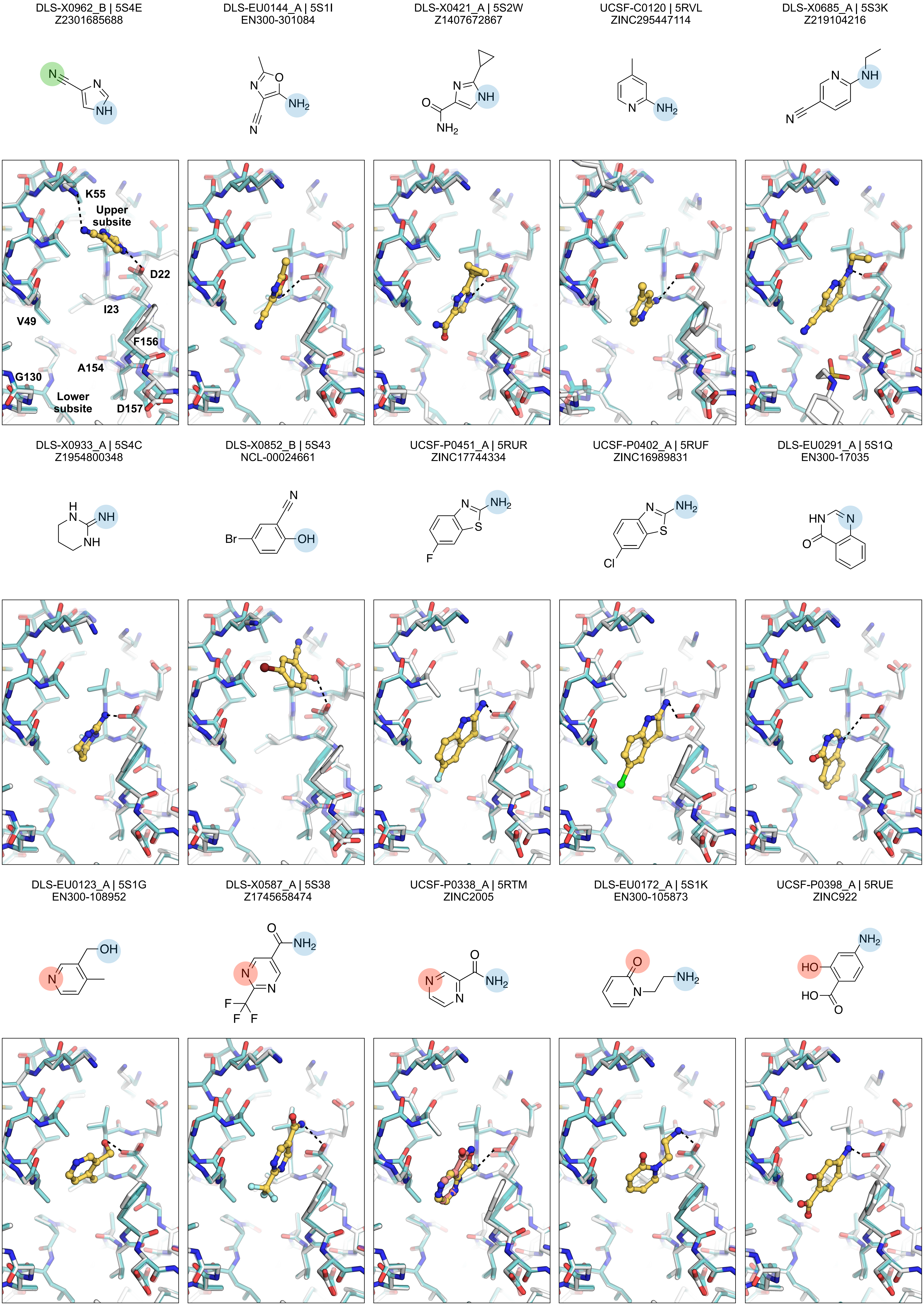
DLS-X0962_B | 5S4E
Z2301685688
DLS-EU0144_A | 5S1I
EN300-301084
DLS-X0421_A | 5S2W
Z1407672867
UCSF-C0120 | 5RVL
ZINC295447114
DLS-X0685_A | 5S3K
Z219104216
K55
Upper subsite
D22
I23
V49
F156
G130
A154
Lower subsite
D157
DLS-X0933_A | 5S4C
Z1954800348
DLS-X0852_B | 5S43
NCL-00024661
UCSF-P0451_A | 5RUR
ZINC17744334
UCSF-P0402_A | 5RUF
ZINC16989831
DLS-EU0291_A | 5S1Q
EN300-17035
DLS-EU0123_A | 5S1G
EN300-108952
DLS-X0587_A | 5S38
Z1745658474
UCSF-P0338_A | 5RTM
ZINC2005
DLS-EU0172_A | 5S1K
EN300-105873
UCSF-P0398_A | 5RUE
ZINC922

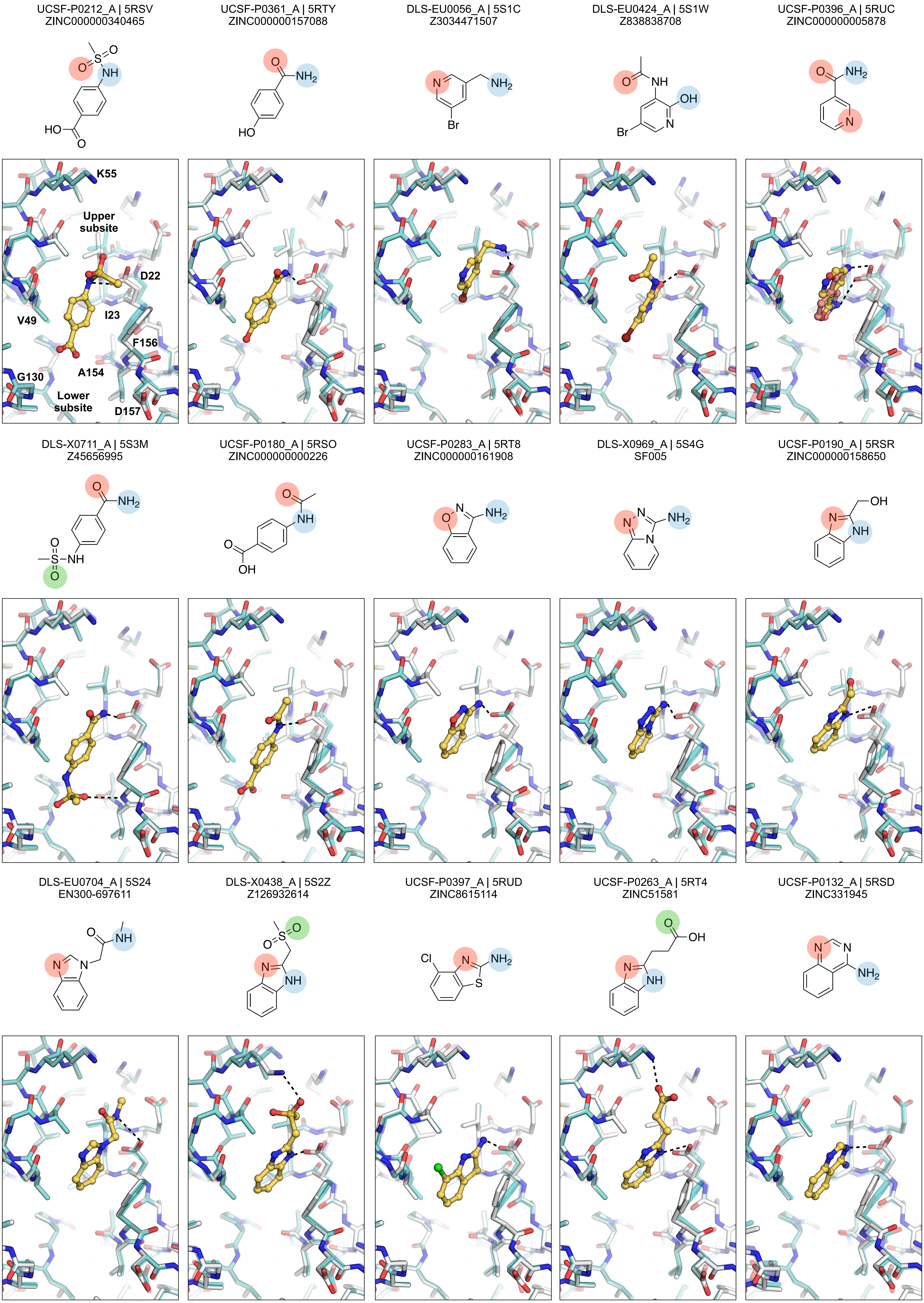

UCSF-P0212_A | 5RSV
ZINC000000340465
UCSF-P0361_A | 5RTY
ZINC000000157088
DLS-EU0056_A | 5S1C
Z3034471507
DLS-EU0424_A | 5S1W
Z838838708
UCSF-P0396_A | 5RUC
ZINC000000005878
K55
Upper subsite
D22
I23
V49
F156
A154
G130
Lower subsite
D157
DLS-X0711_A | 5S3M
Z45656995
UCSF-P0180_A | 5RSO
ZINC000000000226
UCSF-P0283_A | 5RT8
ZINC000000161908
DLS-X0969_A | 5S4G
SF005
UCSF-P0190_A | 5RSR
ZINC000000158650
DLS-EU0704_A | 5S24
EN300-697611
DLS-X0438_A | 5S2Z
Z126932614
UCSF-P0397_A | 5RUD
ZINC8615114
UCSF-P0263_A | 5RT4
ZINC51581
UCSF-P0132_A | 5RSD
ZINC331945

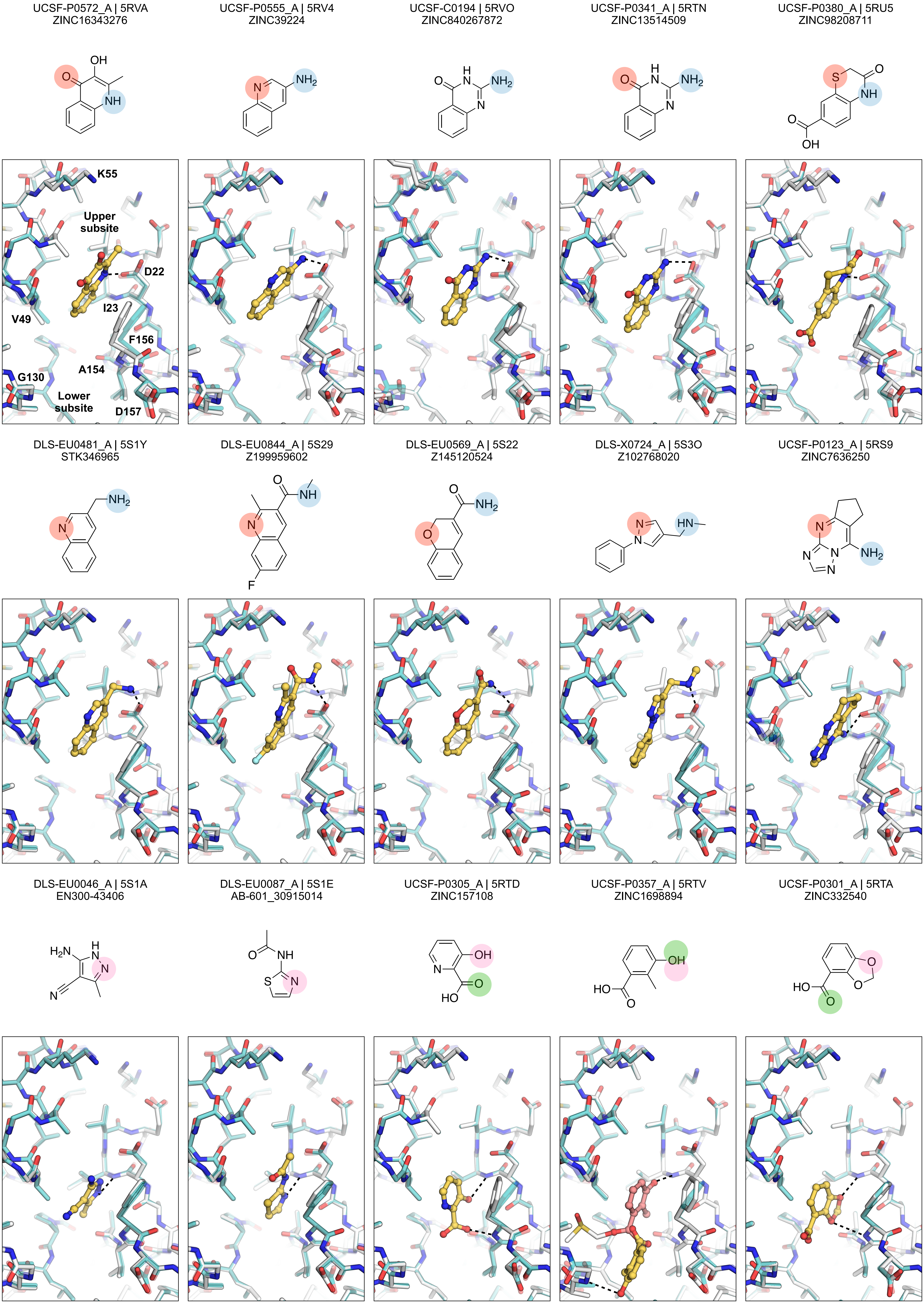
UCSF-P0572_A | 5RVA
ZINC16343276
UCSF-P0555_A | 5RV4
ZINC39224
UCSF-C0194 | 5RVO
ZINC840267872
UCSF-P0341_A | 5RTN
ZINC13514509
UCSF-P0380_A | 5RU5
ZINC98208711
K55
Upper subsite
D22
V49
I23
F156
A154
G130
Lower subsite
D157
DLS-EU0481_A | 5S1Y
STK346965
DLS-EU0844_A | 5S29
Z199959602
DLS-EU0569_A | 5S22
Z145120524
DLS-X0724_A | 5S3O
Z102768020
UCSF-P0123_A | 5RS9
ZINC7636250
DLS-EU0046_A | 5S1A
EN300-43406
DLS-EU0087_A | 5S1E
AB-601_30915014
UCSF-P0305_A | 5RTD
ZINC157108
UCSF-P0357_A | 5RTV
ZINC1698894
UCSF-P0301_A | 5RTA
ZINC332540

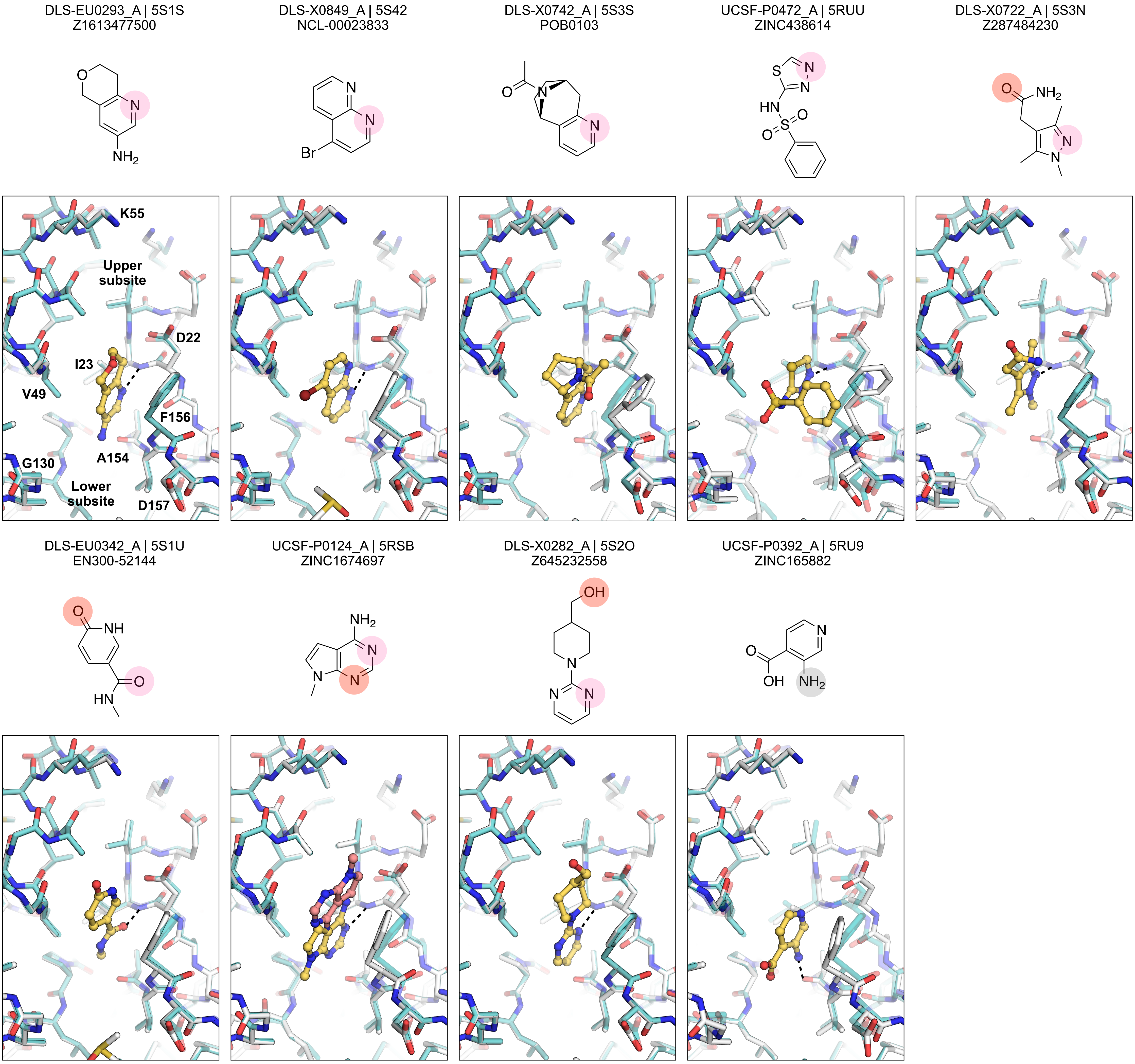

DLS-EU0293_A | 5S1S
Z1613477500
DLS-X0849_A | 5S42
NCL-00023833
DLS-X0742_A | 5S3S
POB0103
UCSF-P0472_A | 5RUU
ZINC438614
DLS-X0722_A | 5S3N
Z287484230
K55
Upper subsite
D22
I23
V49
F156
A154
G130
Lower subsite
D157
DLS-EU0342_A | 5S1U
EN300-52144
UCSF-P0124_A | 5RSB
ZINC1674697
DLS-X0282_A | 5S2O
Z645232558
UCSF-P0392_A | 5RU9
ZINC165882

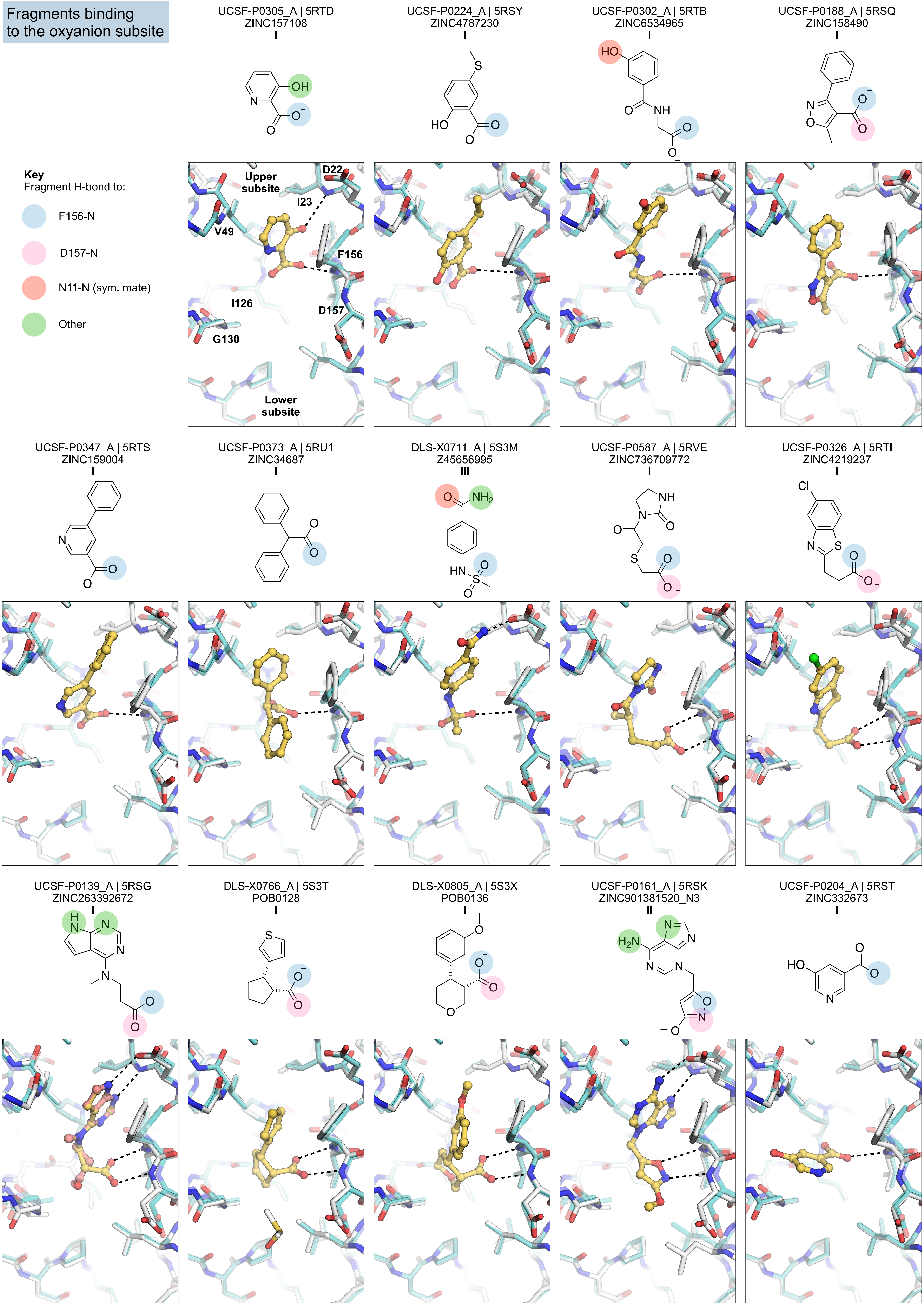

Fragments binding to the oxyanion subsite
Key
Fragment H-bond to:
F156-N
D157-N
N11-N (sym. mate)
Other
UCSF-P0305_A | 5RTD
ZINC157108
I
UCSF-P0224_A | 5RSY
ZINC4787230
I
UCSF-P0302_A | 5RTB
ZINC6534965
I
UCSF-P0188_A | 5RSQ
ZINC158490
I
Upper subsite
D22
I23
V49
F156
I126
D157
G130
Lower subsite
UCSF-P0347_A | 5RTS
ZINC159004
I
UCSF-P0373_A | 5RU1
ZINC34687
I
DLS-X0711_A | 5S3M
Z45656995
III
UCSF-P0587_A | 5RVE
ZINC736709772
I
UCSF-P0326_A | 5RTI
ZINC4219237
I
UCSF-P0139_A | 5RSG
ZINC263392672
I
DLS-X0766_A | 5S3T
POB0128
I
DLS-X0805_A | 5S3X
POB0136
I
UCSF-P0161_A | 5RSK
ZINC901381520_N3
II
UCSF-P0204_A | 5RST
ZINC332673
I

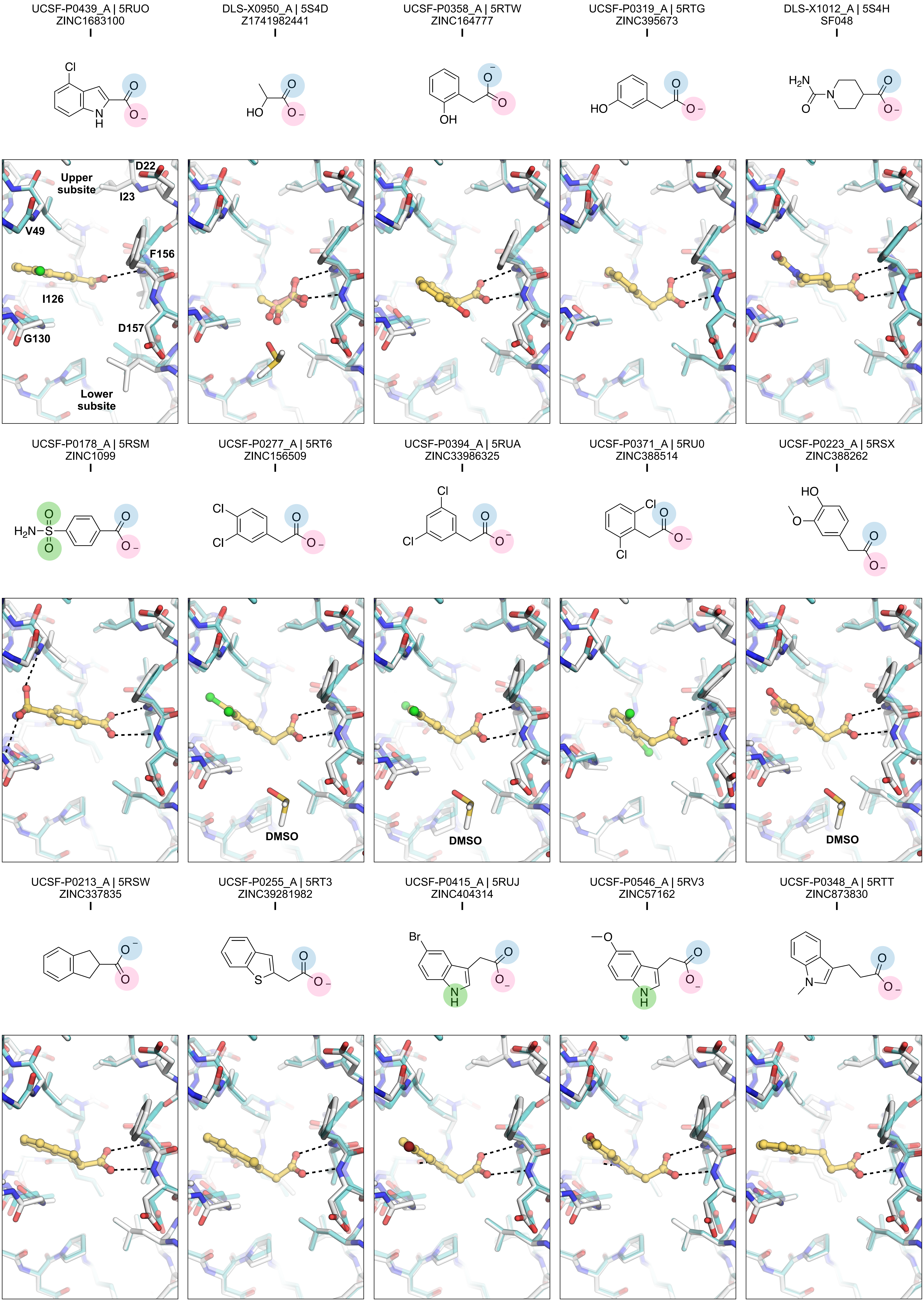
UCSF-P0439_A | 5RUO
ZINC1683100
DLS-X0950_A | 5S4D
Z1741982441
UCSF-P0358_A | 5RTW
ZINC164777
UCSF-P0319_A | 5RTG
ZINC395673
DLS-X1012_A | 5S4H
SF048
D22
Upper subsite
I23
V49
F156
I126
D157
G130
Lower subsite
UCSF-P0178_A | 5RSM
ZINC1099
UCSF-P0277_A | 5RT6
ZINC156509
UCSF-P0394_A | 5RUA
ZINC33986325
UCSF-P0371_A | 5RU0
ZINC388514
UCSF-P0223_A | 5RSX
ZINC388262
DMSO
DMSO
DMSO
UCSF-P0213_A | 5RSW
ZINC337835
UCSF-P0255_A | 5RT3
ZINC39281982
UCSF-P0415_A | 5RUJ
ZINC404314
UCSF-P0546_A | 5RV3
ZINC57162
UCSF-P0348_A | 5RTT
ZINC873830

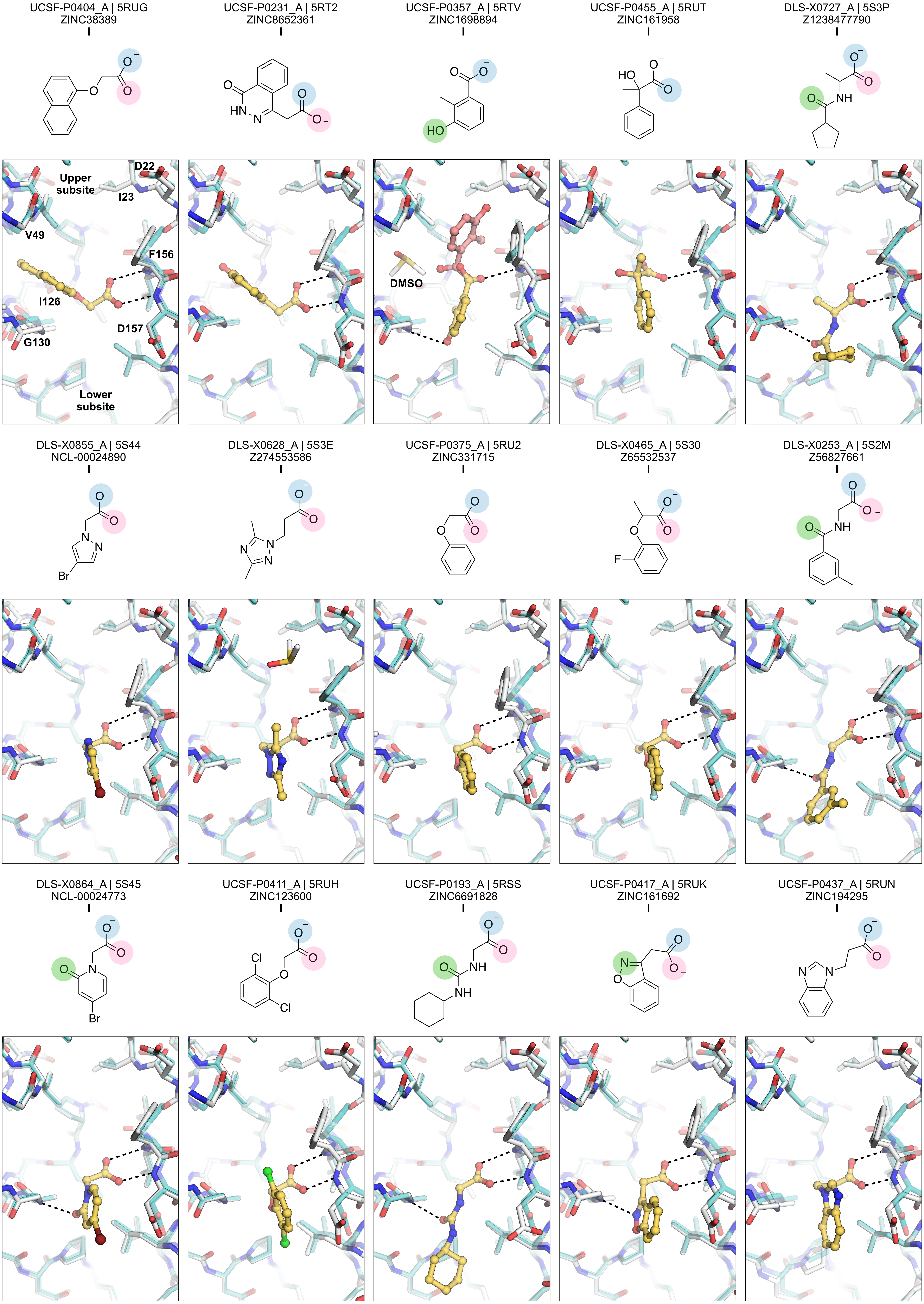
UCSF-P0404_A | 5RUG
ZINC38389
UCSF-P0231_A | 5RT2
ZINC8652361
UCSF-P0357_A | 5RTV
ZINC1698894
UCSF-P0455_A | 5RUT
ZINC161958
DLS-X0727_A | 5S3P
Z1238477790
Upper subsite
D22
I23
V49
F156
I126
D157
G130
Lower subsite
DMSO
DLS-X0855_A | 5S44
NCL-00024890
DLS-X0628_A | 5S3E
Z274553586
UCSF-P0375_A | 5RU2
ZINC331715
DLS-X0465_A | 5S30
Z65532537
DLS-X0253_A | 5S2M
Z56827661
DLS-X0864_A | 5S45
NCL-00024773
UCSF-P0411_A | 5RUH
ZINC123600
UCSF-P0193_A | 5RSS
ZINC6691828
UCSF-P0417_A | 5RUK
ZINC161692
UCSF-P0437_A | 5RUN
ZINC194295

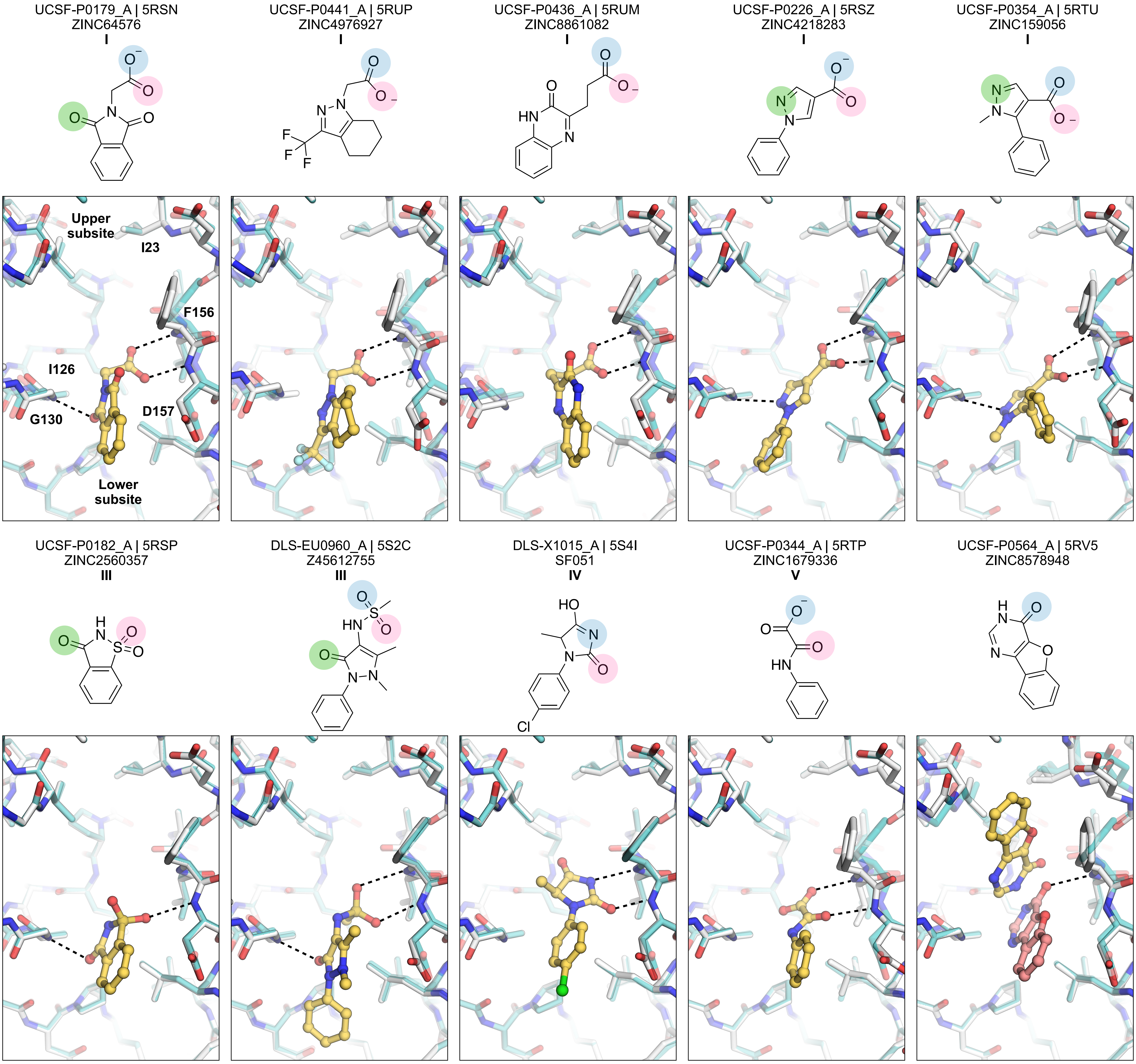

UCSF-P0179_A | 5RSN
ZINC64576
I
UCSF-P0441_A | 5RUP
ZINC4976927
I
UCSF-P0436_A | 5RUM
ZINC8861082
I
UCSF-P0226_A | 5RSZ
ZINC4218283
I
UCSF-P0354_A | 5RTU
ZINC159056
I
Upper subsite
I23
F156
I126
G130
D157
Lower subsite
UCSF-P0182_A | 5RSP
ZINC2560357
III
DLS-EU0960_A | 5S2C
Z45612755
III
DLS-X1015_A | 5S4I
SF051
IV
UCSF-P0344_A | 5RTP
ZINC1679336
V
UCSF-P0564_A | 5RV5
ZINC8578948

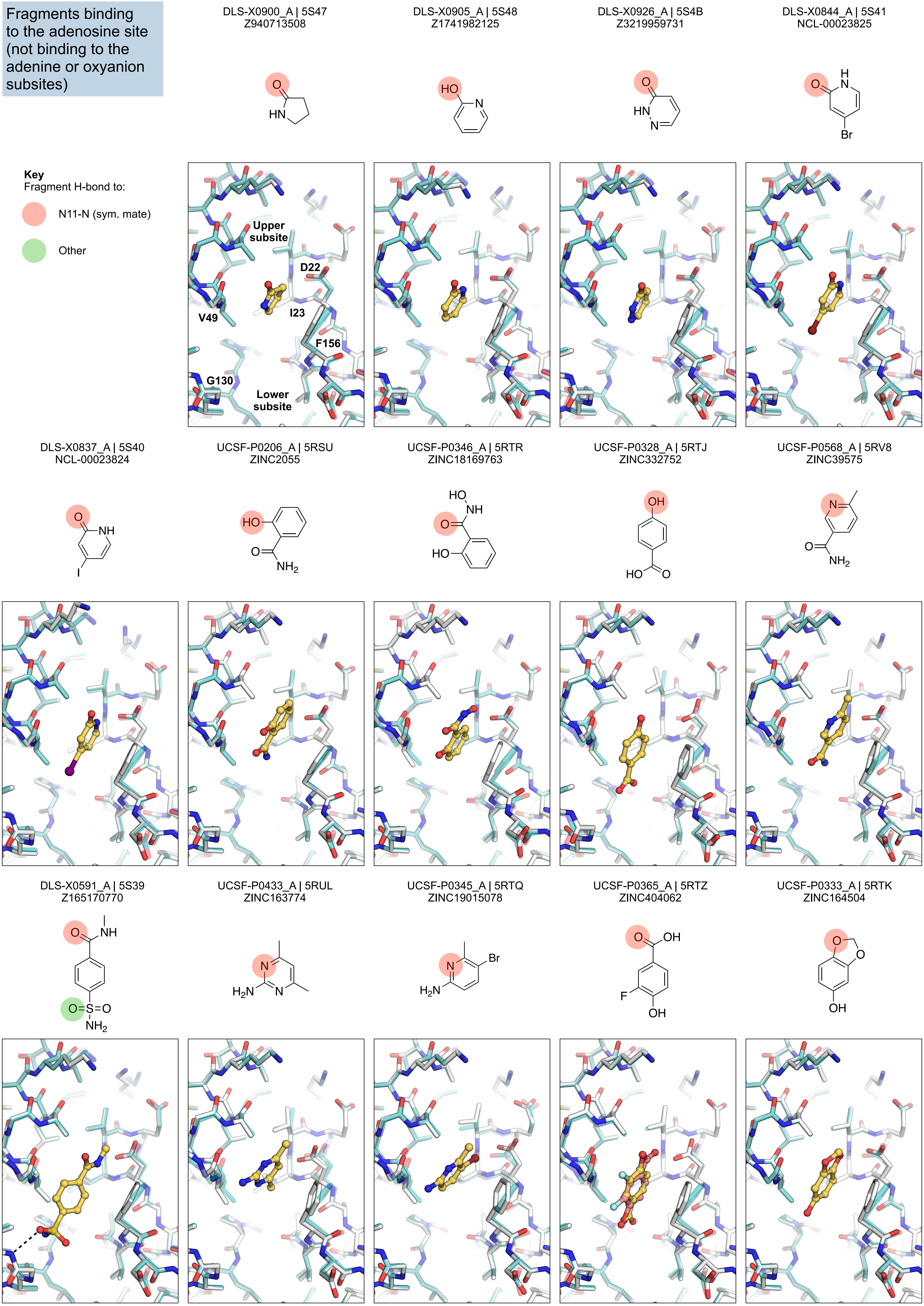
Fragments binding to the adenosine site (not binding to the adenine or oxyanion subsites)
Key
Fragment H-bond to:
N11-N (sym. mate)
Other
DLS-X0900_A | 5S47
Z940713508
DLS-X0905_A | 5S48
Z1741982125
DLS-X0926_A | 5S4B
Z3219959731
DLS-X0844_A | 5S41
NCL-00023825
Upper subsite
D22
V49
I23
F156
G130
Lower subsite
DLS-X0837_A | 5S40
NCL-00023824
UCSF-P0206_A | 5RSU
ZINC2055
UCSF-P0346_A | 5RTR
ZINC18169763
UCSF-P0328_A | 5RTJ
ZINC332752
UCSF-P0568_A | 5RV8
ZINC39575
DLS-X0591_A | 5S39
Z165170770
UCSF-P0433_A | 5RUL
ZINC163774
UCSF-P0345_A | 5RTQ
ZINC19015078
UCSF-P0365_A | 5RTZ
ZINC404062
UCSF-P0333_A | 5RTK
ZINC164504

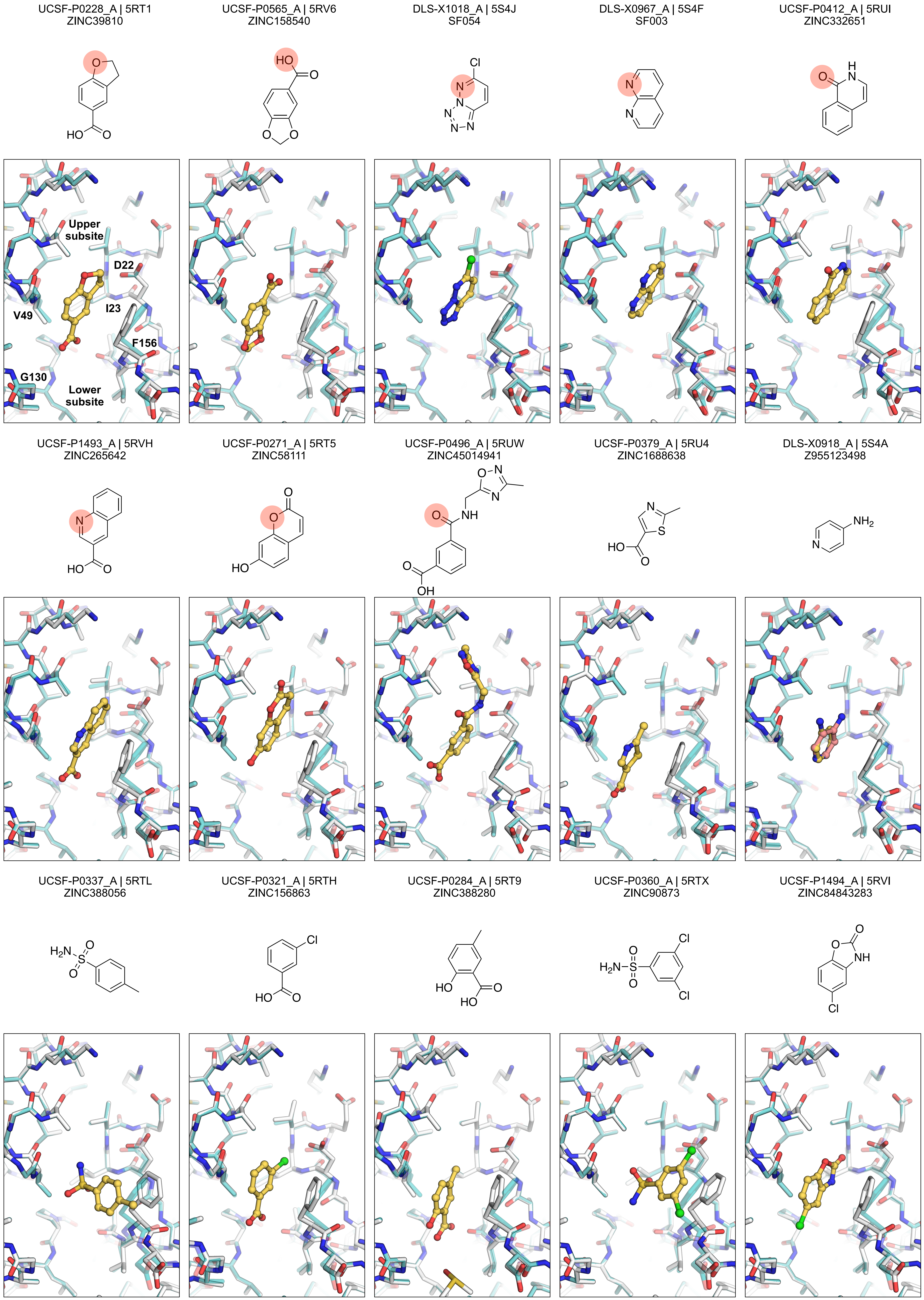
UCSF-P0228_A | 5RT1
ZINC39810
UCSF-P0565_A | 5RV6
ZINC158540
DLS-X1018_A | 5S4J
SF054
DLS-X0967_A | 5S4F
SF003
UCSF-P0412_A | 5RUI
ZINC332651
Upper subsite
D22
V49
I23
F156
G130
Lower subsite
UCSF-P1493_A | 5RVH
ZINC265642
UCSF-P0271_A | 5RT5
ZINC58111
UCSF-P0496_A | 5RUW
ZINC45014941
UCSF-P0379_A | 5RU4
ZINC1688638
DLS-X0918_A | 5S4A
Z955123498
UCSF-P0337_A | 5RTL
ZINC388056
UCSF-P0321_A | 5RTH
ZINC156863
UCSF-P0284_A | 5RT9
ZINC388280
UCSF-P0360_A | 5RTX
ZINC90873
UCSF-P1494_A | 5RVI
ZINC84843283

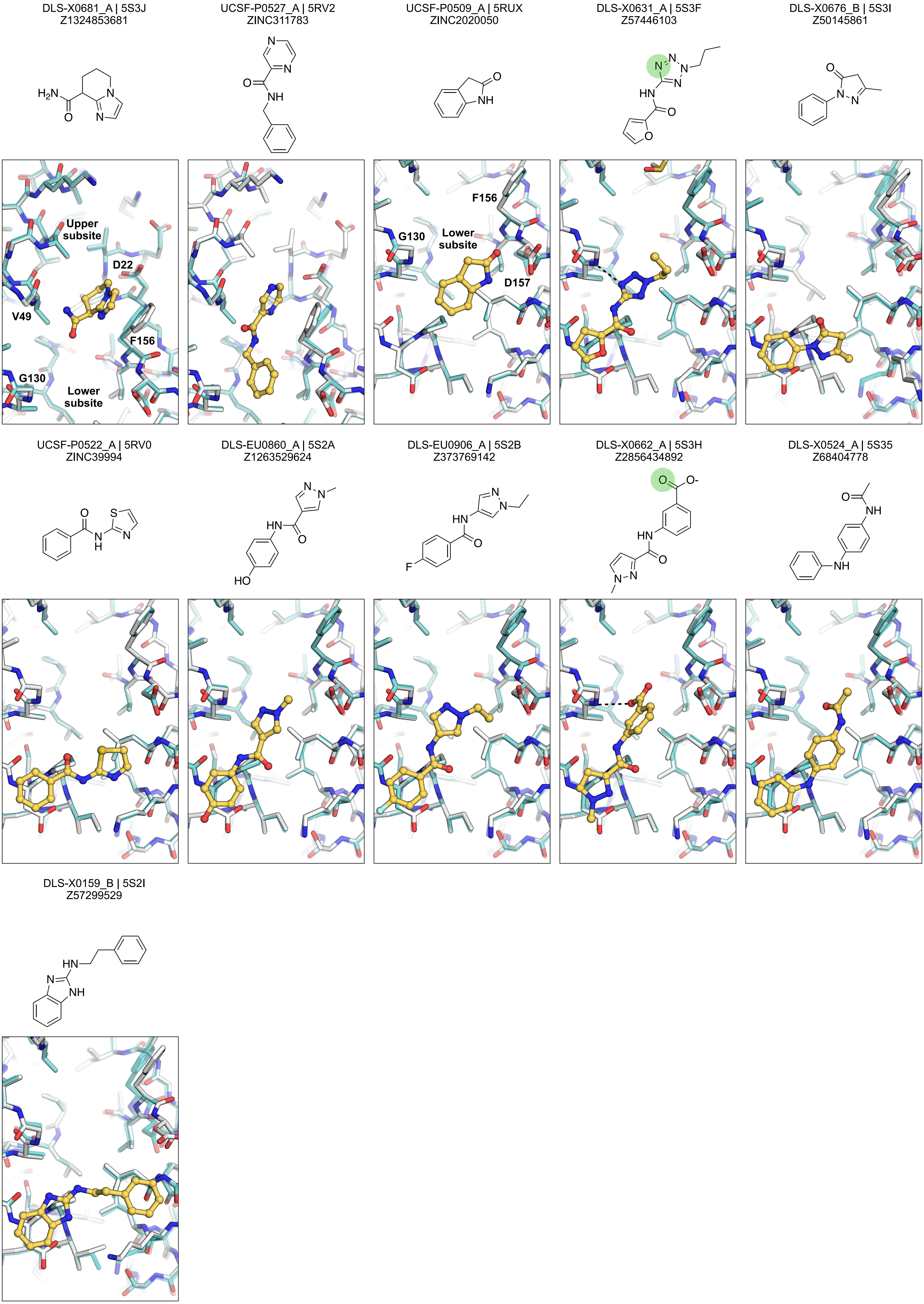
DLS-X0681_A | 5S3J
Z1324853681
Upper subsite
D22
V49
F156
G130
Lower subsite
UCSF-P0527_A | 5RV2
ZINC311783
UCSF-P0509_A | 5RUX
ZINC2020050
F156
G130
Lower subsite
D157
DLS-X0631_A | 5S3F
Z57446103
DLS-X0676_B | 5S3I
Z50145861
UCSF-P0522_A | 5RV0
ZINC39994
DLS-EU0860_A | 5S2A
Z1263529624
DLS-EU0906_A | 5S2B
Z373769142
DLS-X0662_A | 5S3H
Z2856434892
DLS-X0524_A | 5S35
Z68404778
DLS-X0159_B | 5S2I
Z57299529

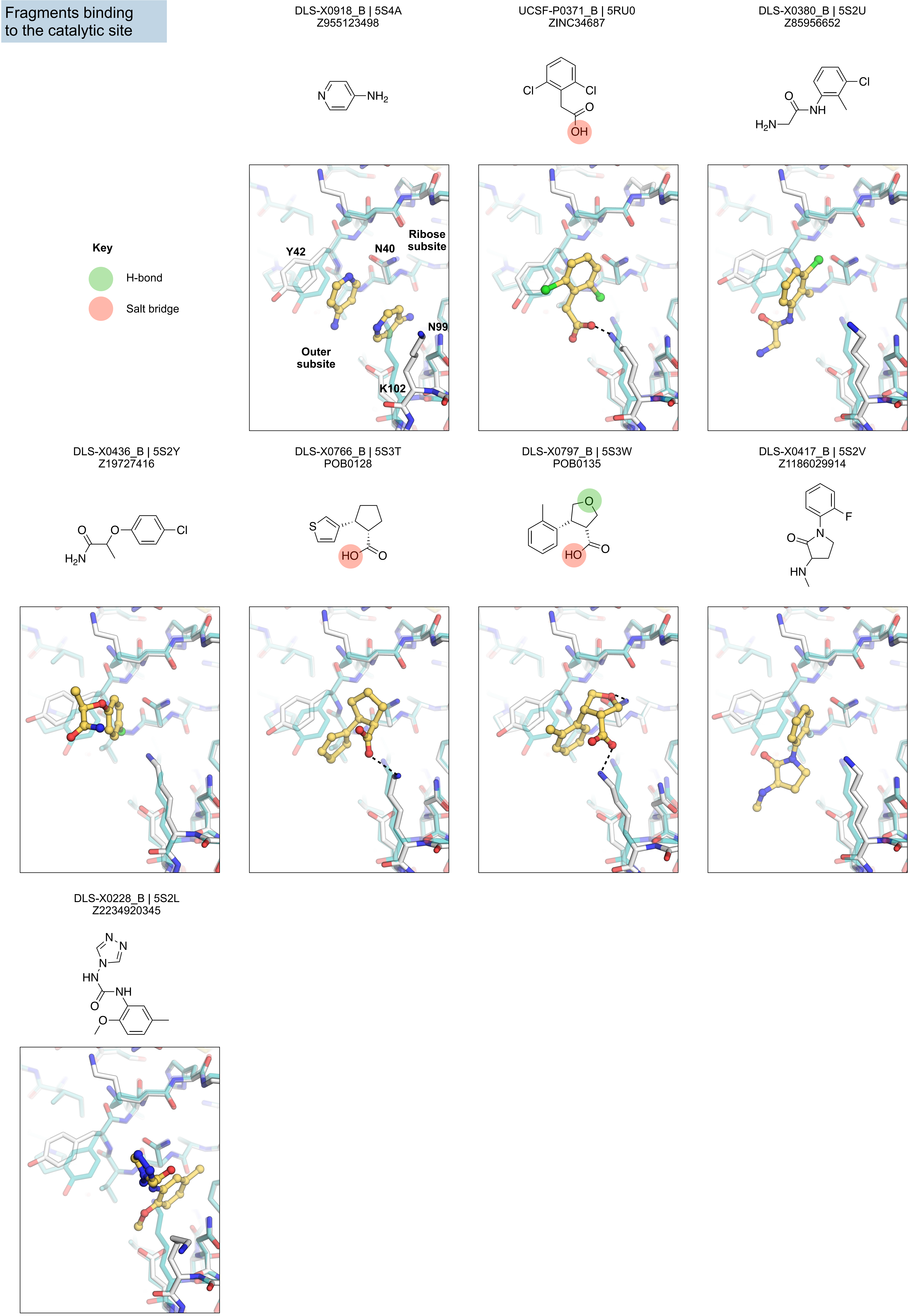

Fragments binding to the catalytic site
DLS-X0918_B | 5S4A
Z955123498
UCSF-P0371_B | 5RU0
ZINC34687
DLS-X0380_B | 5S2U
Z85956652
Key
H-bond
Salt bridge
Y42
N40
Ribose subsite
N99
Outer subsite
K102
DLS-X0436_B | 5S2Y
Z19727416
DLS-X0766_B | 5S3T
POB0128
DLS-X0797_B | 5S3W
POB0135
DLS-X0417_B | 5S2V
Z1186029914
DLS-X0228_B | 5S2L
Z2234920345

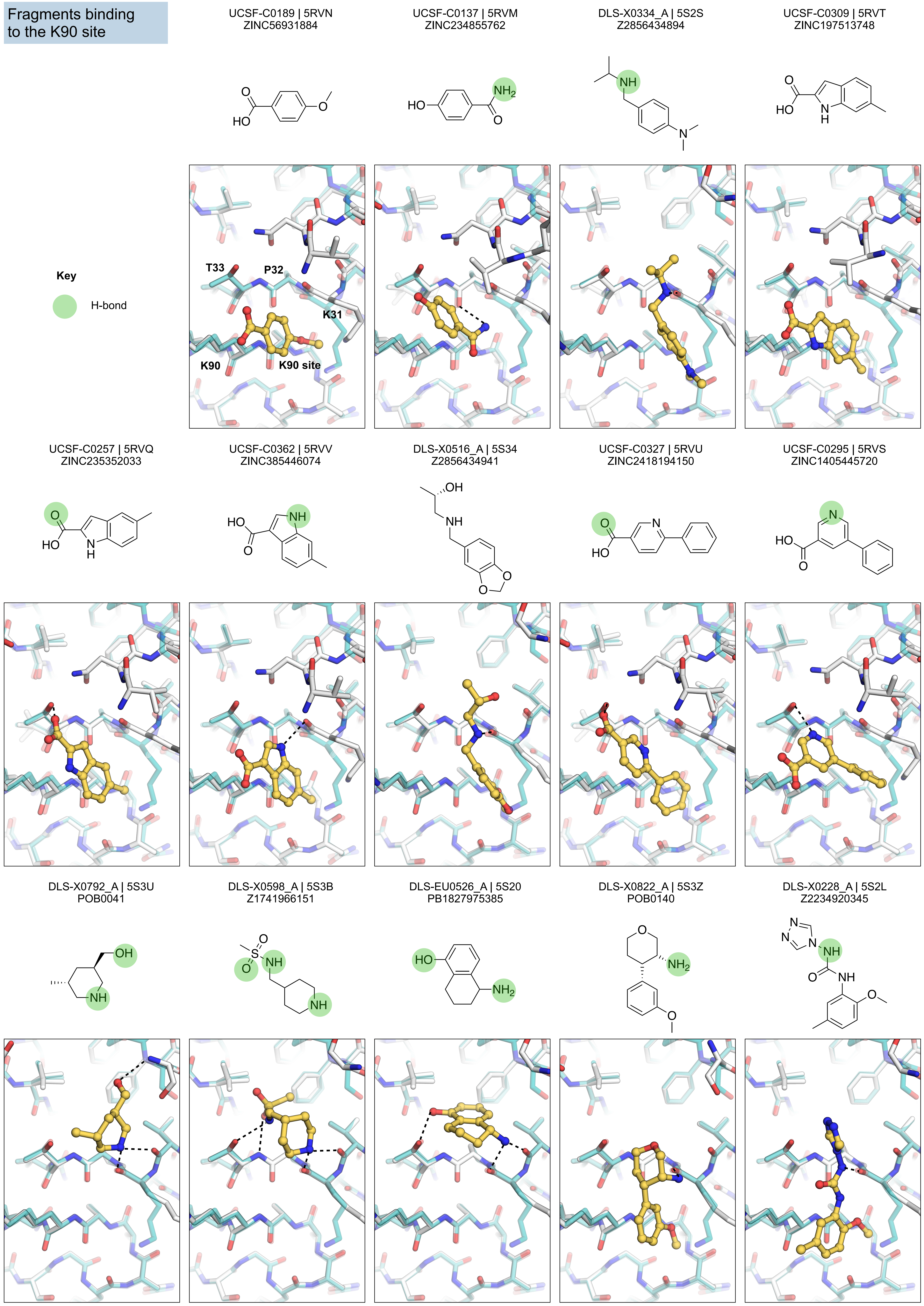

Fragments binding to the K90 site
UCSF-C0189 | 5RVN
ZINC56931884
UCSF-C0137 | 5RVM
ZINC234855762
DLS-X0334_A | 5S2S
Z2856434894
UCSF-C0309 | 5RVT
ZINC197513748
Key
H-bond
T33
P32
K31
K90
K90 site
UCSF-C0257 | 5RVQ
ZINC235352033
UCSF-C0362 | 5RVV
ZINC385446074
DLS-X0516_A | 5S34
Z2856434941
UCSF-C0327 | 5RVU
ZINC2418194150
UCSF-C0295 | 5RVS
ZINC1405445720
DLS-X0792_A | 5S3U
POB0041
DLS-X0598_A | 5S3B
Z1741966151
DLS-EU0526_A | 5S20
PB1827975385
DLS-X0822_A | 5S3Z
POB0140
DLS-X0228_A | 5S2L
Z2234920345

**DSF results for all tested compounds.**

**A)** Changes in $Tm_a$ observed in the presence of ADP-ribose (0-300 μM, positive control), as well as fragments (0-3 mM fragment) observed to bind by crystallography (purple), and those not observed to bind by crystallography (grey). **B)** Corresponding raw DSF data demonstrates the absence of confounding changes in curve shape for all compounds except ZINC157162.

**A** ΔTmₐs for all compounds tested by DSF

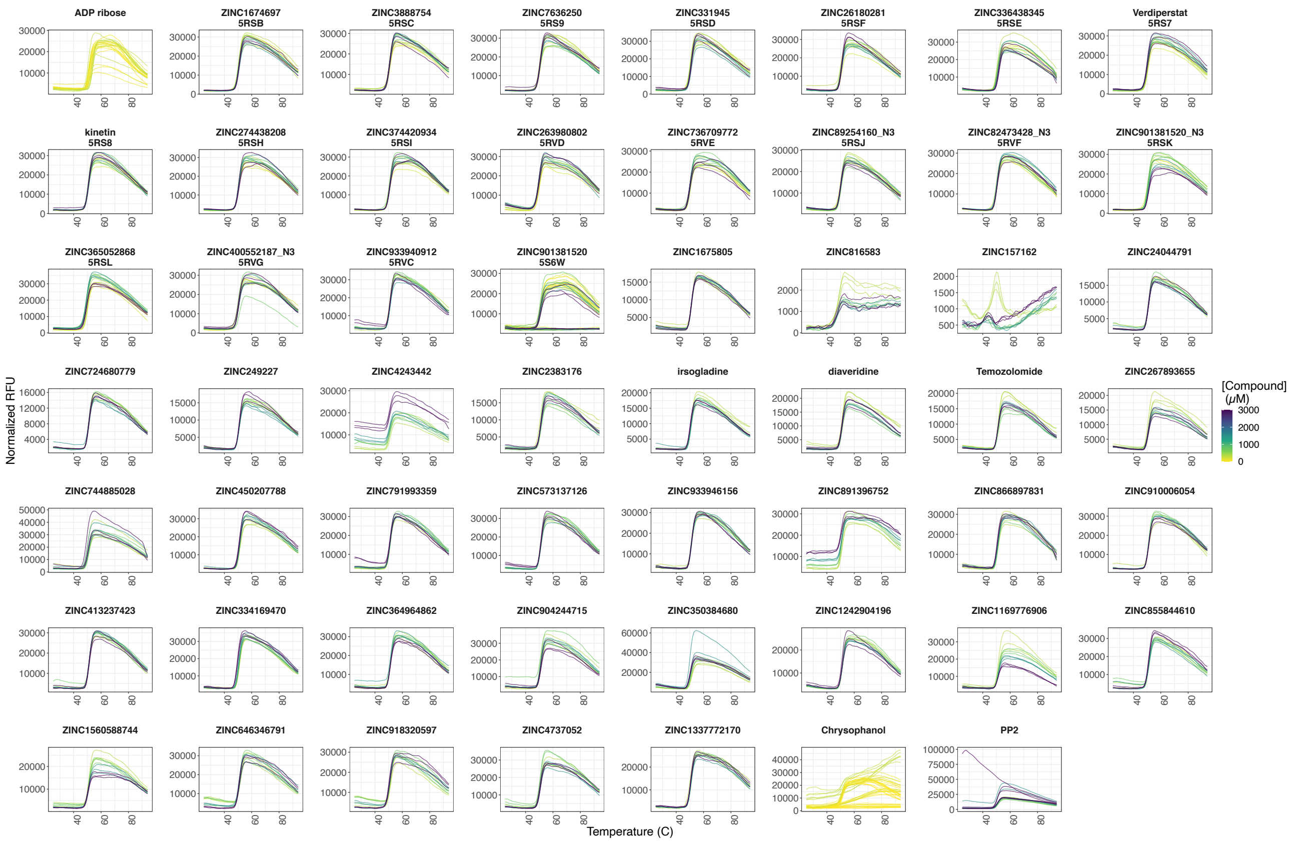


**B** Raw DSF data for all compounds tested by DSF

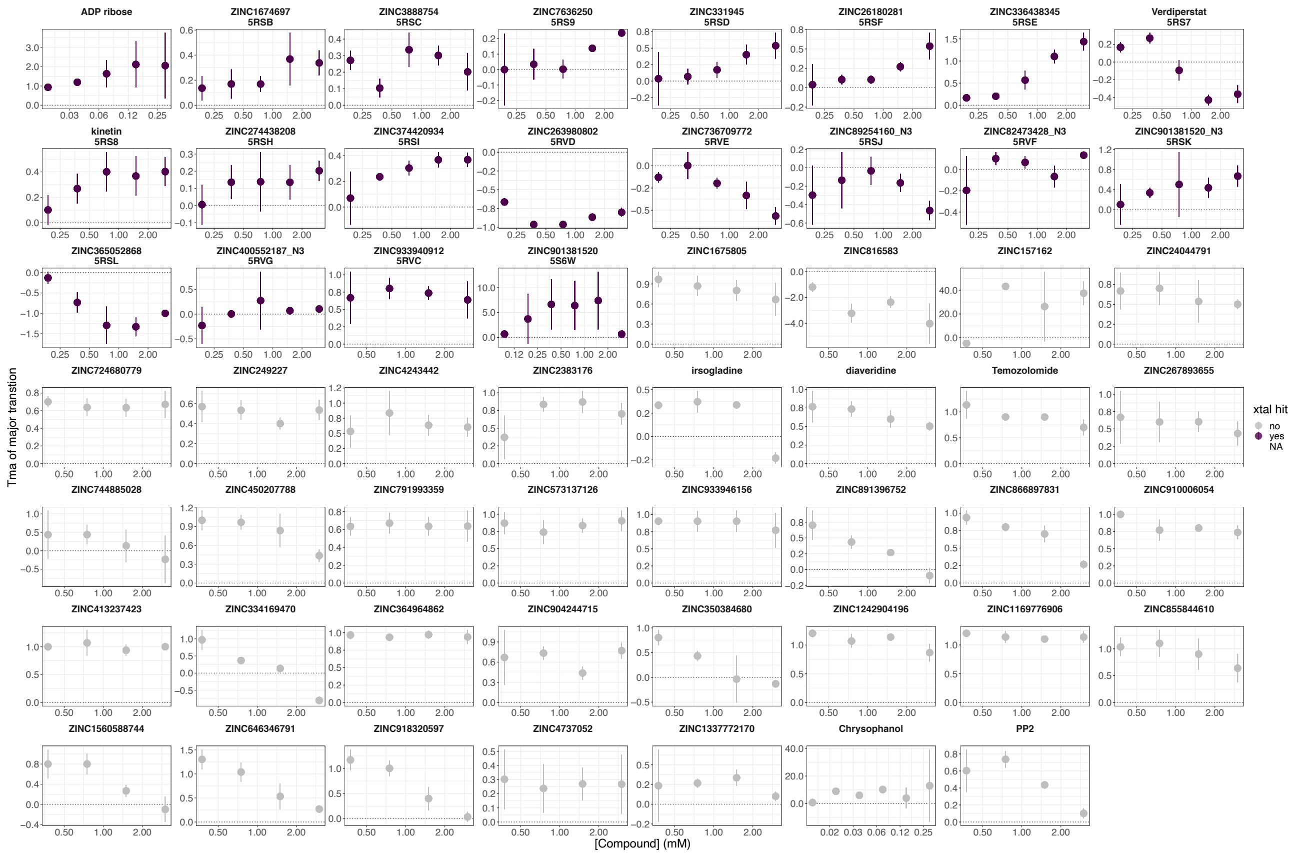

**ITC results for all tested compounds.**
**A)** Raw data for each fragment tested. ZINC26180281 and ZINC331945 were tested twice, therefore raw data from each experiment is plotted. **B)** Binding isotherm for each fragment tested. Integrated heat peaks are represented as blue dots, the non-linear least squares (NLLS) fit using a single-site binding model is represented as a red line. All fragments that could be fit to a NLLS function are shown first. **C)** Residual for each fragment fit to a NLLS function.

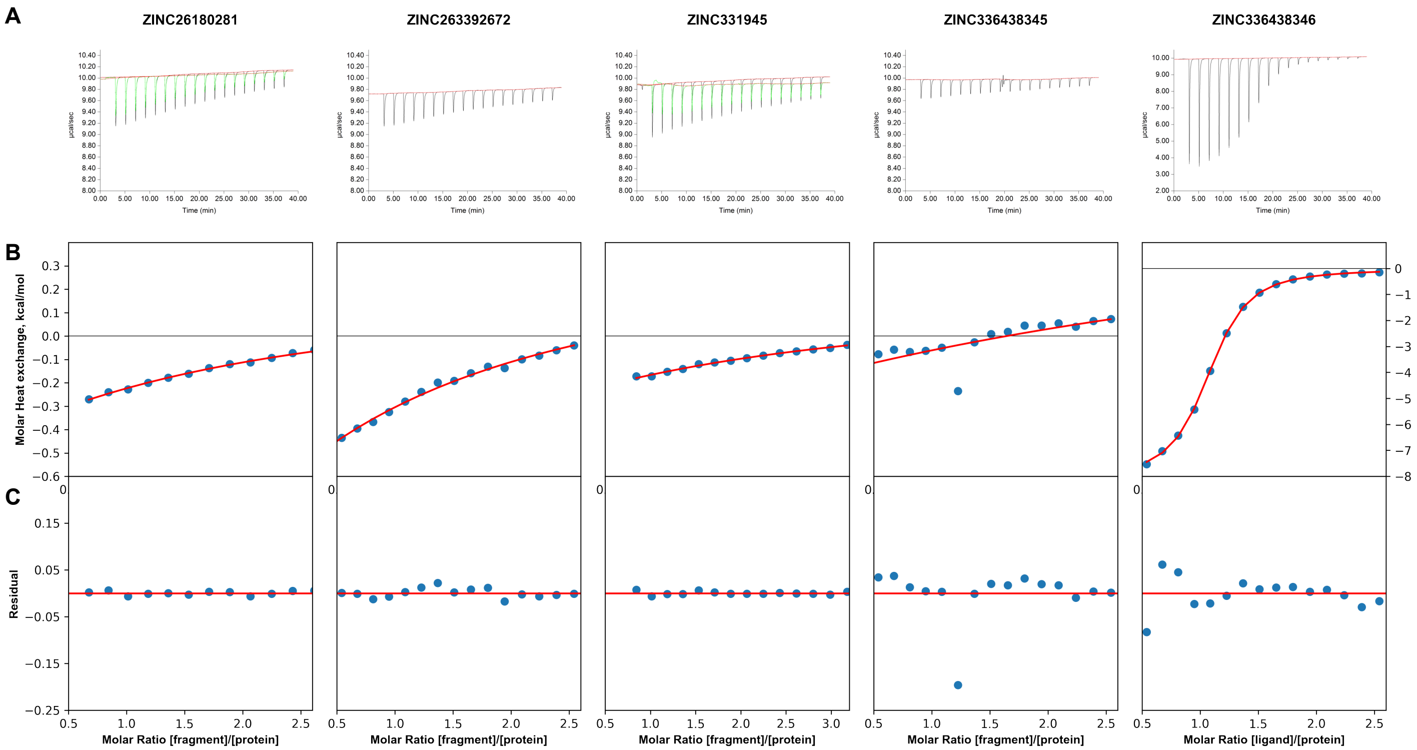


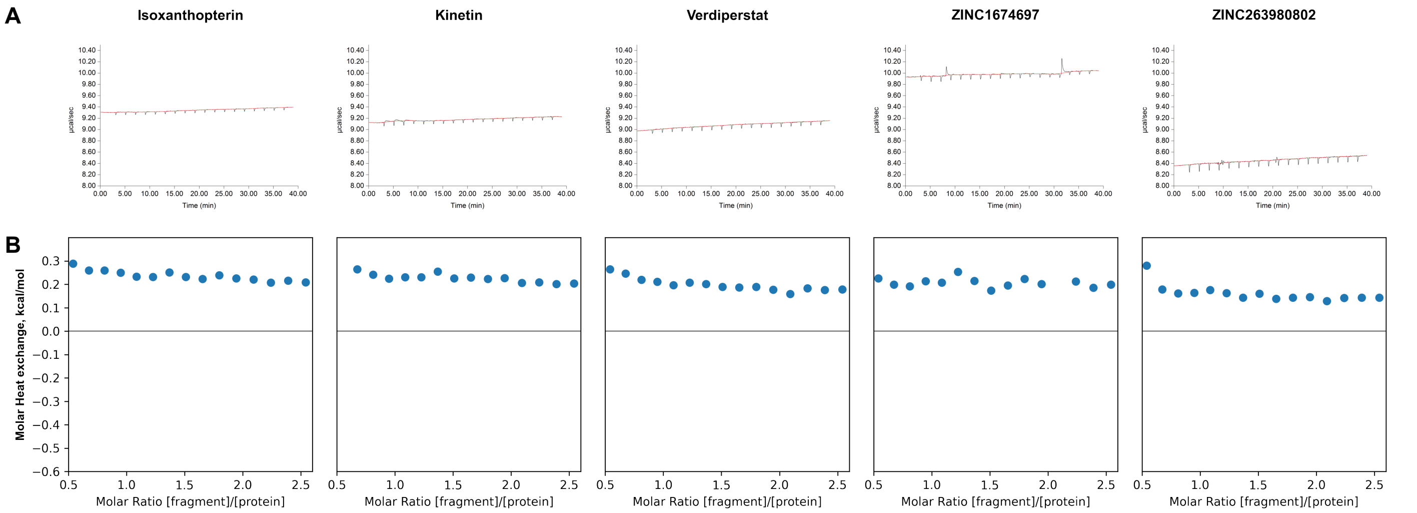

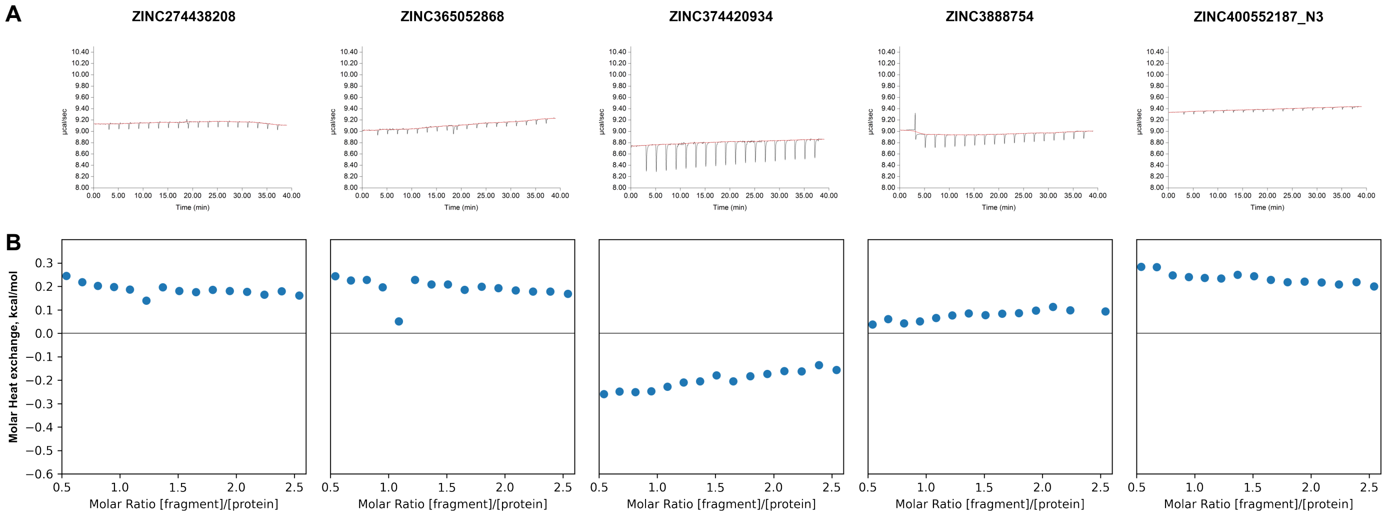
A
ZINC274438208
ZINC365052868
ZINC374420934
ZINC3888754
ZINC400552187_N3
B
Molar Heat exchange, kcal/mol
Molar Ratio [fragment]/[protein]

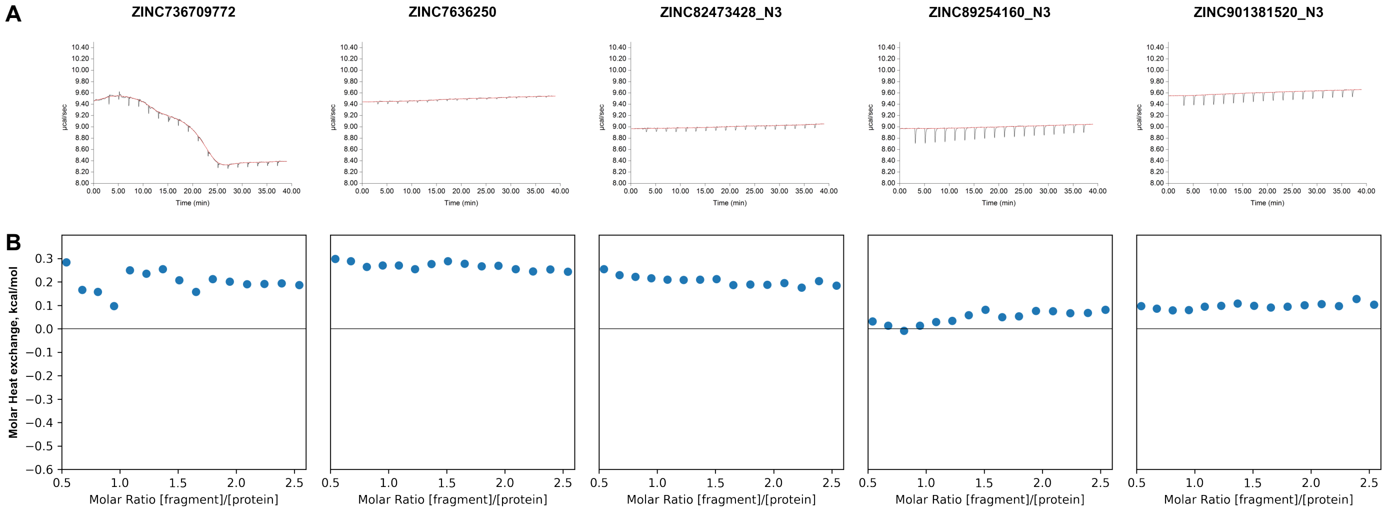
A
ZINC736709772
ZINC7636250
ZINC82473428_N3
ZINC89254160_N3
ZINC901381520_N3
B
Molar Heat exchange, kcal/mol
Molar Ratio [fragment]/[protein]

**Macrodomain peptide displacement results for all tested compounds.**

**A)** Assay results obtained for selected crystallographic hits from the Oxford/XChem screen. **B)** Assay results obtained for docking hits which were crystallographically confirmed. **C)** Assay results obtained for docking hits which could not be crystallographically confirmed. ADP-ribose, used as a control compound in this assay, is shown in the upper left corner of **(A)**. All graphs show normalized compound competition (%) plotted against compound concentration with 1 mM or 2 mM top concentration. Non-linear fits (shown as line) constrained between 0-100% competition were calculated for assay points (mean ± SEM, N = 2). $IC_{50}$ results and competition (%) at top concentration are summarized in Supplementary Table 1.

A

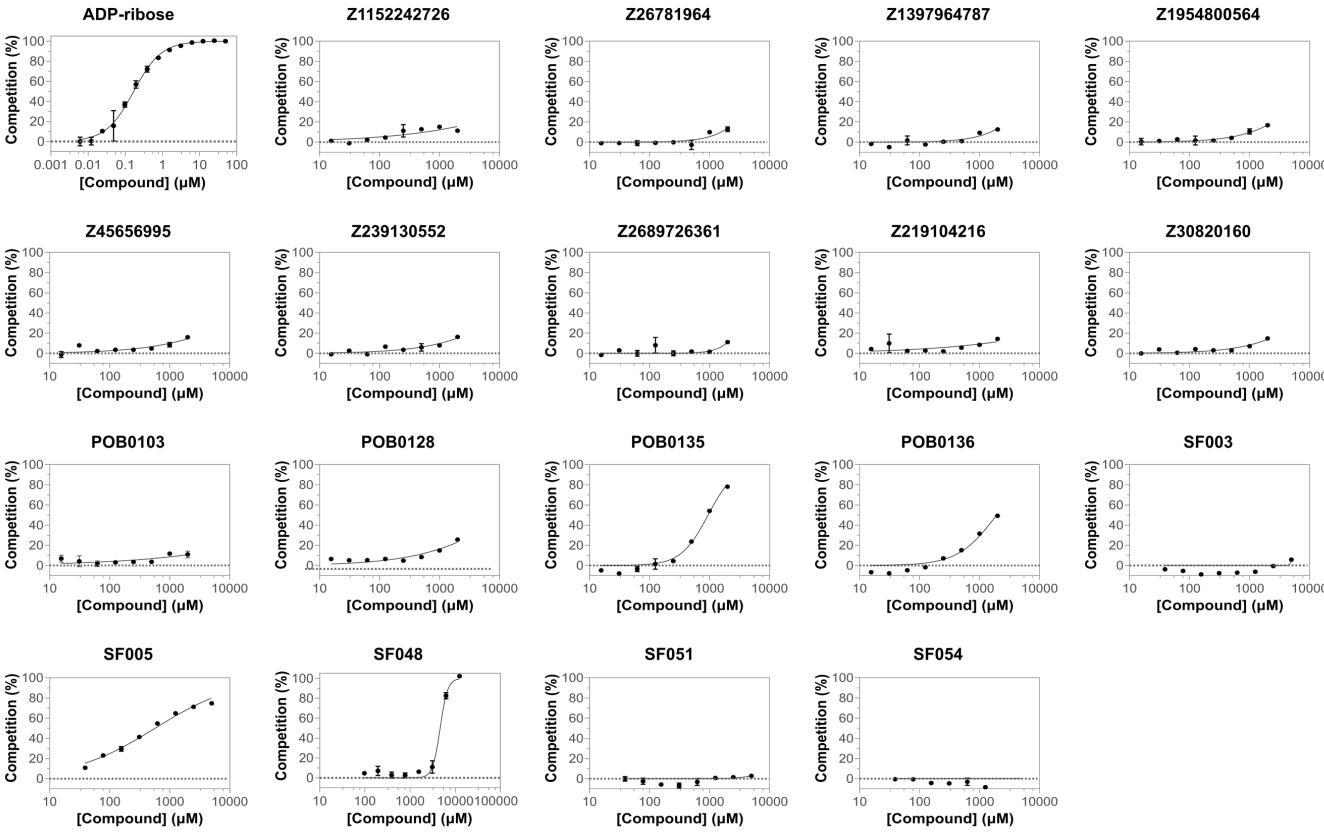
ADP-ribose
Z1152242726
Z26781964
Z1397964787
Z1954800564
Z45656995
Z239130552
Z2689726361
Z219104216
Z30820160
POB0103
POB0128
POB0135
POB0136
SF003
SF005
SF048
SF051
SF054
Competition (%)
[Compound] (μM)

B

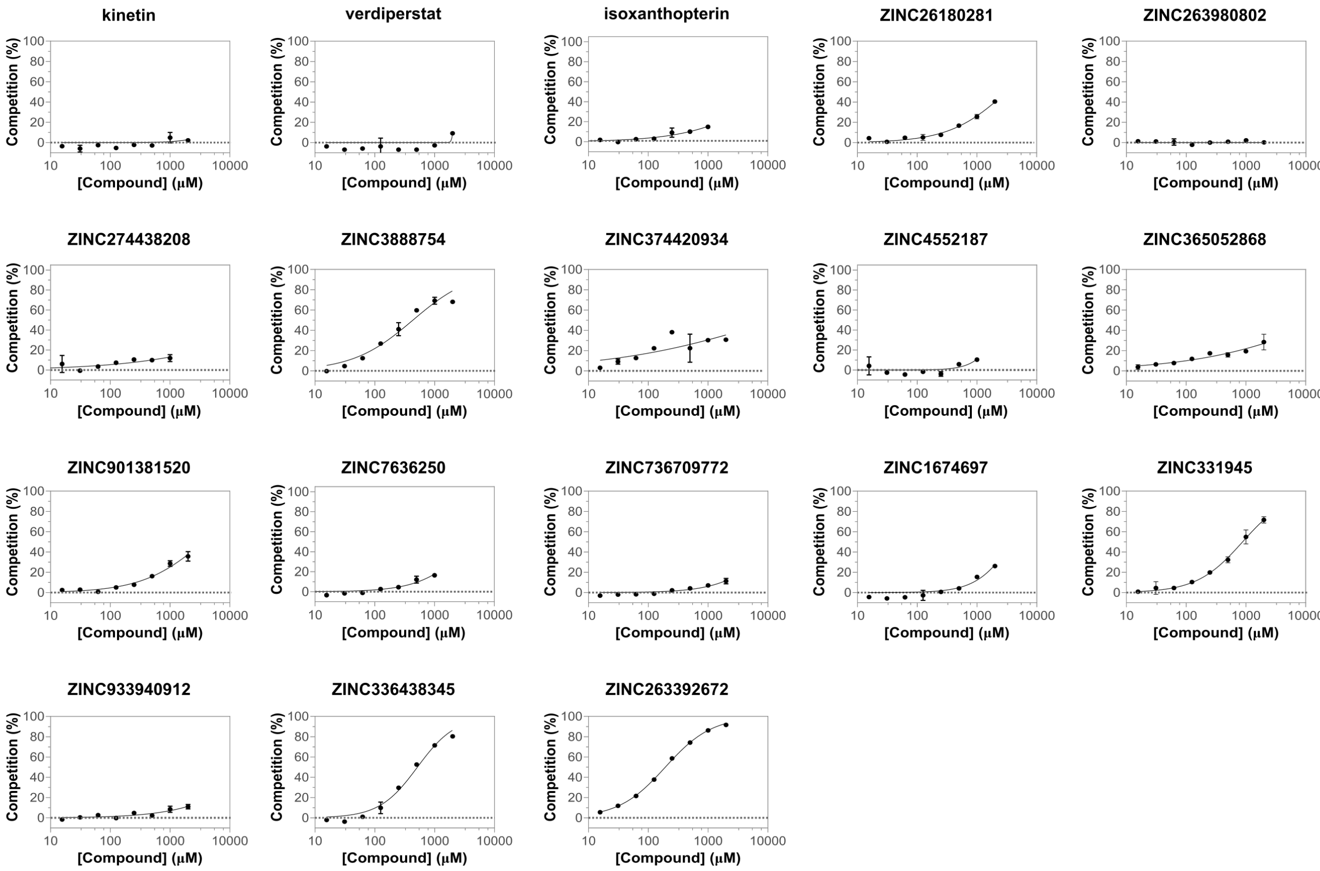
kinetin
verdiperstat
isoxanthopterin
ZINC26180281
ZINC263980802
ZINC274438208
ZINC3888754
ZINC374420934
ZINC4552187
ZINC365052868
ZINC901381520
ZINC7636250
ZINC736709772
ZINC1674697
ZINC331945
ZINC933940912
ZINC336438345
ZINC263392672
Competition (%)
[Compound] (μM)

C

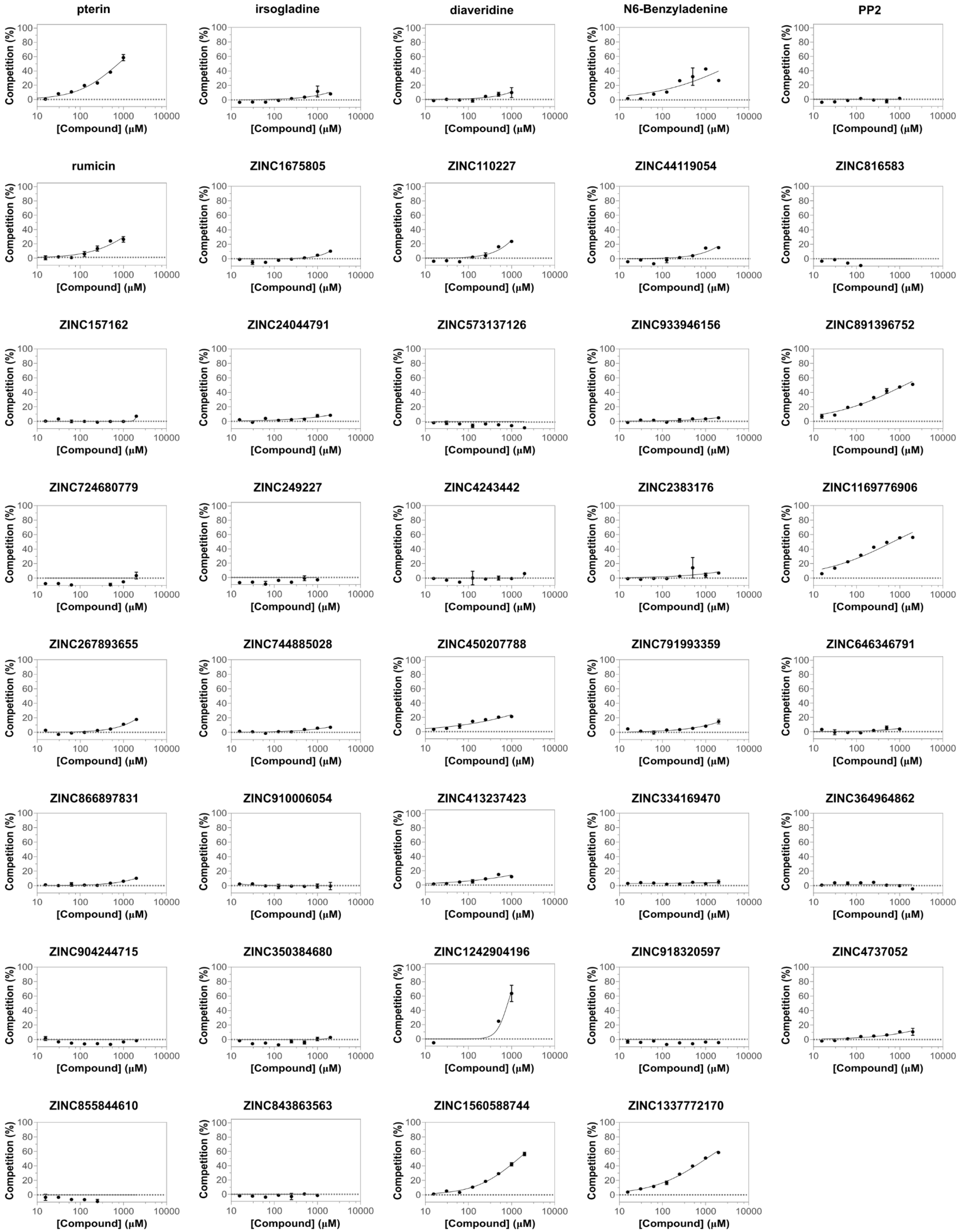
pterin
irsogladine
diaveridine
N6-Benzyladenine
PP2
rumicin
ZINC1675805
ZINC110227
ZINC44119054
ZINC816583
ZINC157162
ZINC24044791
ZINC573137126
ZINC933946156
ZINC891396752
ZINC724680779
ZINC249227
ZINC4243442
ZINC2383176
ZINC1169776906
ZINC267893655
ZINC744885028
ZINC450207788
ZINC791993359
ZINC646346791
ZINC866897831
ZINC910006054
ZINC413237423
ZINC334169470
ZINC364964862
ZINC904244715
ZINC350384680
ZINC1242904196
ZINC918320597
ZINC4737052
ZINC855844610
ZINC843863563
ZINC1560588744
ZINC1337772170
Competition (%)
[Compound] (μM)
